# Probability of outbreaks of forest insects in Europe: a generic model calibrated on six forest insect profiles

**DOI:** 10.1101/2024.09.04.611203

**Authors:** Dorian Collot, Christelle Robinet

**Affiliations:** INRAE, URZF, 45075, Orléans, France

**Keywords:** Modelling, probability, outbreak, Europe, forest, climate, insect, pest, oak processionary moth, *Thaumetopoea processionea*

## Abstract

Insect pests are one of the major threats to forests. Although invasive species cause more and more impacts, native species could also generate temporary very high damages. The population dynamics of insects relies on several factors, going from weather to stand conditions. Due to global change, insects could face conditions they have never encountered before, leading to unusual population outbreaks. Forest managers need to consider these possible emerging pests but predicting insect outbreaks is still very challenging. In this context, we have developed a mathematical model at the crossroad of statistical and mechanistic models to describe the likelihood of outbreaks for a set of 6 insect profiles: bark beetles, longhorn beetles, tortrix moth, other moths, aphids, and Hymenoptera. This model describes the probability of occurrence of an outbreak at a given time and at a given area, based on several conditions. It has been built and parametrized on the most documented orders of European forest pests. Parametrization for these species’ profiles can be used as a baseline to explore the risk of outbreaks for closely related pest species. We provide an illustration of the model application for the oak processionary moth, *Thaumetopoea processionea*, which reach epidemic levels in north-western Europe. This generic outbreak model is particularly performant to point out some years or areas as unlikely for an outbreak, and thus targets correctly factors that inhibit outbreaks. It is still at an exploratory level and should be further improved for an operational use in forest stand surveillance and management.

## 1. Introduction

Forest insects can damage considerably the forest health (Nageleisen et al., 2009; Ramsfield et al., 2016), triggering substantial economic impacts, including not only directly effects of the wood loss, but also indirect effects on the forest sector market (Holmes, 1991). Due to the intensification of international trade, the number of exotic insects newly detected has increased exponentially and recent research has accordingly focused on these invasive species and their impacts (Brockerhoff & Liebhold, 2017). Although native species generally cause less impacts because they have co-evolved with their ecosystem, they could episodically reach high population densities and generate temporary very high damages. Native forest insects are largely distributed all around the world in forested areas and pest management mainly relies on control measures allowing the reduction of the population level under the economically acceptable threshold (Apple & Smith 1976). Due to environmental concerns, chemical treatments are progressively discarded and even forbidden in many countries. Instead, integrated pest management approaches are rising, privileging more environmental-friendly control measures and the identification of the best levers to action, taking into account the forest ecosystem as a whole (Cours et al., 2019). These measures necessitate both efforts for pest surveillance and a good knowledge about its population dynamics, including the drivers of its population growth.

For some of these native forest insects, long series of temporal dynamics are documented either directly with data on population level (e.g., for more than 30 years for the pine processionary moth, *Thaumetopoea pityocampa* ; Li et al., 2015)) or inferred from dendrochronology (e.g. for 1200 years for the larch budmoth, *Zeiraphera diniana*;). Forest insect outbreaks have been studied for a very long time in forestry and a classification arose. Outbreaks’ dynamics have been grouped into three major patterns: gradient pattern (population level is a continuous function of some biotic or abiotic factors), cyclic pattern (the time between peak density is relatively constant), and eruptive pattern (eruptions to very high densities occur at irregular intervals) (Berryman & Stark, 1985). These patterns reflect different underlying mechanisms that are most often not explicitly elucidated.

Although outbreak cycles are documented for a set of well-known forest insects, their drivers are often not known and the underlying mechanism is quite often a black box. For *Zeiraphera diniana*, one of the best examples of cyclic pattern, with very large population fluctuations at about 9 years intervals (Esper et al., 2007), the drivers of its outbreaks are not so clear. The comparison of various models describing the temporal population dynamics highlighted that the parasitoid–budmoth interaction was the main driver of the pest outbreak cycle (Turchin et al., 2003). However, outbreaks of the larch budmoth are shifting in higher elevations, probably because of a better synchronization between the egg hatch and the larch bud flush (Rozenberg et al., 2020). Population outbreaks can thus result from complex interactions.

Another good example of outbreaking species is the oak processionary moth, *Thaumetopoea processionea,* which recolonized a large part of its historical distribution in Europe since the 1970s, and which suddenly reached epidemic levels in north-western Europe where it was considered as almost extinct (Groenen & Meurisse, 2012). Forest management should not ignore this probablility of outbreaks, notably in areas where outbreaks have never been recorded so far or a long time ago, and even where the species is not yet present. Indeed, the oak processionary moth is predicted to expand further in northern Europe by 2050 (Godefroid et al., 2020). This species is a big threat to oaks but also to human and animal populations because the gregarious larvae can release urticating setae in the air (Gottschling & Meyer, 2006). Despite urgent needs of knowledge about this pest dynamics due to noticeable forestry and sanitary impacts, a deep understanding of the outbreak mechanism is still lacking at large scale. Given the difficulty in determining the drivers of outbreaks for relatively well-known species, it appears very challenging to identify drivers and elucidate the outbreak mechanism for less documented case studies.

Furthermore, with climate change, a wide range of effects are observed going from the collapse of population cycles to increased outbreak occurrence, even for a given guild of species within a single region (Haynes et al., 2014). When considering other forest species and other regions, it appears all the more challenging to predict forest insect outbreaks in a changing world. With global changes (including climate change), forest insects face new conditions that could affect their outbreak dynamics, which could turn even more unpredictable.

Pest outbreaks represent major issues for forest managers. Native forest pests being widespread, monitoring requires a lot of time and resources. Determining the conditions triggering these outbreaks is the first step to determine when and where such events could occur. Predicting these outbreaks would contribute to design more efficient ways to manage pests and thus improve forest health protection by prioritizing the species and areas for which a stronger monitoring effort is required.

Models could be useful to make such predictions. Various generic models have been developed to determine the risk of entry (Douma et al., 2017), establishment (Kriticos et al., 2016), spread (Parry et al., 2013; Robinet et al., 2012) and impact of invasive species (Kriticos et al., 2013; Soliman et al., 2012). Contrary to invasive exotic species, native species are generally not subject to specific regulation nor official control, and generic approaches to assess pest risk is not so well developed (Robinet et al., 2020). To describe outbreaks of forest insects, mainly pest-specific models have been developed so far, taking into account some particular life-traits of the species. In addition, previous studies have generally focused on different factors influencing the outbreaks. For instance, the outbreaks of the pine sawfly, *Diprion pini,* are explained by the climatic factors according to Möller et al. (2017) and by topographical factors according to Kosunen et al. (2017). Considering simultaneously these two factors could improve the predictions and help the management of outbreaks.

More generally, two kind of models could describe insect outbreaks: statistical and mechanistic models (Robinet et al., 2020). Statistical models are convenient to describe outbreaks in conditions for which we have already some data but should be applied very carefully to new conditions. Mechanistic models are more robust to extrapolate to new conditions and thus could be helpful to explore the dynamics in changing conditions, but they are in turn more difficult to parametrize as they require a good knowledge about involved processes. For instance, the PHENIPS model (Baier et al., 2007) accurately describes the life-cycle of *Ips typographus* but requires rate of development or rate of successful hibernation, which are unknown for less-studied species. In fact, it is very challenging to identify the whole set of existing outbreak mechanisms. In addition, we should ensure that input variables of the models can be easily measured on the field.

More and more data about population outbreaks and their drivers are available in literature, but linking all the studies and case studies is a difficult task because the data are not standardized throughout published papers on this topic. Therefore, an important step of a generic approach is to gather and standardize all the datasets so that the model can take into account all of them in a single framework and allows to include more and more data as soon as they become available to provide progressively better predictions of the outbreak. In the same way, having the possibility to adapt to local constraint and missing data is also an important feature.

The aim of this study is to develop a generic outbreak model that can provide a first assessment of the outbreak probability for the main forest insect pests in Europe, taking into account every relevant driver. For this purpose, 1) we did a review of literature to extract these drivers, 2) we developed a generic outbreak model providing the probability of an outbreak according to environmental conditions, and 3) we applied this model to a wide range of insect species to document several insect profiles. Since data may not be available to correctly parameterize this outbreak model for any species, the associated species profile can be considered to have a first assessment of the outbreak likelihood of the study species in some conditions. This model was designed to assess the probability of outbreak even in new conditions and for species that are not well known. A big step forward toward the genericity (and also a big amount of work behind this study) was the standardization of the current knowledge as the drivers of insect outbreaks identified in literature were generally different from one case study to another, even if they were closely related (e.g., related to temperature). Due to this standardization, the model may not consider the best one for a given well-known species and may not perform so well than a pest-specific model already developed. However, it provides a rough outbreak probability: (i) for species whose outbreaks are less or not at all documented, and (ii) for sites where outbreaks have not been recorded so far. This generic outbreak model, at the crossroad of statistical and mechanistic models, was designed so that it can easily be refined without re-performing the model calibration as more and more data and information are available for a given species. A shiny app has been developed to provide a friendly user interface for the model. This application provides a visualisation of both: (i) input parameters and functions, and (ii) model outputs. To illustrate this approach, we assess the probability of outbreaks of the oak processionary moth (OPM) in north- western France.

## 2. Materials and Methods

### 2.1. Systematic review of outbreak drivers

The drivers have been determined based on a literature review. This analysis has been performed using Web of Science, and the request “((pest OR pathogen OR insect) NOT non-native NOT exotic NOT invasive) AND (outbreak OR spread) AND forest AND (Europe OR Austria OR Belgium OR Britain OR Croatia OR Czech OR Denmark OR Finland OR France OR Germany OR Greece OR Hungary OR Ireland OR Italy OR Netherlands OR Norway OR Poland OR Portugal OR Romania OR Scotland OR Serbia OR Spain OR Sweden OR Switzerland OR Turkey OR United Kingdom)”. No conditions on the date or language have been requested. Due to a high number of untargeted articles, we then disregarded the articles that described: (i) non-native species, (ii) animal pest, as ticks and mosquitoes, and (iii) the consequence of an outbreak. Therefore, we kept only 22% of articles to extract the outbreak drivers.

### 2.2 Model description

#### 2.2.1 General mathematical structure

The model aims at providing a probability of outbreak given various variables (Fig. 1). For each variable, we estimate a single outbreak probability and then we should merge all these individual probabilities into a single one. The model was thus inspired by data fusion methods used in imagery (Bloch & Maitre, 1994) in which several probabilities are calculated, each one based on one observation (here an environmental variables). These probabilities are then combined into a single one, based on fuzzy set methods (as described below).

**Figure 1:**
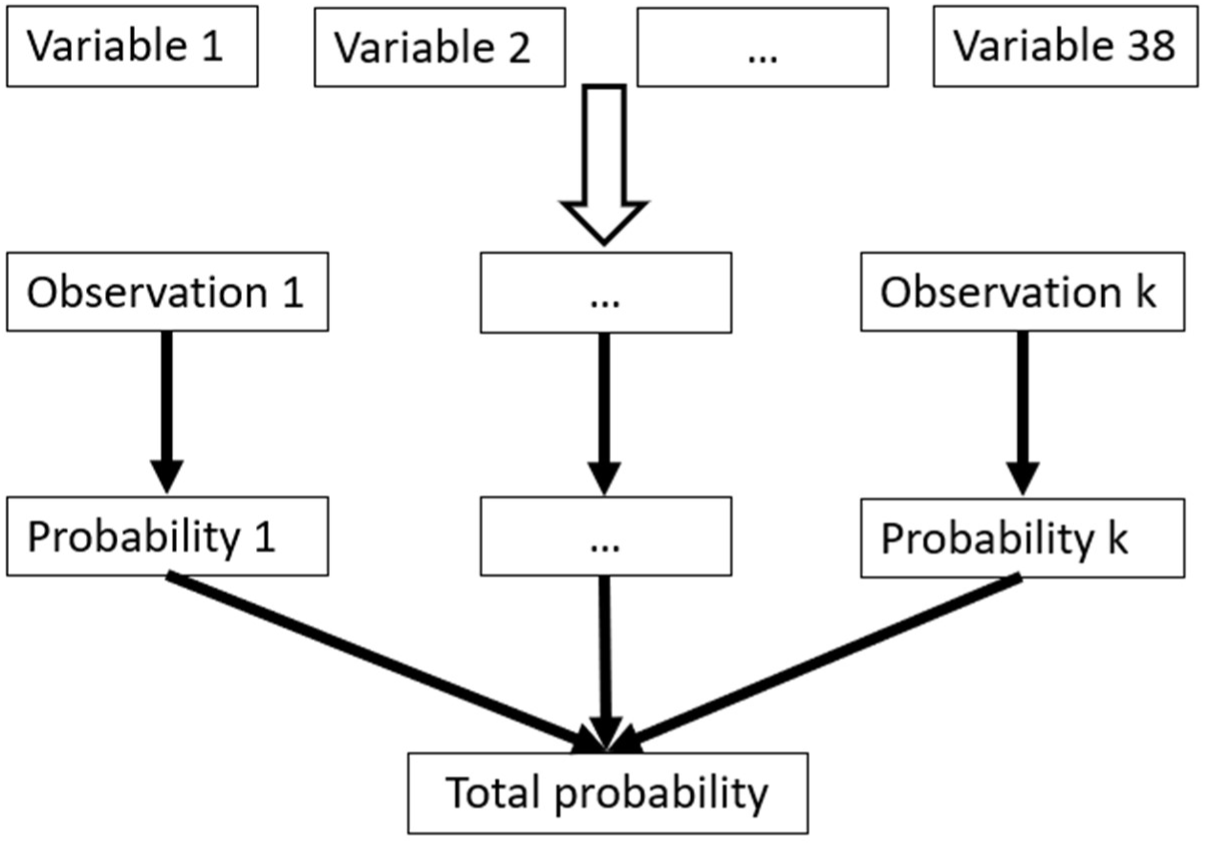
Flow chart of the model to estimate the probability of outbreak. The first step is to choose the relevant variables for the species among the 38 variables used by the model. Each of the *k* relevant variables will be associated to an observation. The second step is the computation of the probability for each observation. The third step is to compute the geometric mean of the *k* previous probabilities to obtain the total probability of outbreak.

The occurrence of an outbreak is described by a Bernoulli law of mean ***p*** *=* **P(outbreak|(*x1,…, xn*))** which depends on different variables ***xi***, as follow:

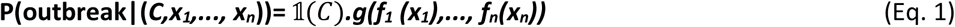

The functions ***fi*** correspond to the responses to each variable ***xi*** individually (i.e., ***fi*** is the probability of outbreak when considering only one environmental variable) and the function ***g*** combines these responses into a single probability. We considered the indicator function 𝟙 to account for the impossibility of an outbreak in some conditions (e.g., absence of hosts):

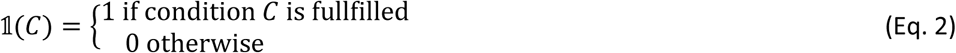

Since this indicator function provides an indication about the possibility for outbreaks (Boolean variable) and not a probability, it is not part of the variables of the function ***g***. This model has been implemented in R (Collot & Robinet, 2021; R Core Team, 2018) with the Shiny package (Chang et al., 2019) to provide a user-friendly interface. This interface allows an easy way to select the species profiles and parameters, and gives a spatial map of the model simulations as well as a summary table for the model outputs (SM1).

#### 2.2.2 Definition of the function ***g***

The function ***g*** defined in (Eq. 1) should fulfil three conditions.

1. It should provide a value between 0 and 1.
2. The probability considering multiple variables cannot be higher or lower than the probability considering isolated event (“cautious” behavior ; Bloch, 1996):

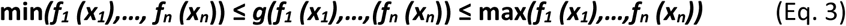

3) If a variable leads to the impossibility of the outbreak when considered alone (e.g., ***fi(xi)*=0**), then the outbreak is impossible, disregarding all the drivers (***g(f1 (x1),…,fn (xn))*=0**).

These criteria ensure that removing or adding a driver will not drastically affect the model output. Indeed, the number of drivers does not affect this probability, as it would do if we simply multiplied or added the functions ***fi***. Any mean function satisfies the two first constraints, however arithmetic, harmonic and quadric means do not verify the third constraint. We thus chose ***g*** as the geometric mean of the functions ***fi***.

#### 2.2.3 Definition of the functions ***fi***

Since the functions ***fi*** describe a probability, like the function ***g***, it is should also provide a value between 0 and 1. These functions ***fi*** are defined by five parameters:

- ***min*** and ***max***: corresponding to the boundaries of the interval of definition of the function.
- ***pmax***: the highest probability reached for the given variable (≤1). The functions then go from

**[*min,max*]** to **[0, *pmax*]**.

- ***a*** and ***b*** determine the shape of the functions. Their effect depends on the type of function which is considered.

We considered four types of function (Fig. 2):

- “Convex” curves are monotonic and could be actually either convex or concave, ***a*** determines the minimal values, and ***b*** defines the convexity of the curves (0 being a straight line).
- “Logistic” curves are also monotonic, but with an inflection point (i.e. they are convex and concave), ***a*** defines the position of the inflection point, and ***b*** the slope at the inflection point.
- “Gaussian” curves are non-monotonic curves, ***a*** defines the mean, and ***b*** defines the variance.
- “Beta” curves are non-monotonic curves as well, but can be asymmetric, contrary to “Gaussian” curves. ***a*** defined the “asymmetry” of the curve, and ***b*** defines the shape. The function reaches a maximum if ***b*** is positive, and a minimum if ***b*** is negative.

**Figure 2:**
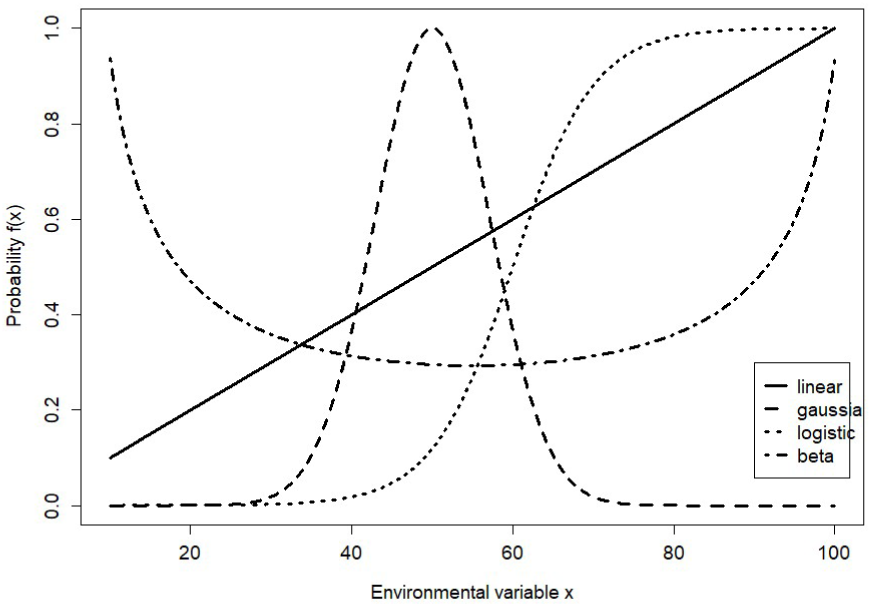
Four types of function ***fi*** considered to provide probabilities of outbreak.

“Beta” and “Convex” functions are defined only on the interval [***min***,***max***] but can be extended to ℝ by setting: ***f(xi)*=*f(min)*** if ***xi*<*min***, and ***f(xi)*=*f(max)*** if ***xi*>*max***.

### 2.3 Datasets and parameters estimation

#### 2.3.1 Datasets

As we collected data from various scientific papers, the environmental data were very heterogeneous and were related to different database sources. To develop a generic model, we had to homogenize the input variables. Climatic data used in this model come from the dataset CRU-TS 4.03 (Harris et al., 2014) downscaled with WorldClim 2.1 (Fick & Hijmans, 2017). This dataset contains the monthly sum of precipitations and average monthly minimal and maximal temperatures from 1960 to 2018 throughout Europe. The mean temperature was computed using the average monthly minimal and maximal temperature and then averaged by trimester. The soil types are those retrieved from the FAO/UNESCO dataset (FAO, 1992). The altitude was based on the SRTM (Shuttle Radar Topography Mission) data, found on the WorldClim website. The slope (in %) and its direction were computed using the altitude. Forest covering data has been retrieved from the EFI (European Forest Institute) (Brus et al., 2012). The age of the trees, the deadwood volume, and the previous outbreaks are not given by such generic databases and were therefore not available for several case studies.

Outbreak data have been retrieved from the database of the Department of Forest Health (DSF) of the French Ministry of Agriculture and Food. This data contains the reported outbreak in the French Forest from 2007 to 2018. These data have been used for the inference of parameters values. The test of the model has been done using the literature data from Havašová et al. (2017), Buse et al. (2007), Lima et al. (2008), Lesniak (1976) and Möller et al. (2017).

#### 2.3.2 Estimation of the parameters

Approximate Bayesian Computation (ABC) methods (Beaumont et al., 2002) were used to infer the values of the parameters from data. In these methods, simulations are made according to a prior distribution of the parameters. Simulations are then compared to the data via a distance (or likelihood when it is possible). The posterior distribution is then defined based on the simulation with the lowest distance.

Prior distribution of the parameters has been set based on the literature. Models previously published for each insect’s order indicate the variables to consider and a first estimate of the parameters. Biological data have also been considered, even if they have not been identified by a modelling approach.

We used the ABC-SMC method (Toni et al., 2009). This algorithm is iterative. At each loop, 100.000 simulations are made, using parameters randomly chosen according to the prior distribution. For each simulation, the likelihood is computed. The simulations with the highest likelihood (>90% of the maximal likelihood) are kept to define the prior of the next loop. The algorithm is the following :

1. 100.000 parameters values are randomly chosen according to the prior distribution.
2. For each one of this parameters’ set, the probability of an outbreak occurring is computed using the data.
3. The likelihood of the data is then computed.
4. The parameters sets are then sorted according to their likelihood and the 90% with the lower likelihood are discarded.
5. The 1.000 remaining parameters sets are used to defined the new prior distribution.

These five steps are repeated 10 times to estimate the model parameters.

#### 2.3.3 Species profile

For many species, data and good knowledge are lacking, and it is very challenging to identify the potential drivers for outbreaks. We therefore defined drivers of outbreaks for different orders or families, supposing that two species of the same profile will have similar probabilities of outbreaking in the same environmental conditions. A total of 147 European pests have been considered (SM3), and the most frequent orders among those outbreaking forest pests have been selected to represent the main pest profiles. This list of species has been established from Branco et al. (2015), and then completed by species encountered in literature during the parametrization phase. This is therefore not an exhaustive review, it should nevertheless give a good indication of which species are the most studied.

#### 2.3.4 Model validation

For each profile, 90% of the data have been used for the estimation of the parameters. The remaining data have been used to check the accuracy of the prediction, by computing the Area Under the ROC Curve (AUC) (Fawcett, 2006).

For each year, the model has been used to predict the probability that an outbreak occurs. In order to compare the data and the prediction, a threshold has to be chosen. If the probability is higher than the threshold, the model predicts that an outbreak should occur. The sensitivity and the specificity were computed for different thresholds. The ROC curves obtained were then used to compute the AUC. The AUC is comprised between 0.5 and 1 and a value of 0.5 means that the predictions are not better than random while a value of 1 indicates a very good prediction.

### 2.4 Application to a case study: the oak processionary moth

To illustrate this approach, we applied this generic outbreak model on the oak processionary moth (OPM), *Thaumetopoea processionea,* in the northeast quarter of France (48.0-51.2°N; 2.0-8.2°E). This area has been discretized in 19,200 points, equally spaced, representing field sites. For each site, input datasets (e.g., environmental data) have been retrieved from 1970 to 2018. This study area has been under pressure of the OPM for years, and it is expected that this species will cause more and more damages in the future.

First, we explored the spatial and temporal dynamics of the OPM outbreaks in the past (1970-2018). The change in the dynamics has been studied by comparing the minimum, maximum and average probabilities predicted for each decade. Then, we compared the model outputs with observation data from the French Forest Health Department (DSF – French Ministry of Agriculture) recorded from 2007 to 2018 in this study area. Both spatial and temporal fitting success of the model outputs have been explored. Indeed, we compared the number of observations in a given area to the maximal expected probability of outbreak for a given area on the period. For that purpose, the study area has been divided in 640 squares, in order to have enough data per square. The predictions and observations on a given year have been compared using the Pearson correlation coefficient.

Secondly, we performed simulations considering various climate change scenarios for the future. Since the OPM corresponds to the “moth” profile, the outbreak model accounts for three weather variables (see Results): the aridity index in spring and fall of the previous year and the rainfall in winter of the current year. The future scenarios have been built using the 2010s decade and adding +1°C, +1.5°C or +2°C to the spring and fall temperature to the 2010s dataset.

## 3. Results

### 3.1 Drivers extracted from the systematic review and standardization process

A total of 129 articles have been considered in the systematic review (SM2). From these articles, the main drivers can be sorted in 5 categories: (1) variables related to weather (mean temperature, sum of precipitation, aridity, and lowest temperature), (2) to the stand, (3) to biotic conditions, (4) to other variables (pollution and management), and (5) to spatial and temporal patterns of the outbreaks.

Hereafter, we describe in more details the results of the systematic review, including the percentage of articles pointing out each category of drivers. For each category, variables representative of each driver have been selected. The selected variables should be easy to measure and should be relevant for many pests in order to allow comparison between them. A total of 38 variables have been selected (Table 1).

**Table 1:** List of the 38 variables used as potential drivers.

| Drivers | Description |
| --- | --- |
| T_w | Mean temperature from January to March of the current year |
| P_w | Sum of rainfall from January to March of the current year |
| I_w | Aridity index from January to March of the current year |
| T_s | Mean temperature from April to June of the current year |
| P_s | Sum of rainfall from April to June of the current year |
| I_s | Aridity index from April to June of the current year |
| T_su | Mean temperature from July to September of the current year |
| P_su | Sum of rainfall from July to September of the current year |
| I_su | Aridity index from July to September of the current year |
| T_f | Mean temperature from October to December of the current year |
| P_f | Sum of rainfall from October to December of the current year |
| I_f | Aridity index from October to December of the current year |
| T_w_prec | Mean temperature from January to March of the previous year |
| P_w_prec | Sum of rainfall from January to March of the previous year |
| I_w_prec | Aridity index from January to March of the previous year |
| T_s_prec | Mean temperature from April to June of the previous year |
| P_s_prec | Sum of rainfall from April to June of the previous year |
| I_s_prec | Aridity index from April to June of the previous year |
| T_su_prec | Mean temperature from July to September of the previous year |
| P_su_prec | Sum of rainfall from July to September of the previous year |
| I_su_prec | Aridity index from July to September of the previous year |
| T_f_prec | Mean temperature from October to December of the previous year |
| P_f_prec | Sum of rainfall from October to December of the previous year |
| I_f_prec | Aridity index from October to December of the previous year |
| Min | minimal temperature current year |
| min_prec | minimal temperature previous year |
| Age | mean age of the host |
| Soil | rich/poor soil |
| direction | direction of the slope (South=0, East=West=0.5, North=1) |
| Slope | % of the slope |
| deadwood.volume | volume of deadwood |
| elevation | mean elevation |
| conifer | presence of conifer (0=absente, 1=present) |
| broadleaf | presence of broadleaf tree (0=absente, 1=present) |
| host density | density of the host<br>(conifer or braodleaf tree) (0 to 100%) |
| adjacent | outbreak on an adjacent stand the previous year (0= no outbreak,<br>1=occurrence of an outbreak) |
| previous | outbreak on the stand the previous year (0= no outbreak, 1=occurrence of<br>an outbreak) |
| Cycle | time since the last outbreak |
| latitude |  |
| longitude |  |

#### 3.1.1 Variables related to the weather

Weather was clearly the main driver identified, as it was found in 50% of the articles. The weather variables describing this category of driver are very heterogeneous. For instance, the weather variable goes from the North Atlantic Oscillation (NAO) (Lima et al., 2008) that describes the weather of the whole year, to very precise weather conditions a few days around after the budding of the host (Möller et al., 2017). To standardize this kind of dataset and build a generic model, we considered 4 types of variables related to the weather: the mean temperature on each trimester (***T***), the sum of precipitations on each trimester (***P***), the de Martonne aridity index (***IdM***), and the lowest absolute annual temperature for a given year (***LT***). De Martonne index (Pellicone et al., 2019) is calculated as 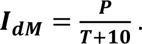. It is well-defined if the mean temperature is above -10°C. This index is a proxy for other variables harder to compute, such as evapotranspiration, available water content, or moisture. Temperature, precipitation, and aridity index were measured for each trimester, corresponding roughly to each season (winter: January to March, spring: April to June, summer: July to September, and fall: October to December), leading to a total of 12 variables, plus the minimal temperature (Pellicone et al., 2019). Since the development of insects can start the year before an outbreak occurs, these variables were considered for two consecutive years, leading to a total of 26 variables.

Windfalls and storms were important drivers of outbreaks (7% of the articles), but they are described by the amount of deadwood on the stand and were thus considered as part of the variables to related to the stand (see hereafter).

#### 3.1.2 Variables related to the stand

The stand situation was found to be a driver of an outbreak in 19% of the articles of the review. These drivers do not change from year to year (or slowly). As a consequence, they cannot actually trigger an outbreak, but instead they can affect its location. These drivers have to be considered to understand why a stand is attacked while another is not, even if the climatic condition is similar on both sites. The stand situation was described by its topography and information about the tree population, such as the age of the tree, or the composition of the mix. The stand situation in the model was thus described by 6 variables: the altitude, the slope, and its direction, the soil type, the age of the trees, and the deadwood volume. Soil was sorted into 3 types: coarse, fine and medium. These three categories are a proxy for the richness of the soil. The age is used as a proxy for other stand factors, such as the diameters of the tree.

#### 3.1.3 Variables related to biotic conditions

Biotic relations are important drivers of the outbreak, found in 21% of the articles. The population densities of the pest can be affected by the population density of its host and its predators/parasites. Predators and parasites generally arise when the population level of their host is high, thus inducing mortality which makes host density decrease (Matek & Pernek, 2018). Predators and parasites are therefore a driver for the end of the outbreak. In addition, it is relatively difficult to obtain data about predators or parasites to feed a generic model. Consequently, the presence or density of predators/parasites was not considered in our generic outbreak model. We considered only three variables related to the host and the forest cover: the conifer presence, the broadleaves presence, and the density of the host (broadleaves or conifer).

#### 3.1.4 Other variables

Forest management and pollution were also found to be important drivers of insect outbreaks in the literature. However, management (11% of the articles) can be related to some drivers already identified. For instance, a clear-cut is a modification of the absence/presence of the host, and could therefore be modelled via a host-related variable. Pollution was not so frequently reported (2% of the articles) and it was not commonly measured in published studies. It was thus not possible to obtain an estimation of its impact on different pest profiles. Consequently, we discarded pollution and management variables from our generic outbreak model.

#### 3.1.5 Outbreak spatial and temporal patterns

Variables that described the past outbreaks have not been found explicitly as drivers, but have been added to the model. The occurrence of an outbreak on a given site or an adjacent site the previous year(s) can enhance or inhibit the occurrence of a new outbreak. Three variables described this phenomenon: the occurrence of an outbreak the previous year, the time since the last outbreak, and the occurrence of an outbreak on an adjacent site.

### 3.2 Species profile

The four most represented pests were *Coleoptera* (37%), *Lepidoptera* (31%), *Hemiptera* (16%), and *Hymenoptera* (8%)*. Diptera* (4%) were also important pests, but we did not find enough information to make a complete profile. *Lepidoptera* were mainly represented by moths.

Since *Coleoptera* and *Lepidoptera* are very large orders, we refined these profiles and considered families instead of orders. *Coleoptera* were divided in two groups: bark beetles and ambrosia beetles (16%) and longhorn beetles (4%). *Lepidoptera* were also divided in two groups: tortrix moths (7%) and other moths (24 %). Therefore, in total, there are 6 insect profiles. The parameters of these profiles are given in Table 2. They have been implemented directly into the apps by default.

**Table 2:** Baseline parameters for each of the 6 insect profiles.

| profile | driver | Type of function | a | b | P_max | [min;max] |
| --- | --- | --- | --- | --- | --- | --- |
| Bark beetles and ambrosia beetles | T_w | gaussian | 0 | 0.2 | 1 | [0;10] |
|  | direction | gaussian | -0.5 | 0.05 | 1 | [0;3.14] |
|  | I_su | linear | 0.2 | 0 | 1 | [-5;10] |
|  | Deadwood.volume | logistic | -0.2 | 3 | 1 | [0;2] |
| Longhorn beetles | T_s | gaussian | 0.05 | 0.05 | 1 | [5;15] |
|  | Adjacent | linear | 0.5 | 0 | 0.9 | [0;1] |
|  | age | logistic | 0 | 2.3 | 1 | [10;50] |
|  | broadleaf | indicator | 0.5 | 2 | 0 | [0;1] |
| Tortrix Moth | T_su | gaussian | 0.2 | 0.2 | 1 | [15;25] |
|  | Age | Logistic | -0.2 | 1.6 | 1 | [0;80] |
|  | P_w | Logistic | 0 | 0.2 | 1 | [50;300] |
|  | Soil | Indicator | 0.5 | 2 | 0.2 | [0;1] |
| Other moths | I_s_prec | gaussian | -0.03 | -0.98 | 1 | [5;15] |
|  | I_f_prec | gaussian | -0.29 | 0.14 | 0.98 | [5;15] |
|  | P_w | beta | -0.84 | 0.31 | 0.99 | [44;301] |
|  | Host density | logistic | -0.35 | 1.6 | 1 | [0;1] |
| Aphids | I_su_prec | gaussian | 0.37 | 1 | 1 | [-5;10] |
|  | Previous | Linear | 0.1 | 0 | 1 | [0;1] |
|  | T_s_prec | Gaussian | -0.028 | 0.94 | 1 | [0;15] |
|  | T_s | Gaussian | 0.34 | 0.88 | 1 | [-5;10] |
|  | I_su | gaussian | 0.28 | 0.99 | 1 | [-5;10] |
| Hymenoptera | T_su | Gaussian | -0.17 | -0.62 | 1 | [15;25] |
|  | Host density | Indicator | 50 | 200 | 1 | [0;1] |
|  | P_s | Logistic | 0 | 0.2 | 1 | [90;200] |
|  | P_s_prec | Logistic | 0 | 0.2 | 1 | [90;200] |
|  | T_su_prec | gaussian | -0.17 | -0.58 | 1 | [15;25] |
|  | I_su_prec | Gaussian | -0.35 | -0.46 | 1 | [5;15] |
|  | I_su | gaussian | -0.34 | -0.98 | 1 | [5;15] |

### 3.3 Validation

The validation has been made on the data in the literature for each insect profile. We therefore obtained the different graphs (fig 3). The AUC was between 0.6 and 0.7, which is quite low but relatively good for a generic model. When comparing occurrences of outbreaks with predicted probabilities, we found that the model was globally able to discriminate years unlikely to have an outbreak (SM4). When the probability was low, no outbreak effectively occurred, but high probability of outbreak has been predicted for some years without outbreak.

**Figure 3:**
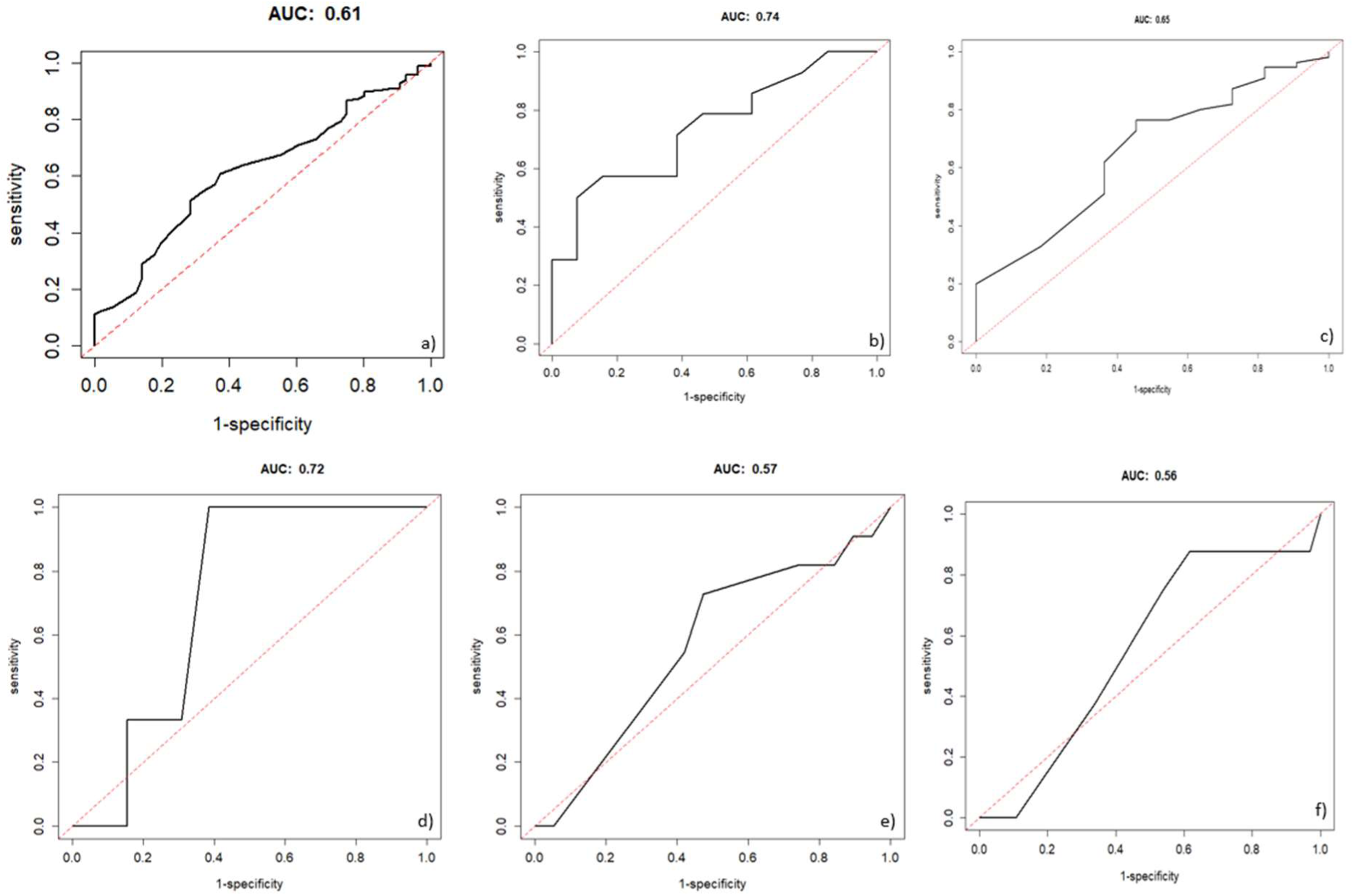
Roc curves for the different profiles: a) Bark beetles, b) Other moth, c) Longhorn beetles, d) Hymenoptera, e) Hemiptera and f) Tortrix moth.

### 3.4 Application on the Oak Processionary Moth in the North-East of France

When predicting the OPM outbreaks in the past (from 1970’s to 2010’s), a large and persistent spatial variability was observed (Fig. 4). The temporal change in the probability of outbreak did not show a clear pattern (prediction for every year from 1970 to 2018 is given in SM5). Surprisingly, probability of outbreaks was high in the 1970’s. Then, they were low in the 1980’s, and then they slightly increased. When we compared the model outputs to observations during the whole period 2007-2018, the number of observations of the OPM was slightly correlated to the maximal probability of outbreak (correlation coefficient of 0.12) and, more importantly, both globally matched on the map (Fig. 5). When comparing predictions to observations each year, we observed a mismatch that could be corrected when shifting the predictions to one year before the observations (Fig. 6).

**Figure 4:**
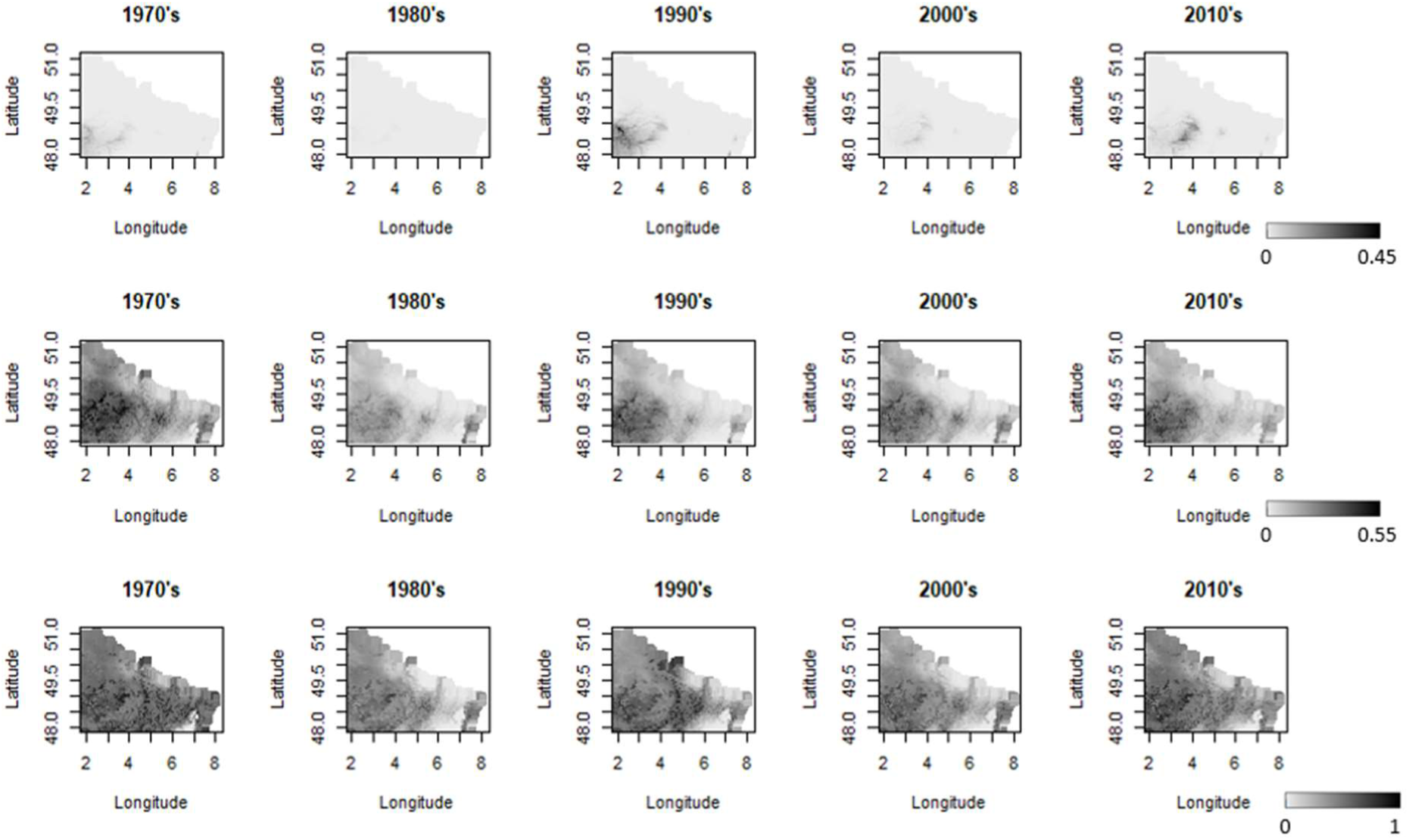
Average minimal, mean and maximal probability of outbreak (respectively at the top, the centre and the bottom) of the oak processionary moth for each decade, following the outputs of the generic outbreak model using the “Other moth” profile.

**Figure 5:**
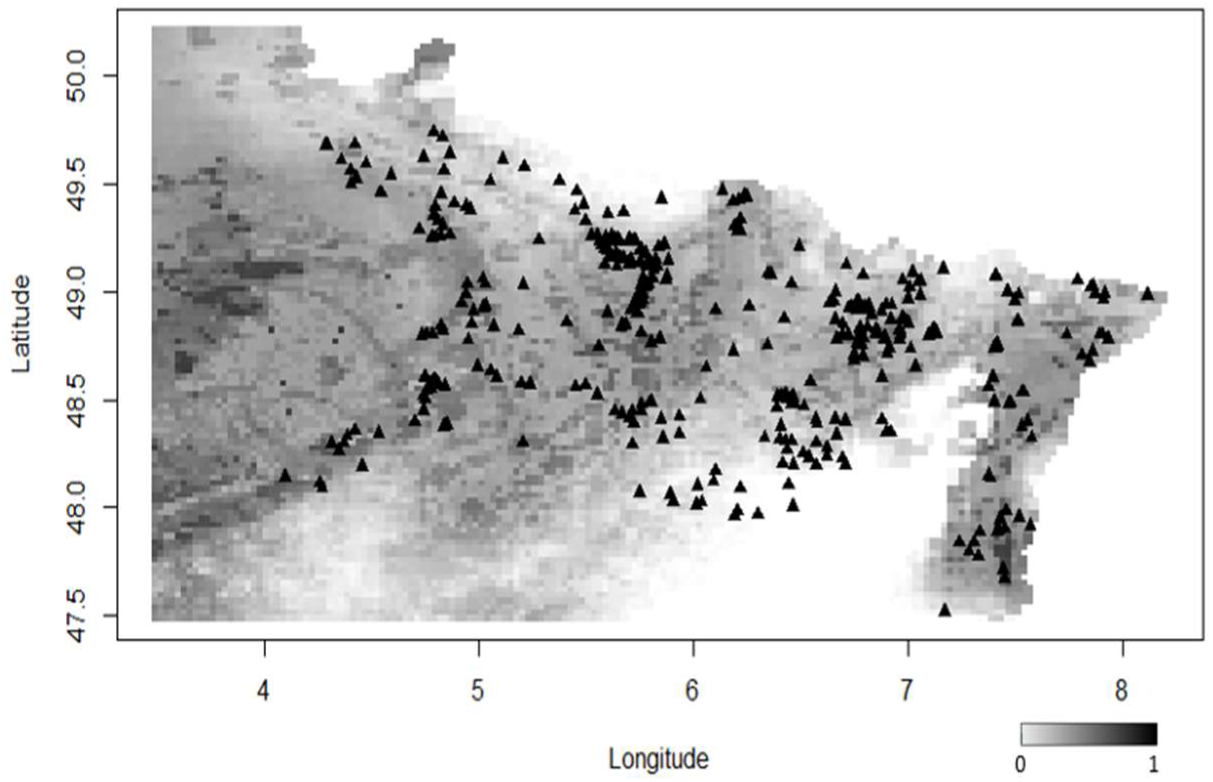
Area of outbreak for the oak processionary moth for the period 2007-2018. The shades of grey represent the maximal probability that an outbreak occur on this period following the generic outbreak model outputs, and the black triangles correspond to sites where nests of OPM have been observed during this period.

**Figure 6:**
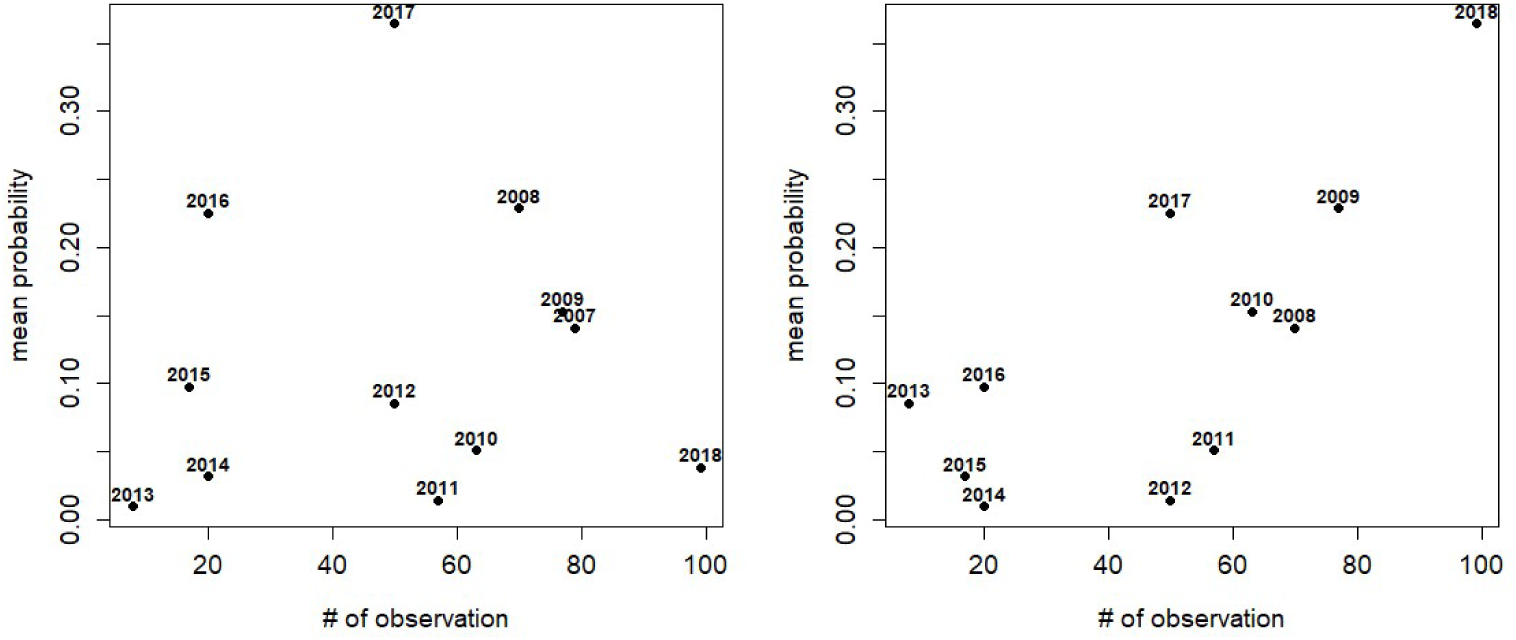
Correlation between the average probability that an outbreak occurs in a given year, and the number OPM observations that year. On the left, the probability is correlated with the number of OPM observations the same year. On the right, the probability is computed with a one-year shift. The date corresponds to the date of the observation.

Lastly, under further climate warming, the spatial pattern of OPM outbreaks should roughly remain the same, but probabilities of outbreak will progressively increase in most of the study area (Fig. 7).

**Figure 7:**
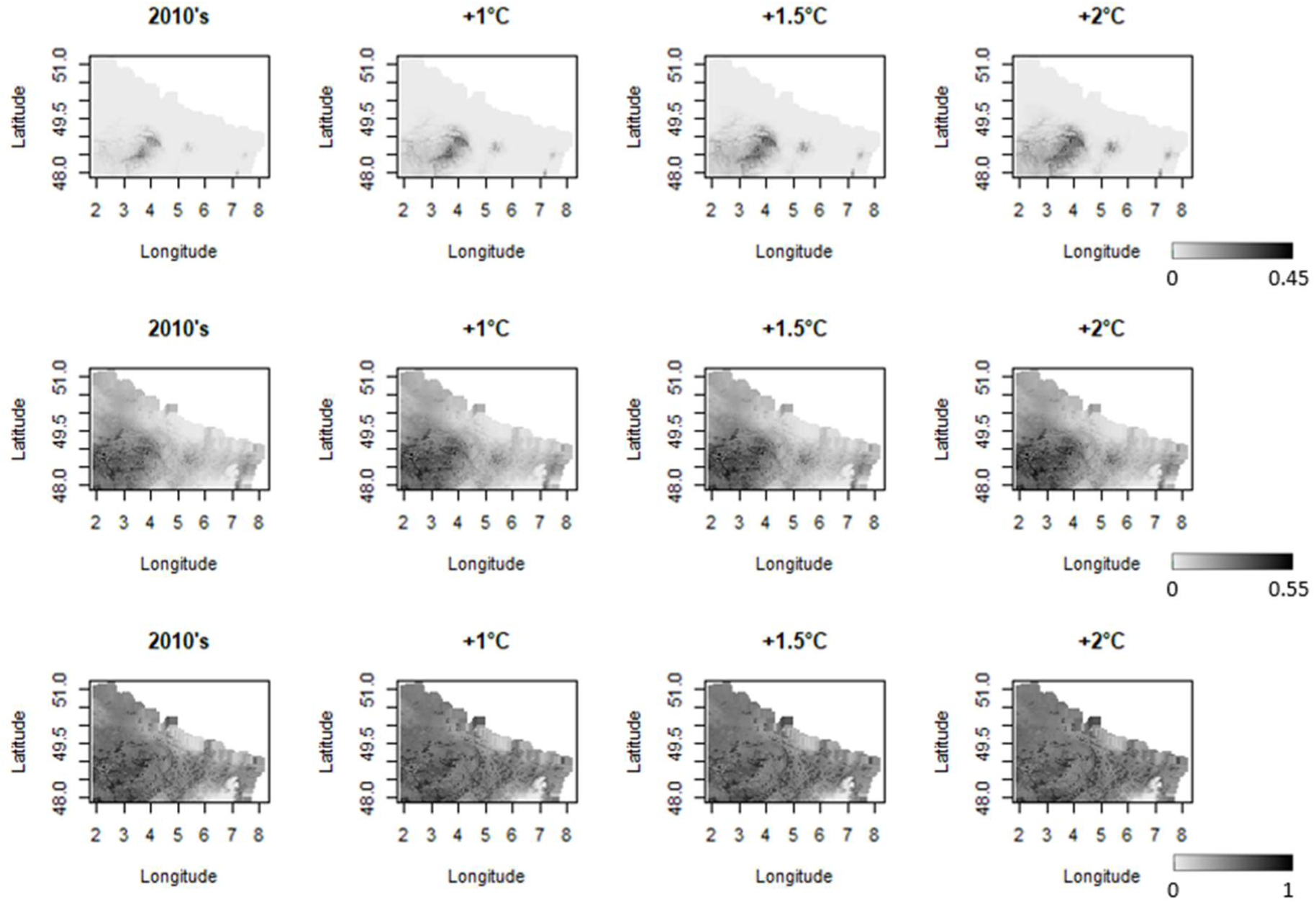
Average minimal, mean and maximal probability of outbreak (respectively at the top, the centre and the bottom) of the oak processionary moth under future climate warming scenarios (+1, 1.5 and 2°C compare to the temperatures in the 2010 decade).

## 4. Discussion

In this study, we have built a generic model to assess the probability of outbreak of some forest pests. We parametrized this model for six pest profiles to provide a baseline library of parameterization. The choice for these profiles was driven by a balance between enough data (as it is not possible to have good datasets at finer species group levels) and ability of the model to provide good outputs (at least a good indication of the likelihood of a species to outbreak or not). Insect profiles have been chosen to represent the main pest orders. For the largest ones (*Coleoptera* and *Lepidoptera*), profiles have been parametrised for families rather than orders. Some pests have been discarded, as *Curculionidae* or *Buprestidae* for *Coleoptera*. These insects, such as *Agrilus* spp., are important forest pests, as some of them are invasive (e.g. *Agrilus planipennis* (Valenta et al., 2017)) or emerging (e.g. *Agrilus biguttatus* (Moraal & Hilszczanski, 2000)) but not enough information has been found on these families to parameterize the model. Similarly, *Hemiptera* was only represented by aphids, due to a lack of data on scale insects.

The model we have developed is, to our knowledge, the very first generic outbreak model and it is thus a very important step forward in the understanding of forest insect outbreaks. It can relatively correctly predict the outbreak occurrence. More precisely, the model is particularly performant to point out some years or some areas as unlikely for an outbreak to occur for a given insect profile. This suggests that the model better accounts for factors that inhibit outbreaks than factors promoting them. The second explanation is that some outbreaks may have occurred but other factors not explicitly included in the model (e.g., forest management or biotic interactions) prevented them. However, this generic model is still an exploratory tool and cannot be used for forest management before some improvements. This generic model does not aim to provide a better assessment of the outbreak probability for well-known and documented species (such as *Ips typographus*), but instead it provides a baseline to explore potential outbreaks for less documented species.

The application of the generic outbreak model to the OPM gave relatively good results. First, the model predicted an increase of the outbreak probability of the species from 1980’s to 2010’s, in agreement with the general trend observed in Europe (Grenen & Meurisse 2012). Second, the comparison of observations and predictions matched quite well in space (the outbreak mainly occurs in high-probability area), and in time (although with a 1-year shift). Considering that no data of the OPM have been used to parametrize the “moth” profile, it highlights the good performance of the model in case no data is available for further parametrization. There are, however, a few deviations of the model simulations. First, the model predicts high probability of outbreaks in the 1970s. To our knowledge, these high population levels have not been observed in this decade. As discussed previously, the model tends to predict more outbreaks that observed (or in other words, there are more “false positives” than “false negatives”). Nevertheless, we know that the OPM has recolonized Belgium, the Netherlands and Germany since the 1970s (Groenen & Meurisse, 2012) and, thus, it was perhaps not present in the north east quarter of France in the 1970s although the conditions were likely favourable for outbreaks. Secondly, there is a one-year shift between predictions and observations: the model predicts outbreak one year earlier. This could be attributed to a latency in the life-cycle of the OPM, which could lay more eggs than usual during years with good conditions for outbreaking, but the population actually reaches high population size on the following year (compared to other species in the same profile). Furthermore, we compared the probability of outbreaks from our model to observations of OPM presence on the field, which does not necessarily mean that OPM outbreaks occurred because we have no information about the OPM population level (and thus, outbreaks) at these spatial and temporal scales. Following the various sources of uncertainties, and the expected (moderate) increase of the outbreak probabilities with further climate warming, it could be interesting to generate more accurate predictions. For instance, input variables could be refined. The aridity index from May to July, as well as severe droughts in summer, are known to drive the outbreak of this species (Csóka et al., 2018) but were not considered in the insect profile (see Moth profile, Table 2). Considering these refined variables would contribute to draw more pest-specific and accurate predictions for pest management in the coming years.

Hereafter, we review three main directions to improve the outputs of this generic model, notably in terms of pest-specific results and thus pest management.

### 1. Refining or adding input variables to increase the model performance for well-documented insect species

More relevant variables (or more precise data) can be added to improve the model performance on a given insect species, as proposed for the OPM. The variables have been selected to be easy to measure and cover many species. As a consequence, these variables are average or proxy of other variables. For instance, the evapotranspiration is often relevant (e.g. (Ismail et al., 2010)), but is difficult to measure, we therefore approximated it by an aridity index. In the same way, insects are sensitive to temperature at given periods of their life-cycle (Faccoli, 2009). Therefore, considering the temperature on a full trimester may not be the most relevant. Considering a more appropriate variable rather than a general proxy could improve the model performance.

### 2. Considering species life-traits to refine the generic outbreak model

Another way to increase the model performance across insect species is to explore a potential link between their outbreak pattern and their phenotype (e.g., species life-traits), rather than considering that two species of the same profile (namely, orders or families) have similar probabilities of outbreaking in the same environmental conditions. Considering life-traits in this model will strengthen the mechanistic causal-relationship of the outbreaks and will certainly increase the model robustness. We started to explore this approach but the lack of data and precise description of these relationships for many species prevented us to make this link so far. Collecting data about outbreaks and species phenotypes is necessary to build an appropriate dataset in the future and make this improvement.

### 3. Predicting population size

The definition of a probability threshold above which we consider that an outbreak is expected can affect the prediction success. Instead, the model could predict the population size rather than the occurrence of an outbreak. In this case, the probability law has to be changed by a binomial law and there is no need to define a threshold for an outbreak. However, predicting population size leads to a more difficult estimation of the parameters as the population size is not so often measured and reported. Moreover, different species of the same orders can reach outbreak levels at various population size. It would therefore be difficult to describe the probability of outbreak based on the same population abundance scale and threshold in the model.

Even if improvements are possible, this model already provides a good baseline to explore outbreak probability for various insect profiles in Europe. Knowing which years an outbreak is unlikely to occur is an important information for forest managers. In this case, monitoring efforts could be reduced to save time and resources. In turn, these efforts could be reallocated to somewhere else or to some other insects that have a higher probability to make an outbreak and cause tree damage. In the future, this kind of model could support surveillance prioritization, which is actually an important step in forest stand management.

## Acknowledgements

We thank François-Xavier Saintonge and Fabien Caroulle (DSF – French Ministry of Agriculture) for providing datasets, Alain Roques, Jérôme Rousselet, and Mathieu Laparie (INRAE URZF – France) for fruitful discussions, Bob Douma (Wageningen University – Netherlands) and Andrea Battisti (Padova University – Italy) for their insightful comments.

## Author contributions

DC and CR designed the study, DC made the systematic review and developed the model, CR supervised the work, DC and CR wrote the paper.

## SUPPLEMENTARY MATERIAL

SM1: Print screens of Shiny Ap interface developed for this generic outbreak model.

SM2: List of publications found and studied in the systematic review, later used to extract the drivers

SM3: List of the 147 European forest insects considered to build profiles SM4: Comparison of data and prediction for the six pest profiles

SM5: Prediction of OPM outbreak from 1970 to 2018 in the north-west of France

## References

1. Baier, P., Pennerstorfer, J., & Schopf, A. (2007). PHENIPS-A comprehensive phenology model of Ips typographus (L.) (Col., Scolytinae) as a tool for hazard rating of bark beetle infestation. Forest Ecology and Management, 249(3), 171–186. 10.1016/j.foreco.2007.05.020

2. Beaumont, M. a, Zhang, W., & Balding, D. J. (2002). Approximate Bayesian computation in population genetics. Genetics, 162(4), 2025–2035. http://www.pubmedcentral.nih.gov/articlerender.fcgi?artid=1462356&tool=pmcentrez&rendertype=abstract

3. Berryman, A., & Stark, R. W. (1985). Assessing the risk of forest insect outbreaks. Zeitschrift Für Angewandte Entomologie, 99(1–5), 199–208. 10.1111/j.1439-0418.1985.tb01979.x

4. Bloch, I. (1996). Information Combination Operators for Data Fusion: A Comparative Review with Classification. *TRANSACTIONS ON SYSTEMS, MAN*, AND CYBERNETICS, 26(1), 52–67.

5. Bloch, I., & Maitre, H. (1994). Fusion de données en traitement d’images: modèles d’information et de décision. Traitement Du Signal, 11(6), 435–446.

6. Branco, M., Brockerhoff, E. G., Castagneyrol, B., Orazio, C., & Jactel, H. (2015). Host range expansion of native insects to exotic trees increases with area of introduction and the presence of congeneric native trees. Journal of Applied Ecology, 52(1), 69–77. 10.1111/1365-2664.12362

7. Brockerhoff, E. G., & Liebhold, A. M. (2017). Ecology of forest insect invasions. Biological Invasions, 19(11), 3141–3159. 10.1007/s10530-017-1514-1

8. Brus, D. J., Hengeveld, G. M., Walvoort, D. J. J., Goedhart, P. W., Heidema, A. H., Nabuurs, G. J., & Gunia, K. (2012). Statistical mapping of tree species over Europe. European Journal of Forest Research, 131(1), 145–157. 10.1007/s10342-011-0513-5

9. Buse, J., Schröder, B., & Assmann, T. (2007). Modelling habitat and spatial distribution of an endangered longhorn beetle - A case study for saproxylic insect conservation. Biological Conservation, 137(3), 372–381. 10.1016/j.biocon.2007.02.025

10. Chang, W., Cheng, J., Allaire, J., Xie, Y., & McPherson, J. (2019). shiny: Web Application Framework for R. https://cran.r-project.org/package=shiny

11. Collot, D., & Robinet, C. (2021). R script to simulate the probability of outbreak of forest pests. Zenodo. 10.5281/zenodo.5112868

12. Cours, J., Nageleisen, L.-M., & Touffait, R. (2019). Gestion forestière intégrée des insectes ravageurs : exemple par l’étude de la niche écologique du hanneton forestier (Melolontha hippocastani Fabr. 1801). Revue Forestière Française, 2017(6), 553. 10.4267/2042/70886

13. Csóka, G., Hirka, A., Szocs, L., Móricz, N., Rasztovits, E., & Pödör, Z. (2018). Weather-dependent fluctuations in the abundance of the oak processionary moth, Thaumetopoea processionea (Lepidoptera: Notodontidae). European Journal of Entomology, 115(2012), 249–255. 10.14411/eje.2018.024

14. Douma, J. C., Van Der Werf, W., Hemerik, L., Magnusson, C., & Robinet, C. (2017). Development of a pathway model to assess the exposure of European pine trees to pine wood nematode via the trade of wood. Ecological Applications, 27(3), 769–785. 10.1002/eap.1480

15. Esper, J., Büntgen, U., Frank, D. C., Nievergelt, D., & Liebhold, A. (2007). 1200 Years of regular outbreaks in alpine insects. Proceedings of the Royal Society B: Biological Sciences, 274(1610), 671–679. 10.1098/rspb.2006.0191

16. Faccoli, M. (2009). Effect of Weather on Ips typographus (Coleoptera Curculionidae) Phenology, Voltinism, and Associated Spruce Mortality in the Southeastern Alps . Environmental Entomology, 38(2), 307–316. 10.1603/022.038.0202

17. Fawcett, T. (2006). An introduction to ROC analysis. Pattern Recognition Letters, 27(8), 861–874. 10.1016/j.patrec.2005.10.010

18. Fick, S. E., & Hijmans, R. J. (2017). WorldClim 2: new 1-km spatial resolution climate surfaces for global land areas. International Journal of Climatology, 37(12), 4302–4315. 10.1002/joc.5086

19. Godefroid, M., Meurisse, N., Groenen, F., Kerdelhué, C., & Rossi, J. P. (2020). Current and future distribution of the invasive oak processionary moth. Biological Invasions, 22(2), 523–534. 10.1007/s10530-019-02108-4

20. Gottschling, S., & Meyer, S. (2006). An epidemic airborne disease caused by the oak processionary caterpillar. Pediatric Dermatology, 23(1), 64–66. 10.1111/j.1525-1470.2006.00173.x

21. Groenen, F., & Meurisse, N. (2012). Historical distribution of the oak processionary moth Thaumetopoea processionea in Europe suggests recolonization instead of expansion. Agricultural and Forest Entomology, 14(2), 147–155. 10.1111/j.1461-9563.2011.00552.x

22. Harris, I., Jones, P. D., Osborn, T. J., & Lister, D. H. (2014). Updated high-resolution grids of monthly climatic observations - the CRU TS3.10 Dataset. International Journal of Climatology, 34(3), 623–642. 10.1002/joc.3711

23. Havašová, M., Ferenčík, J., & Jakuš, R. (2017). Interactions between windthrow, bark beetles and forest management in the Tatra national parks. Forest Ecology and Management, 391, 349–361. 10.1016/j.foreco.2017.01.009

24. Haynes, K. J., Allstadt, A. J., & Klimetzek, D. (2014). Forest defoliator outbreaks under climate change: Effects on the frequency and severity of outbreaks of five pine insect pests. Global Change Biology, 20(6), 2004–2018. 10.1111/gcb.12506

25. Hentschel, R., Möller, K., Wenning, A., Degenhardt, A., & Schröder, J. (2018). Importance of ecological variables in explaining population dynamics of three important pine pest insects. Frontiers in Plant Science, 871(November), 1–17. 10.3389/fpls.2018.01667

26. Holmes, T. P. (1991). Price and Welfare Effects of Catastrophic Forest Damage From Southern Pine- Beetle Epidemics. Forest Science, 37(2), 500–516.

27. Ismail, R., Mutanga, O., & Kumar, L. (2010). Modeling the Potential Distribution of Pine Forests Susceptible to Sirex Noctilio Infestations in Mpumalanga, South Africa. Transactions in GIS, 14(5), 709–726. 10.1111/j.1467-9671.2010.01229.x

28. Kosunen, M., Kantola, T., Starr, M., Blomqvist, M., Talvitie, M., & Lyytikäinen-Saarenmaa, P. (2017). Influence of soil and topography on defoliation intensity during an extended outbreak of the common pine sawfly (Diprion pini L.). IForest, 10(1), 164–171. 10.3832/ifor2069-009

29. Kriticos, D. J., Leriche, A., Palmer, D. J., Cook, D. C., Brockerhoff, E. G., Stephens, A. E. A., & Watt, M. S. (2013). Linking Climate Suitability, Spread Rates and Host-Impact When Estimating the Potential Costs of Invasive Pests. PLoS ONE, 8(2), 1–12. 10.1371/journal.pone.0054861

30. Kriticos, D., Maywald, G., Yonow, T., Zurcher, E., Herrmann, N., & Sutherst, R. (2016). CLIMEX Version 4: Exploring the Effects of Climate on Plants, Animals and Diseases. CSIRO.

31. Lesniak, A. (1976). Forest stand and site conditions of a pine moth outbreaks. Ekologia Polska, 24(4), 549–563.

32. Li, S., Daudin, J. J., Piou, D., Robinet, C., & Jactel, H. (2015). Periodicity and synchrony of pine processionary moth outbreaks in France. Forest Ecology and Management, 354, 309–317. 10.1016/j.foreco.2015.05.023

33. Lima, M., Harrington, R., Saldaña, S., & Estay, S. (2008). Non-linear feedback processes and a latitudinal gradient in the climatic effects determine green spruce aphid outbreaks in the UK. Oikos, 117(6), 951–959. 10.1111/j.0030-1299.2008.16615.x

34. Matek, M., & Pernek, M. (2018). First record of Dendrolimus pini outbreak on aleppo pine in Croatia and severe case of population collapse caused by entomopathogen Beauveria bassiana. South- East European Forestry, 9(2), 91–96. 10.15177/seefor.18-17

35. Möller, K., Hentschel, R., Wenning, A., & Schröder, J. (2017). Improved outbreak prediction for common pine sawfly (Diprion pini L.) by analyzing floating “climatic windows” as keys for changes in voltinism. Forests, 8(9). 10.3390/f8090319

36. Moraal, L. G., & Hilszczanski, J. (2000). The oak buprestid beetle, Agrilus biguttatus (F.) (Col., Buprestidae), a recent factor in oak decline in Europe. Anzeiger Fur Schadlingskunde, 73(5), 134–138. 10.1007/BF02956447

37. Nageleisen, L.-M., Piou, D., Saintonge, F.-X., & Riou-Nivert, P. (2009). *La santé des Forêts - Maladies, insectes, accidents climatiques… Diagnostic et prévention*. Ministère de l’Agriculture et de la Pêche; Département de la Santé des Forêts; Paris.

38. Parry, H., Sadler, R., & Kriticos, D. (2013). Practical guidelines for modelling post-entry spread in invasion ecology. NeoBiota, 18, 41–66. 10.3897/neobiota.18.4305

39. Pellicone, G., Caloiero, T., & Guagliardi, I. (2019). The De Martonne aridity index in Calabria (Southern Italy). Journal of Maps, 15(2), 788–796. 10.1080/17445647.2019.1673840

40. R Core Team. (2018). R: A Language and Environment for Statistical Computing. https://www.r-project.org/

41. Ramsfield, T. D., Bentz, B. J., Faccoli, M., Jactel, H., & Brockerhoff, E. G. (2016). Forest health in a changing world: Effects of globalization and climate change on forest insect and pathogen impacts. Forestry, 89(3), 245–252. 10.1093/forestry/cpw018

42. Robinet, C., Kehlenbeck, H., Kriticos, D. J., Baker, R. H. A., Battisti, A., Brunel, S., Dupin, M., Eyre, D., Faccoli, M., Ilieva, Z., Kenis, M., Knight, J., Reynaud, P., Yart, A., & van der Werf, W. (2012). A Suite of Models to Support the Quantitative Assessment of Spread in Pest Risk Analysis. PLoS ONE, 7(10). 10.1371/journal.pone.0043366

43. Robinet, C., Van Den Dool, R., Collot, D., & Douma, J. (2020). Modelling for risk and biosecurity related to forest health. Emerging Topics in Life Sciences, 4(5).

44. Rozenberg, P., Pâques, L., Huard, F., & Roques, A. (2020). Direct and Indirect Analysis of the Elevational Shift of Larch Budmoth Outbreaks Along an Elevation Gradient. Frontiers in Forests and Global Change, 3(July). 10.3389/ffgc.2020.00086

45. Soliman, T., Mourits, M. C. M., van der Werf, W., Hengeveld, G. M., Robinet, C., & Lansink, A. G. J. M. O. (2012). Framework for Modelling Economic Impacts of Invasive Species, Applied to Pine Wood Nematode in Europe. PLoS ONE, 7(9), 1–12. 10.1371/journal.pone.0045505

46. Toni, T., Welch, D., Strelkowa, N., Ipsen, A., & Stumpf, M. P. H. (2009). Approximate Bayesian computation scheme for parameter inference and model selection in dynamical systems. Journal of the Royal Society Interface, 6, 187–202. 10.1098/rsif.2008.0172

47. Turchin, P., Wood, S. N., Ellner, S. P., Kendall, B. E., Murdoch, W. W., Fischlin, A., Casas, J., McCauley, E., & Briggs, C. J. (2003). Dynamical effects of plant quality and parasitism on population cycles of larch budmoth. Ecology, 84(5), 1207–1214. 10.1890/0012-9658(2003)084[1207:DEOPQA]2.0.CO;2

48. Valenta, V., Moser, D., Kapeller, S., & Essl, F. (2017). A new forest pest in Europe: a review of Emerald ash borer (Agrilus planipennis) invasion. Journal of Applied Entomology, 141(7), 507–526. 10.1111/jen.12369

