## Supplementary Material 1 for "Probability of outbreaks of forest insects in Europe: a generic model calibrated on six forest insect profiles"

SM1: Print screens of Shiny Ap interface developed for this generic outbreak model.

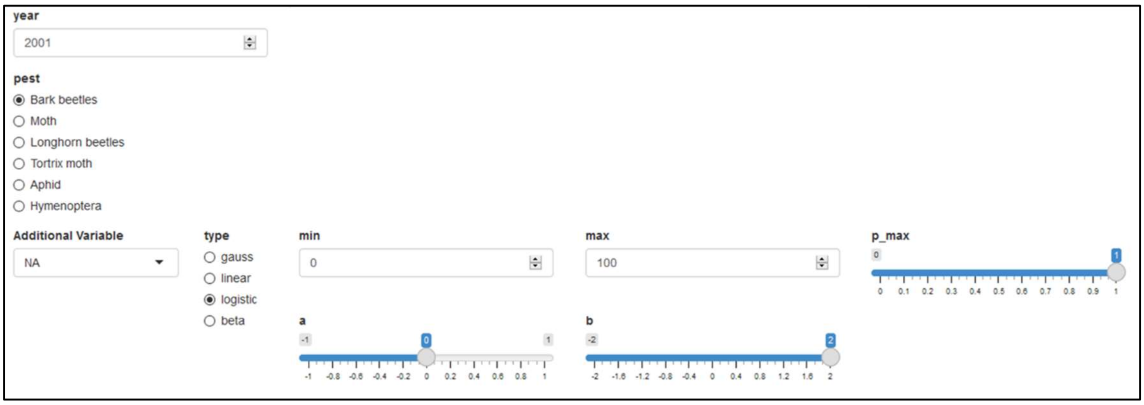


Figure 1: Panel where the user can select the insect profile, and eventually an additional driver.

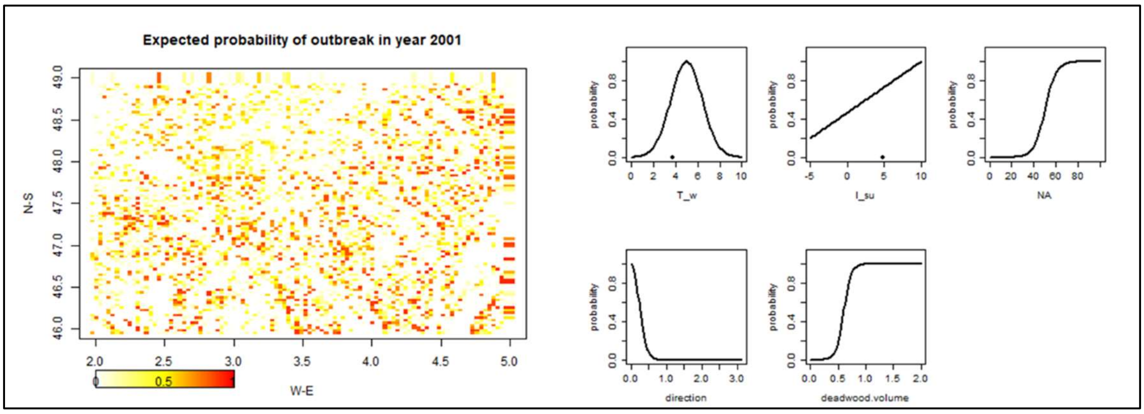


Figure 2: Panel showing the shape of the probability functions for each single driver (on the right) and the simulated overall outbreak probability on the study area (on the left).

| Species | mean probability of outbreak | % of area with a probability > 0.7 | |
|---|---|---|---|
| Bark beetles | 0.14 | 7 | |
| Moth | 0.22 | 0 | |
| Longhorn beetles | 0.01 | 0 | |
| Tortrix moth | 0.11 | 0 | |
| Aphid | 0.62 | 10 | |
| Hymenoptera | 0.73 | 81 | ** |

Figure 3: Panel showing two summary variables related to the outbreak probability for the 6 insect profiles.
