## Supplementary Material 2 for "Probability of outbreaks of forest insects in Europe: a generic model calibrated on six forest insect profiles"

Supplementary Material 2: List of the 147 European forest insects considered to build profiles

| **Species** | **Order** | **Family** |
|---|---|---|
| *Acanthocinus aedilis* | coleoptera | Cerambycidae |
| *Acantholyda hieroglyphica* | hymenoptera | Pamphiliidae |
| *Acronicta aceris* | lepidoptera | Noctuidae |
| *Acronicta rumicis* | lepidoptera | Noctuidae |
| *Actebia fennica* | lepidoptera | Noctuidae |
| *Adelges (Gilletteella) cooleyi* | homoptera | Adelgidae |
| *Adelges laricis* | homoptera | Adelgidae |
| *Agelastica alni* | coleoptera | Chrysomaelidae |
| *Agrilus biguttatus* | coleoptera | Buprestidae |
| *Agrilus suvorovi (ater)* | coleoptera | Buprestidae |
| *Agrilus viridis* | coleoptera | Buprestidae |
| *Agriopis aurantiaria* | lepidoptera | Geometridae |
| *Agrotis segetum* | lepidoptera | Noctuidae |
| *Alcis repandata* | lepidoptera | Geometridae |
| *Alsophila aescularia* | lepidoptera | Geometridae |
| *Altica quercetorum* | coleoptera | Chrysomaelidae |
| *Amphipyra pyramidea* | lepidoptera | Noctuidae |
| *Anthonomus phyllocola* | coleoptera | Curculionidaes |
| *Apoderus coryli* | coleoptera | Curculionidaes |
| *Aradus cinnamomeus* | hemiptera | Aradidae |
| *Archarius salicivorus* | coleoptera | Curculionidaes |
| *Archips crataegana* | lepidoptera | Tortricidae |
| *Archips oporana* | lepidoptera | Tortricidae |
| *Archips xylosteana* | lepidoptera | Tortricidae |
| *Argyresthia dilectella* | lepidoptera | Yponomeutidae |
| *Argyresthia glabratella* | lepidoptera | Yponomeutidae |
| *Argyresthia laevigatella* | lepidoptera | Yponomeutidae |
| *Asterodiaspis quercicola* | hemiptera | Coccoidae |
| *Attelabus nitens* | coleoptera | Curculionidaes |
| *Bena bicolorana* | lepidoptera | Nolidae |
| *Biston betularia* | lepidoptera | Geometridae |
| *Blastesthia posticana* | lepidoptera | Tortricidae |
| *Brachyderes incanus* | coleoptera | Curculionidaes |
| *Bupalus piniaria* | lepidoptera | Geometridae |
| *Caliroa cerasi* | hymenoptera | Tenthredinidae |
| *Calliteara pudibunda* | lepidoptera | Lymantriidae |
| *Calomicrus pinicola* | coleoptera | Chrysomaelidae |
| *Carulaspis juniperi (Diaspis visci)* | hemiptera | Diaspididae |
| *Carulaspis minima* | hemiptera | Diaspididae |
| *Cephalcia abietis* | hymenoptera | Pamphiliidae |
| *Cerambyx cerdo* | coleoptera | Cerambycidae |
| *Chaitophorus leucomelas* | hemiptera | Aphididae |

| | | |
|---|---|---|
| *Chionaspis salicis* | hemiptera | Diaspididae |
| *Choristoneura murinana* | lepidoptera | Tortricidae |
| *Chrysobothris igniventris* | coleoptera | Buprestidae |
| *Chrysomela populi* | coleoptera | Chrysomaelidae |
| *Cimbex luteus* | hymenoptera | Tenthredinidae |
| *Cinara piceae* | hemiptera | Aphididae |
| *Cinara pilicornis* | hemiptera | Aphididae |
| *Clytus arietis* | coleoptera | Cerambycidae |
| *Coleophora laricella* | lepidoptera | Coleophoridae |
| *Colotois pennaria* | lepidoptera | Geometridae |
| *Contarinia sp.* | diptera | Cecidomyiidae |
| *Coraebus (Coroebus) florentinus* | coleoptera | Buprestidae |
| *Cosmia trapezina* | lepidoptera | Noctuidae |
| *Cossus cossus* | lepidoptera | Cossidae |
| *Crocallis elinguaria* | lepidoptera | Geometridae |
| *Cryphalus abietis* | coleoptera | bark beetle |
| *Cryphalus abietis* | coleoptera | bark beetle |
| *Cryphalus piceae* | coleoptera | bark beetle |
| *Cryphalus piceae* | coleoptera | bark beetle |
| *Cryptocephalus pini* | coleoptera | Chrysomaelidae |
| *Cryptorhynchus lapathi* | coleoptera | Curculionidaes |
| *Crypturgus pusillus* | coleoptera | bark beetle |
| *Cucujus cinnaberinus* | coleoptera | Cucujidae |
| *Cucujus cinnaberinus* | coleoptera | Cucujidae |
| *Curculio elephas* | coleoptera | Curculionidaes |
| *Curculio glandium* | coleoptera | Curculionidaes |
| *Cydia splendana* | lepidoptera | Tortricidae |
| *Cydia strobilella* | lepidoptera | Tortricidae |
| *Dasineura laricis* | diptera | Cecidomyiidae |
| *Deileptenia ribeata* | lepidoptera | Geometridae |
| *Dendroctonus micans* | coleoptera | bark beetle |
| *Dendrolimus pini* | lepidoptera | Lasiocampidae |
| *Diaspidiotus perniciosus* | hemiptera | Coccoidae |
| *Diaspidiotus perniciosus* | hemiptera | Coccoidae |
| *Diaspidiotus perniciosus* | hemiptera | Coccoidae |
| *Diaspidiotus perniciosus* | hemiptera | Coccoidae |
| *Diaspidiotus perniciosus* | hemiptera | Coccoidae |
| *Diaspidiotus perniciosus* | hemiptera | Coccoidae |
| *Diaspidiotus perniciosus* | hemiptera | Coccoidae |
| *Diaspidiotus perniciosus* | hemiptera | Coccoidae |
| *Diloba caeruleocephala* | lepidoptera | Noctuidae |
| *Dioryctria mendacella* | lepidoptera | Pyralidae |
| *Dioryctria mutatella* | lepidoptera | Pyralidae |
| *Dioryctria mutatella* | lepidoptera | Pyralidae |
| *Dioryctria sylvestrella* | lepidoptera | Pyralidae |
| *Diprion pini* | hymenoptera | Tenthredinidae |

| | | |
|---|---|---|
| *Diprion similis* | hymenoptera | Tenthredinidae |
| *Ditula angustiorana* | lepidoptera | Tortricidae |
| *Drepanosiphum platanoidis* | hemiptera | Aphididae |
| *Dreyfusia nordmannianae* | homoptera | Adelgidae |
| *Dreyfusia nordmannianae* | homoptera | Adelgidae |
| *Dryocoetes autographus* | coleoptera | bark beetle |
| *Dryocoetes autographus* | coleoptera | bark beetle |
| *Dynaspidiotus (Nuculaspis) abietis* | hemiptera | Coccoidae |
| *Dynaspidiotus (Nuculaspis) abietis* | hemiptera | Coccoidae |
| *Dysaphis aucupariae* | hemiptera | Aphididae |
| *Ectropis crepuscularia* | lepidoptera | Geometridae |
| *Elateroides (Hylecoetus) dermestoides* | coleoptera | Lymexylidae |
| *Elatobium abietinum* | hemiptera | Aphididae |
| *Ennomos quercaria* | lepidoptera | Geometridae |
| *Eotetranychus pruni* | trombidiforme | Tetrachynidae |
| *Epinotia tedella* | lepidoptera | Tortricidae |
| *Epirrita autumnata* | lepidoptera | Geometridae |
| *Epirrita dilutata* | lepidoptera | Geometridae |
| *Erannis defoliaria* | lepidoptera | Geometridae |
| *Ergates faber* | coleoptera | Cerambycidae |
| *Ernobius mollis* | coleoptera | Ptinidae |
| *Eupithecia pusillata* | lepidoptera | Geometridae |
| *Eupithecia pusillata* | lepidoptera | Geometridae |
| *Euproctis chrysorrhoea* | lepidoptera | Lymantriidae |
| *Euproctis similis* | lepidoptera | Lymantriidae |
| *Exoteleia dodecella* | lepidoptera | Gelechiidae |
| *Furcipus rectirostris* | coleoptera | Curculionidaes |
| *Gastrodes grossipes* | heteroptera | Lygaeidae |
| *Gilpinia hercyniae* | hymenoptera | Diprionidae |
| *Gonioctena quinquepunctata* | coleoptera | Chrysomaelidae |
| *Gypsonoma aceriana* | lepidoptera | Tortricidae |
| *Haematoloma dorsatum* | hemiptera | Cercopidae |
| *Hapleginella laevifrons* | diptera | Chloropidae |
| *Hapleginella laevifrons* | diptera | Chloropidae |
| *Hemithea aestivaria* | lepidoptera | Geometridae |
| *Herophila (Dorcatypus) tristis* | coleoptera | Cerambycidae |
| *Hylaea fasciaria* | lepidoptera | Geometridae |
| *Hylastes ater* | coleoptera | bark beetle |
| *Hylesinus fraxini* | coleoptera | bark beetle |
| *Hylobius abietis* | coleoptera | Curculionidaes |
| *Hylobius piceus* | coleoptera | Curculionidaes |
| *Hylobius piceus* | coleoptera | Curculionidaes |
| *Hylotrupes bajulus* | coleoptera | Cerambycidae |
| *Hylurgops palliatus* | coleoptera | bark beetle |
| *Hylurgus ligniperda* | coleoptera | bark beetle |
| *Hypothenemus eruditus* | coleoptera | bark beetle |

| | | |
|---|---|---|
| *Icosium tomentosum* | coleoptera | Cerambycidae |
| *Ips typographus* | coleoptera | bark beetle |
| *Kleidocerys resedae* | heteroptera | Lygaeidae |
| *Lasiocampa quercus* | lepidoptera | Lasiocampidae |
| *Leucaspis pini* | hemiptera | Coccoidae |
| *Leucoma salicis* | lepidoptera | Erebidae |
| *Lithophane leautieri* | lepidoptera | Noctuidae |
| *Lomographa bimaculata* | lepidoptera | Geometridae |
| *Lucasianus levaillantii* | coleoptera | Cerambycidae |
| *Luperus pinicola* | coleoptera | Chrysomaelidae |
| *Lymantria dispar* | lepidoptera | Erebidae |
| *Lymantria monacha* | lepidoptera | Erebidae |
| *Lymexylon navale* | coleoptera | Lymexylidae |
| *Macaria liturata* | lepidoptera | Geometridae |
| *Magdalis duplicata* | coleoptera | Curculionidaes |
| *Malacosoma neustria* | lepidoptera | Lasiocampidae |
| *Melanchra pisi* | lepidoptera | Noctuidae |
| *Melolontha hippocastani* | coleoptera | Scarabaeidea |
| *Melolontha hippocastani* | coleoptera | Scarabaeidea |
| *Melolontha melolontha* | coleoptera | Scarabaeidea |
| *Melolontha melolontha* | coleoptera | Scarabaeidea |
| *Microdiprion pallipes* | hymenoptera | Diprionidae |
| *Microdiprion pallipes* | hymenoptera | Diprionidae |
| *Mindarus abietinus* | hemiptera | Aphididae |
| *Molorchus minor* | coleoptera | Cerambycidae |
| *Monochamus sutor* | coleoptera | Cerambycidae |
| *Morimus asper* | coleoptera | Cerambycidae |
| *Nathrius brevipennis* | coleoptera | Cerambycidae |
| *Neodiprion sertifer* | hymenoptera | Diprionidae |
| *Neohydatothrips gracilicornis* | thysanoptera | Thripidae |
| *Oligonychus ununguis* | trombidiforme | Tetrachynidae |
| *Operophtera brumata* | lepidoptera | Geometridae |
| *Orchestes fagi* | coleoptera | Curculionidaes |
| *Orchestes quercus* | coleoptera | Curculionidaes |
| *Orgyia antiqua* | lepidoptera | Erebidae |
| *Orsillus maculatus/ depressus* | heteroptera | Lygaeidae |
| *Orthosia cerasi (stabilis)* | lepidoptera | Noctuidae |
| *Orthosia gothica* | lepidoptera | Noctuidae |
| *Orthosia incerta* | lepidoptera | Noctuidae |
| *Orthotomicus erosus* | coleoptera | bark beetle |
| *Orthotomicus erosus* | coleoptera | bark beetle |
| *Orthotomicus laricis* | coleoptera | bark beetle |
| *Otiorhynchus armadillo* | coleoptera | Curculionidaes |
| *Ovalisia (Palmar) festiva* | coleoptera | Buprestidae |
| *Panolis flammea* | lepidoptera | Noctuidae |
| *Paranthrene tabaniformis* | lepidoptera | Sesiidae |

| | | |
|---|---|---|
| *Parthenolecanium rufulum* | hemiptera | Coccoidae |
| *Pemphigus spirothecae* | hemiptera | Aphididae |
| *Penichroa fasciata* | coleoptera | Cerambycidae |
| *Phalera bucephala* | lepidoptera | Notodontidae |
| *Phigalia pilosaria* | lepidoptera | Geometridae |
| *Phloeomyzus passerinii* | hemiptera | Aphididae |
| *Phloeosinus aubei* | coleoptera | bark beetle |
| *Phloeosinus bicolor/ thujae* | coleoptera | bark beetle |
| *Phyllaphis fagi* | hemiptera | Aphididae |
| *Phyllobius pyri* | coleoptera | Curculionidaes |
| *Phyllocnistis suffusella* | lepidoptera | Gracillaridae |
| *Phyllocnistis unipunctella* | lepidoptera | Gracillaridae |
| *Phyllopertha horticola* | coleoptera | Scarabaeidea |
| *Phymatodes glabratus* | coleoptera | Cerambycidae |
| *Phytobia cambii* | diptera | Agromyzidae |
| *Pineus pini* | hemiptera | Phylloxeroidae |
| *Pissodes castaneus* | coleoptera | Curculionidaes |
| *Pissodes castaneus* | coleoptera | Curculionidaes |
| *Pissodes piceae* | coleoptera | Curculionidaes |
| *Pissodes piceae* | coleoptera | Curculionidaes |
| *Pissodes validirostris* | coleoptera | Curculionidaes |
| *Pityogenes bidentatus* | coleoptera | bark beetle |
| *Pityogenes chalcographus* | coleoptera | bark beetle |
| *Pityogenes chalcographus* | coleoptera | bark beetle |
| *Pityokteines curvidens* | coleoptera | bark beetle |
| *Pityokteines curvidens* | coleoptera | bark beetle |
| *Pityokteines curvidens* | coleoptera | bark beetle |
| *Pityokteines spinidens* | coleoptera | bark beetle |
| *Pityophthorus pityographus* | coleoptera | bark beetle |
| *Planococcus vovae* | hemiptera | Pseudococcidae |
| *Planococcus vovae* | hemiptera | Pseudococcidae |
| *Platypus cylindrus* | coleoptera | Ambrosia beetle |
| *Pogonocherus fasciculatus* | coleoptera | Cerambycidae |
| *Polydrusus formosus* | coleoptera | Curculionidaes |
| *Polygraphus punctifrons* | coleoptera | bark beetle |
| *Polyphylla fullo* | coleoptera | Scarabaeidea |
| *Polyphylla fullo* | coleoptera | Scarabaeidea |
| *Prionus coriarius* | coleoptera | Cerambycidae |
| *Pristiphora abietina* | hymenoptera | Tenthredinidae |
| *Prociphilus fraxini* | hemiptera | Aphididae |
| *Prociphilus fraxini* | hemiptera | Aphididae |
| *Pseudococcyx tessulatana* | lepidoptera | Tortricidae |
| *Ptycholoma lecheana* | lepidoptera | Tortricidae |
| *Retinia resinella* | lepidoptera | Tortricidae |
| *Rhagium bifasciatum* | coleoptera | Cerambycidae |
| *Rhyacionia buoliana* | lepidoptera | Tortricidae |

| | | |
|---|---|---|
| *Rhyacionia buoliana* | lepidoptera | Tortricidae |
| *Rhyacionia duplana* | lepidoptera | Tortricidae |
| *Rhyacionia pinicolana* | lepidoptera | Tortricidae |
| *Sacchiphantes viridis* | homoptera | Adelgidae |
| *Saperda carcharias* | coleoptera | Cerambycidae |
| *Saperda carcharias* | coleoptera | Cerambycidae |
| *Saperda carcharias* | coleoptera | Cerambycidae |
| *Saperda carcharias* | coleoptera | Cerambycidae |
| *Saperda populnea* | coleoptera | Cerambycidae |
| *Schizolachnus pineti* | hemiptera | Aphididae |
| *Scolytus intricatus* | coleoptera | bark beetle |
| *Semanotus laurasi* | coleoptera | Cerambycidae |
| *Semanotus russicus* | coleoptera | Cerambycidae |
| *Sesia apiformis* | lepidoptera | Sesiidae |
| *Sphinx (Hyloicus) pinastri* | lepidoptera | Sphingidae |
| *Stephanopachys substriatus* | coleoptera | Bostrichidae |
| *Stictoleptura (Corymbia) rubra* | coleoptera | Cerambycidae |
| *Stigmella trimaculella* | lepidoptera | Nepticulidae |
| *Strobilomyia anthracina* | diptera | Anthomyiidae |
| *Strophosoma melanogrammum* | coleoptera | Curculionidaes |
| *Tetropium fuscum* | coleoptera | Cerambycidae |
| *Thaumetopoea pityocampa* | lepidoptera | Notodontidae |
| *Thaumetopoea processionea* | lepidoptera | Notodontidae |
| *Thecodiplosis brachyntera* | diptera | Cecidomyiidae |
| *Thera britannica* | lepidoptera | Geometridae |
| *Thera obeliscata* | lepidoptera | Geometridae |
| *Tischeria ekebladella* | lepidoptera | Tischeriidae |
| *Tomicus destruens* | coleoptera | bark beetle |
| *Tomicus piniperda* | coleoptera | bark beetle |
| *Tortrix viridana* | lepidoptera | Tortricidae |
| *Trichoferus fasciculatus* | coleoptera | Cerambycidae |
| *Trypodendron lineatum* | coleoptera | Ambrosia beetle |
| *Urocerus gigas* | hymenoptera | Siricidea |
| *Xeris spectrum* | hymenoptera | Siricidea |
| *Xyleborinus saxesenii* | coleoptera | Ambrosia beetle |
| *Xyleborinus saxesenii* | coleoptera | Ambrosia beetle |
| *Xyleborus dispar* | coleoptera | Ambrosia beetle |
| *Xyleborus monographus* | coleoptera | Ambrosia beetle |
| *Zeiraphera diniana* | lepidoptera | Tortricidae |
| *Zeuzera pyrina* | lepidoptera | Cossidae |
