## Supplementary Material 3 for "Probability of outbreaks of forest insects in Europe: a generic model calibrated on six forest insect profiles"

Supplementary Material 3: List of publications found and studied in the systematic review, later used to extract the drivers

| Authors | year | Title | Journal |
|---|---|---|---|
| Aakala, T; Kuuluvainen, T; De Grandpre, L; Gauthier, S | 2007 | Trees dying standing in the northeastern boreal old-growth forests of Quebec: spatial patterns, rates, and temporal variation | CANADIAN JOURNAL OF FOREST RESEARCH-REVUE CANADIENNE DE RECHERCHE FORESTIERE |
| Abdullah, H; Skidmore, AK; Darvishzadeh, R; Heurich, M | 2019 | Sentinel-2 accurately maps green-attack stage of European spruce bark beetle (Ips typographus, L.) compared with Landsat-8 | REMOTE SENSING IN ECOLOGY AND CONSERVATION |
| Aimi, A; Larsson, S; Ronnas, C; Frazao, J; Battisti, A | 2008 | Growth and survival of larvae of Thaumetopoea pinivora inside and outside a local outbreak area | AGRICULTURAL AND FOREST ENTOMOLOGY |
| Akbulut, S; Yuksel, B; Baysal, I; Vieira, P; Mota, M | 2008 | Pine Wilt Disease: A Threat to Pine Forests in Turkey? | PINE WILT DISEASE: A WORLDWIDE THREAT TO FOREST ECOSYSTEMS |
| Akinci, HA; Aksu, Y | 2018 | Analyzing the Local Spread of Ips typographus (L.) (Coleoptera: Curculionidae, Scolytinae) by Pheromone Catches in Turkey's Hatila Valley National Park | POLISH JOURNAL OF ENVIRONMENTAL STUDIES |
| Akkuzu, E; Guner, S | 2008 | Defoliation levels of oriental spruce by Ips typographus (L.) in relation to elevation and exposure | JOURNAL OF ENVIRONMENTAL BIOLOGY |
| Akkuzu, E; Guzel, H | 2015 | EDGE EFFECTS OF Pinus nigra FORESTS ON ABUNDANCE AND BODY LENGHT OF Ips sexdentatus | SUMARSKI LIST |
| Alalouni, U; Schadler, M; Brandl, R | 2013 | Natural enemies and environmental factors affecting the population dynamics of the gypsy moth | JOURNAL OF APPLIED ENTOMOLOGY |
| Amezaga, I | 1996 | Monterrey pine (Pinus radiata D Don) suitability for the pine shoot beetle (Tomicus piniperda L) (Coleoptera: Scolytidae) | FOREST ECOLOGY AND MANAGEMENT |

| | | | |
|---|---|---|---|
| Ammunet, T; Heisswolf, A; Klemola, N; Klemola, T | 2010 | Expansion of the winter moth outbreak range: no restrictive effects of competition with the resident autumnal moth | ECOLOGICAL ENTOMOLOGY |
| Arac, K; Pernek, M | 2014 | OCCURRENCE AND SPREADING OF THE LARGE LARCH BARK BEETLE (Ips cembrae) IN CROATIA AND POSSIBLITIES OF MONITORING BY USING PHEROMONE TRAPS | SUMARSKI LIST |
| Arnberger, A; Ebenberger, M; Schneider, IE; Cottrell, S; Schlueter, AC; von Ruschkowski, E; Venette, RC; Snyder, SA; Gobster, PH | 2018 | Visitor Preferences for Visual Changes in Bark Beetle-Impacted Forest Recreation Settings in the United States and Germany | ENVIRONMENTAL MANAGEMENT |
| Arnberger, A; Eder, R; Allex, B; Preisel, H; Ebenberger, M; Husslein, M | 2018 | Trade-offs between wind energy, recreational, and bark-beetle impacts on visual preferences of national park visitors | LAND USE POLICY |
| Arnold, C; Bachmann, O; Schnitzler, A | 2017 | Insights into the Vitis complex in the Danube floodplain (Austria) | ECOLOGY AND EVOLUTION |
| Augustaitis, A | 2007 | Pine sawfly (Diprion pini l.) - Related changes in Scots pine crown defoliation and possibilities of recovery | POLISH JOURNAL OF ENVIRONMENTAL STUDIES |
| Avtzis, N; Avtzis, D | 2003 | The attack of Aesculus hippocastanum L. by Cameraria ohridella Deschka and Dimic (Lepidoptera : Gracillariidae) in Greece | ECOLOGY, SURVEY AND MANAGEMENT OF FOREST INSECTS, PROCEEDINGS |
| Awan, HUM; Pettenella, D | 2017 | Pine Nuts: A Review of Recent Sanitary Conditions and Market Development | FORESTS |
| Babst, F; Esper, J; Parlow, E | 2010 | Landsat TM/ETM plus and tree-ring based assessment of spatiotemporal patterns of the autumnal moth (Epirrita autumnata) in northernmost Fennoscandia | REMOTE SENSING OF ENVIRONMENT |

| | | | |
|---|---|---|---|
| Baders, E; Jansons, A; Matisons, R; Elferts, D; Desaine, I | 2018 | Landscape Diversity for Reduced Risk of Insect Damage: A Case Study of Spruce Bud Scale in Latvia | FORESTS |
| Balciauskas, L; Balciauskiene, L; Stirke, V | 2019 | Mow the Grass at the Mouse's Peril: Diversity of Small Mammals in Commercial Fruit Farms | ANIMALS |
| Baldrian, P | 2017 | Forest microbiome: diversity, complexity and dynamics | FEMS MICROBIOLOGY REVIEWS |
| Band, HT; Bachli, G; Band, RN | 2005 | Behavioral constancy for interspecies dependency enables Nearctic Chymomyza amoena (Loew) (Diptera : Drosophilidae) to spread in orchards and forests in Central and Southern Europe | BIOLOGICAL INVASIONS |
| Barrios, JM; Verstraeten, WW; Maes, P; Aerts, JM; Farifteh, J; Coppin, P | 2013 | Relating land cover and spatial distribution of nephropathia epidemica and Lyme borreliosis in Belgium | INTERNATIONAL JOURNAL OF ENVIRONMENTAL HEALTH RESEARCH |
| Barron, E; Averyanova, A; Kvacek, Z; Momohara, A; Pigg, KB; Popova, S; Postigo-Mijarra, JM; Tiffney, BH; Utescher, T; Zhou, ZK | 2017 | The Fossil History of Quercus | OAKS PHYSIOLOGICAL ECOLOGY. EXPLORING THE FUNCTIONAL DIVERSITY OF GENUS QUERCUS L. |
| Bassler, C; Muller, J; Svoboda, M; Lepsova, A; Hahn, C; Holzer, H; Pouska, V | 2012 | Diversity of wood-decaying fungi under different disturbance regimes-a case study from spruce mountain forests | BIODIVERSITY AND CONSERVATION |
| Bebi, P; Seidl, R; Motta, R; Fuhr, M; Firm, D; Krumm, F; Conedera, M; Ginzler, C; Wohlgemuth, T; Kulakowski, D | 2017 | Changes of forest cover and disturbance regimes in the mountain forests of the Alps | FOREST ECOLOGY AND MANAGEMENT |
| Bednarz, B; Kacprzyk, M | 2012 | An Innovative Method for Sex Determination of the European Spruce Bark Beetle Ips typographus (Coleoptera: Scolytinae) | ENTOMOLOGIA GENERALIS |
| Bendel, M; Kienast, F; Rigling, D | 2006 | Genetic population structure of three Armillaria species at the landscape scale: a case | MYCOLOGICAL RESEARCH |

| | | | |
|---|---|---|---|
| | | study from Swiss Pinus mugo forests | |
| Berec, L; Dolezal, P; Hais, M | 2013 | Population dynamics of Ips typographus in the Bohemian Forest (Czech Republic): Validation of the phenology model PHENIPS and impacts of climate change | FOREST ECOLOGY AND MANAGEMENT |
| Bergseng, E; Okland, B; Gobakken, T; Magnusson, C; Rafoss, T; Solberg, B | 2012 | Combining ecological and economic modelling in analysing a pest invasion contingency plan - The case of pine wood nematode in Norway | SCANDINAVIAN JOURNAL OF FOREST RESEARCH |
| Bestard, AB; Font, AR | 2010 | Estimating the aggregate value of forest recreation in a regional context | JOURNAL OF FOREST ECONOMICS |
| Beudert, B; Bassler, C; Thorn, S; Noss, R; Schroder, B; Dieffenbach-Fries, H; Foullois, N; Muller, J | 2015 | Bark Beetles Increase Biodiversity While Maintaining Drinking Water Quality | CONSERVATION LETTERS |
| Beule, L; Gruning, MM; Karlovsky, P; L-M-Arnold, A | 2017 | Changes of Scots Pine Phyllosphere and Soil Fungal Communities during Outbreaks of Defoliating Insects | FORESTS |
| Bezos, D; Martinez-Alvarez, P; Fernandez, M; Diez, JJ | 2017 | Epidemiology and Management of Pine Pitch Canker Disease in Europe - a Review | BALTIC FORESTRY |
| Biedermann, PHW; Muller, J; Gregoire, JC; Gruppe, A; Hagge, J; Hammerbacher, A; Hofstetter, RW; Kandasamy, D; Kolarik, M; Kostovcik, M; Krokene, P; Salle, A; Six, DL; Turrini, T; Vanderpool, D; Wingfield, MJ; Bassler, C | 2019 | Bark Beetle Population Dynamics in the Anthropocene: Challenges and Solutions | TRENDS IN ECOLOGY & EVOLUTION |
| Biernat, B; Stanczak, J; Michalik, J; Sikora, B; Cieniuch, S | 2016 | Rickettsia helvetica and R-monacensis infections in immature Ixodes ricinus ticks derived from sylvatic passerine birds in west-central Poland | PARASITOLOGY RESEARCH |
| Billinis, C | 2013 | Wildlife diseases that pose a risk to small ruminants and their farmers | SMALL RUMINANT RESEARCH |
| Biondi, F; Hartsough, P | 2010 | Using Automated Point Dendrometers to Analyze Tropical Treeline Stem | SENSORS |

| | | | |
|---|---|---|---|
| | | Growth at Nevado de Colima, Mexico | |
| Biuw, M; Jepsen, JU; Cohen, J; Ahonen, SH; Tejesvi, M; Aikio, S; Wali, PR; Vindstad, OPL; Markkola, A; Niemela, P; Ims, RA | 2014 | Long-term Impacts of Contrasting Management of Large Ungulates in the Arctic Tundra-Forest Ecotone: Ecosystem Structure and Climate Feedback | ECOSYSTEMS |
| Bjerke, JW; Treharne, R; Vikhamar-Schuler, D; Karlsen, SR; Ravolainen, V; Bokhorst, S; Phoenix, GK; Bochenek, Z; Tommervik, H | 2017 | Understanding the drivers of extensive plant damage in boreal and Arctic ecosystems: Insights from field surveys in the aftermath of damage | SCIENCE OF THE TOTAL ENVIRONMENT |
| Blande, JD; Tiiva, P; Oksanen, E; Holopainen, JK | 2007 | Emission of herbivore-induced volatile terpenoids from two hybrid aspen (Populus tremula x tremuloides) clones under ambient and elevated ozone concentrations in the field | GLOBAL CHANGE BIOLOGY |
| Blank, L; Martin-Garcia, J; Bezos, D; Vettraino, AM; Krasnov, H; Lomba, JM; Fernandez, M; Diez, JJ | 2019 | Factors Affecting the Distribution of Pine Pitch Canker in Northern Spain | FORESTS |
| Blennow, K | 2012 | Adaptation of forest management to climate change among private individual forest owners in Sweden | FOREST POLICY AND ECONOMICS |
| Bobiec, A | 2002 | Living stands and dead wood in the Bialowieza forest: suggestions for restoration management | FOREST ECOLOGY AND MANAGEMENT |
| BOMBOSCH, S; DEDEK, W | 1994 | INTEGRATED PEST-CONTROL AGAINST IPS-TYPOGRAPHUS (L) - COMBINED USE OF PHEROMONES AND THE SYSTEMIC INSECTICIDE METHAMIDOPHOS (IPIDEX) | ZEITSCHRIFT FUR PFLANZENKRANKHEITEN UND PFLANZENSCHUTZ-JOURNAL OF PLANT DISEASES AND PROTECTION |
| Botella, L; Santamaria, O; Diez, JJ | 2010 | Fungi associated with the decline of Pinus halepensis in Spain | FUNGAL DIVERSITY |
| Bottero, A; Garbarino, M; Long, JN; Motta, R | 2013 | The interacting ecological effects of large-scale disturbances and salvage | FOREST ECOLOGY AND MANAGEMENT |

| | | | |
|---|---|---|---|
| | | logging on montane spruce forest regeneration in the western European Alps | |
| Bragard, C; Di Serio, F; Gonthier, P; Jacques, MA; Jaques Miret, JA; Justesen, AMF; MacLeod, A; Magnusson, CS; Milonas, P; Navas-Cortes, JA; Parnell, S; Potting, R; Reignault, PL; Thulke, HH; Van der Werf, W; Vicent, A; Yuen, J; Zappala, L; Boberg, J; Jeger, M; Pautasso, M; Dehnen-Schmutz, K | 2018 | Pest categorisation of Melampsora farlowii | EFSA JOURNAL |
| Bragard, C; Di Serio, F; Gonthier, P; Jacques, MA; Miret, JAJ; Justesen, AF; MacLeod, A; Magnusson, CS; Milonas, P; Navas-Cortes, JA; Parnell, S; Potting, R; Reignault, PL; Thulke, HH; Van der Werf, W; Vicent, A; Yuen, J; Zappala, L; Boberg, J; Pautasso, M; Dehnen-Schmutz, K | 2018 | Pest categorisation of Arceuthobium spp. (non-EU) | EFSA JOURNAL |
| Brauners, I; Bruna, L; Gaitnieks, T | 2014 | TESTING THE 'ROTSTOP' BIOLOGICAL PREPARATION FOR CONTROLLING HETEROBASIDION ROOT ROT IN LATVIA | RESEARCH FOR RURAL DEVELOPMENT 2014, VOL 2 |
| Brodde, L; Adamson, K; Camarero, JJ; Castano, C; Drenkhan, R; Lehtijarvi, A; Luchi, N; Migliorini, D; Sanchez-Miranda, A; Stenlid, J; Ozdag, S; Oliva, J | 2019 | Diplodia Tip Blight on Its Way to the North: Drivers of Disease Emergence in Northern Europe | FRONTIERS IN PLANT SCIENCE |
| Brunette, M; Holecy, J; Sedliak, M; Tucek, J; Hanewinkel, M | 2015 | An actuarial model of forest insurance against multiple natural hazards in fir (Abies Alba Mill.) stands in Slovakia | FOREST POLICY AND ECONOMICS |
| Bryner, SF; Prospero, S; Rigling, D | 2014 | Dynamics of Cryphonectria hypovirus Infection in Chestnut Blight Cankers | PHYTOPATHOLOGY |
| Bugmann, H; Lindner, M; Lasch, P; Flechsig, M; Ebert, B; Cramer, W | 2000 | Scaling issues in forest succession modelling | CLIMATIC CHANGE |
| Burdulanyuk, AO; Tatarynova, VI; Vlasenko, VA; Demenko, VM; Rozhkova, TO; Bakumenko, OM | 2018 | Dynamics of the number of bark beetles in the ecosystems of Polissya coniferous forests (Sumy oblast, Ukraine) | UKRAINIAN JOURNAL OF ECOLOGY |
| BYLUND, H; TENOW, O | 1994 | LONG-TERM DYNAMICS OF LEAF MINERS, ERIOCRANIA SPP, ON | ECOLOGICAL ENTOMOLOGY |

| | | | |
|---|---|---|---|
| | | MOUNTAIN BIRCH - ALTERNATE YEAR FLUCTUATIONS AND INTERACTION WITH EPIRRITA-AUTUMNATA | |
| Bystricky, V; Moravcova, J; Polensky, J; Pecenka, J | 2017 | LAND USE CHANGES IN THE LAST HALF CENTURY AND THEIR IMPACT ON WATER RETENTION IN THE SUMAVA MOUNTAINS AND FOOTHILLS (CZECH REPUBLIC) | EUROPEAN COUNTRYSIDE |
| Caccianiga, M; Payette, S; Filion, L | 2008 | Biotic disturbance in expanding subarctic forests along the eastern coast of Hudson Bay | NEW PHYTOLOGIST |
| Cakir, G; Sivrikaya, F; Terzioglu, S; Keles, S; Baskent, EZ | 2007 | Monitoring thirty years of land cover change: Secondary forest succession in the Artvin Forest Planning Unit of Northeastern Turkey | SCOTTISH GEOGRAPHICAL JOURNAL |
| Campioli, M; Vincke, C; Jonard, M; Kint, V; Demaree, G; Ponette, Q | 2012 | Current status and predicted impact of climate change on forest production and biogeochemistry in the temperate oceanic European zone: review and prospects for Belgium as a case study | JOURNAL OF FOREST RESEARCH |
| Catala, S; Perez-Sierra, A; Abad-Campos, P | 2015 | The Use of Genus-Specific Amplicon Pyrosequencing to Assess Phytophthora Species Diversity Using eDNA from Soil and Water in Northern Spain | PLOS ONE |
| Catry, FX; Branco, M; Sousa, E; Caetano, J; Naves, P; Nobrege, F | 2017 | Presence and dynamics of ambrosia beetles and other xylophagous insects in a Mediterranean cork oak forest following fire | FOREST ECOLOGY AND MANAGEMENT |
| Cavaletto, G; Faccoli, M; Marini, L; Martinez-Sanudo, I; Mazzon, L | 2018 | Oviposition site preference of Barbitistes vicetinus (Orthoptera, Tettigoniidae) during outbreaks | AGRICULTURAL AND FOREST ENTOMOLOGY |

| | | | |
|---|---|---|---|
| Cayuela, L; Hodar, JA; Zamora, R | 2011 | Is insecticide spraying a viable and cost-efficient management practice to control pine processionary moth in Mediterranean woodlands? | FOREST ECOLOGY AND MANAGEMENT |
| Cerny, K; Peskova, V; Modlinger, R | 2015 | DISTRIBUTION OF PHYTOPHTHORA DISEASE OF ALDERS IN FOREST STANDS IN THE CZECH REPUBLIC - PRELIMINARY FINDINGS | REPORTS OF FORESTRY RESEARCH-ZPRAVY LESNICKEHO VYZKUMU |
| Cerny, K; Peskova, V; Soukup, F; Havrdova, L; Strnadova, V; Zahradnik, D; Hrabetova, M | 2016 | Gemmamyces bud blight of Picea pungens: a sudden disease outbreak in Central Europe | PLANT PATHOLOGY |
| Chao, W; Kolski-Andreaco, A | 2014 | May 2014: This Month in JoVE - Bioengineering Limb Prostheses, Protecting Trees from Fungal Pathogens, and Investigating the Kinematics of Swallowing | JOVE-JOURNAL OF VISUALIZED EXPERIMENTS |
| Charbonnier, YM; Barbaro, L; Barnagaud, JY; Ampoorter, E; Nezan, J; Verheyen, K; Jactel, H | 2016 | Bat and bird diversity along independent gradients of latitude and tree composition in European forests | OECOLOGIA |
| Chen, L; Huang, JG; Dawson, A; Zhai, LH; Stadt, KJ; Comeau, PG; Whitehouse, C | 2018 | Contributions of insects and droughts to growth decline of trembling aspen mixed boreal forest of western Canada | GLOBAL CHANGE BIOLOGY |
| Chen, ZQ; Lunden, K; Karlsson, B; Vos, I; Olson, A; Lundqvist, SO; Stenlid, J; Wu, HX; Gil, MRG; Elfstrand, M | 2018 | Early selection for resistance to Heterobasidion parviporum in Norway spruce is not likely to adversely affect growth and wood quality traits in late-age performance | EUROPEAN JOURNAL OF FOREST RESEARCH |
| Christensen, TR; Johansson, T; Olsrud, M; Strom, L; Lindroth, A; Mastepanov, M; Malmer, N; Friborg, T; Crill, P; Callaghan, TV | 2007 | A catchment-scale carbon and greenhouse gas budget of a subarctic landscape | PHILOSOPHICAL TRANSACTIONS OF THE ROYAL SOCIETY A-MATHEMATICAL PHYSICAL AND ENGINEERING SCIENCES |
| Chrysopolitou, V; Apostolakis, A; Avtzis, D; Avtzis, N; Diamandis, S; Kemitzoglou, D; Papadimos, D; Perlerou, C; Tsiaoussi, V; Dafis, S | 2013 | Studies on forest health and vegetation changes in Greece under the | BIODIVERSITY AND CONSERVATION |

| | | | |
|---|---|---|---|
| | | effects of climate changes | |
| Chumak, V; Obrist, MK; Moretti, M; Duelli, P | 2015 | Arthropod diversity in pristine vs. managed beech forests in Transcarpathia (Western Ukraine) | GLOBAL ECOLOGY AND CONSERVATION |
| Cienciala, E; Tumajer, J; Zatloukal, V; Beranova, J; Hola, S; Hunova, I; Russ, R | 2017 | Recent spruce decline with biotic pathogen infestation as a result of interacting climate, deposition and soil variables | EUROPEAN JOURNAL OF FOREST RESEARCH |
| Ciesielski, M; Balazy, R; Hycza, T; Dmyterko, E; Bruchwald, A | 2016 | Estimating the damage caused by the wind in the forest stands using satellite imagery and data from the State Forests Information System | SYLWAN |
| Cirovic, D; Stamenkovic, S | 2018 | MAMMAL FAUNA OF SERBIA - VALORISATION OF FUNCTIONAL ROLE AND SPECIES IMPORTANCE IN ECOSYSTEMS | ECOLOGICAL AND ECONOMIC SIGNIFICANCE OF FAUNA OF SERBIA |
| Clouet, M | 2003 | Bill size and breeding periode of pine forest crossbills | REVUE D ECOLOGIE-LA TERRE ET LA VIE |
| Coban, HO; Ozcelik, R; Avci, M | 2014 | Monitoring of damage from cedar shoot moth Dichelia cedricola Diakonoff (Lep.: Tortricidae) by multi-temporal Landsat imagery | IFOREST-BIOGEOSCIENCES AND FORESTRY |
| Conedera, M; Colombaroli, D; Tinner, W; Krebs, P; Whitlock, C | 2017 | Insights about past forest dynamics as a tool for present and future forest management in Switzerland | FOREST ECOLOGY AND MANAGEMENT |
| Contarini, M; Ruiu, L; Pilarska, D; Luciano, P | 2015 | POTENTIAL IMPACT OF ENTOMOPHAGA MAIMAIGA HUMBER, SHIMAZU, AND SOPER (ENTOMOPHTHORALES ENTOMOPHTHORACEAE) ON THE LEPIDOPTERAN FAUNA INHABITING CORK FORESTS IN SARDINIA (ITALY) | REDIA-GIORNALE DI ZOOLOGIA |

| | | | |
|---|---|---|---|
| Cosson, JF | 2019 | Ecology of Lyme disease | SANTE PUBLIQUE |
| Cottrell, S; Mattor, KM; Morris, JL; Fettig, CJ; McGrady, P; Maguire, D; James, PMA; Clear, J; Wurtzebach, Z; Wei, Y; B[illegible] R | | | |
| Crane, PE | 2001 | Morphology, taxonomy, and nomenclature of the Chrysomyxa ledi complex and related rust fungi on spruce and Ericaceae in North America and Europe | CANADIAN JOURNAL OF BOTANY-REVUE CANADIENNE DE BOTANIQUE |
| Csoka, G | 1997 | Increased insect damage in Hungarian forests under drought impact | BIOLOGIA |
| Cuellar, AC; Kjaer, LJ; Baum, A; Stockmarr, A; Skovgard, H; Nielsen, SA; Andersson, MG; Lindstrom, A; Chirico, J; Luhken, R; Steinke, S; Kiel, E; Gethmann, J; Conraths, FJ; Larska, M; Smreczak, M; Orlowska, A; Hamnes, I; Sviland, S; Hopp, P; Brugger, K; Rubel, F; Balenghien, T; Garros, C; Rakotoarivony, I; Allene, X; Lhoir, J; Chavernac, D; Delecolle, JC; Mathieu, B; Delecolle, D; Setier-Rio, ML; Venail, R; Scheid, B; Chueca, MAM; Barcelo, C; Lucientes, J; Estrada, R; Mathis, A; Tack, W; Bodker, R | 2018 | Monthly variation in the probability of presence of adult Culicoides populations in nine European countries and the implications for targeted surveillance | PARASITES & VECTORS |
| Cunze, S; Kochmann, J; Kuhn, T; Frank, R; Dorge, DD; Klimpel, S | 2018 | Spatial and temporal patterns of human Puumala virus (PUUV) infections in Germany | PEERJ |
| Dadasoglu, F; Tozlu, G; Kotan, R; Gokturk, T; Karagoz, K | 2016 | Biological Control of Pine Sawfly (Diprion pini L.) and Molecular Characterisation of Effective Strains | ROMANIAN BIOTECHNOLOGICAL LETTERS |
| Dale, VH; Kline, KL; Parish, ES; Cowie, AL; Emory, R; Malmsheimer, RW; Slade, R; Smith, CT; Wigley, TB; Bentsen, NS; Berndes, G; Bernier, P; | 2017 | Status and prospects for renewable energy using wood pellets from the | GLOBAL CHANGE BIOLOGY BIOENERGY |

| | | | |
|---|---|---|---|
| Brandao, M; Chum, HL; Diaz-Chavez, R; Egnell, G; Gustavsson, L; Schweinle, J; Stupak, I; Trianosky, P; Walter, A; Whittaker, C; Brown, M; Chescheir, G; Dimitriou, I; Donnison, C; Eng, AG; Hoyt, KP; Jenkins, JC; Johnson, K; Levesque, CA; Lockhart, V; Negri, MC; Nettles, JE; Wellisch, M | | southeastern United States | |
| David, G; Giffard, B; Piou, D; Jactel, H | 2014 | Dispersal capacity of Monochamus galloprovincialis, the European vector of the pine wood nematode, on flight mills | JOURNAL OF APPLIED ENTOMOLOGY |
| Day, KR | 1997 | The influence of temperature on egg mortality in the budmoth Zeiraphera diniana (Lepidoptera:Tortricidae), and its role in determining the regional abundance of an important forest pest | BULLETIN OF ENTOMOLOGICAL RESEARCH |
| de Dios, VR; Fischer, C; Colinas, C | 2007 | Climate change effects on mediterranean forests and preventive measures | NEW FORESTS |
| De Dobbelaere, I; Vercauteren, A; Speybroeck, N; Berkvens, D; Van Bockstaele, E; Maes, M; Heungens, K | 2010 | Effect of host factors on the susceptibility of Rhododendron to Phytophthora ramorum | PLANT PATHOLOGY |
| de Groot, M; Diaci, J; Ogris, N | 2019 | Forest management history is an important factor in bark beetle outbreaks: Lessons for the future | FOREST ECOLOGY AND MANAGEMENT |
| de Groot, M; Ogris, N; Kobler, A | 2018 | The effects of a large-scale ice storm event on the drivers of bark beetle outbreaks and associated management practices | FOREST ECOLOGY AND MANAGEMENT |
| de Queiroz, DL; Majer, J; Burckhardt, D; Zanetti, R; Fernandez, JIR; de Queiroz, EC; Garrastazu, M; Fernandes, BV; dos Anjos, N | 2013 | Predicting the geographical distribution of Glycaspis brimblecombei (Hemiptera: Psylloidea) in Brazil | AUSTRALIAN JOURNAL OF ENTOMOLOGY |
| De Somviele, B; Lyytikainen-Saarenmaa, P; Niemela, P | 2007 | Stand edge effects on distribution and condition of Diprionid sawflies | AGRICULTURAL AND FOREST ENTOMOLOGY |

| | | | |
|---|---|---|---|
| Delattre, P; De Sousa, B; Fichet-Calvet, E; Quere, JP; Giraudoux, P | 1999 | Vole outbreaks in a landscape context: evidence from a six year study of Microtus arvalis | LANDSCAPE ECOLOGY |
| DENBOER, PJ; SZYSZKO, J; VERMEULEN, R | 1993 | SPREADING THE RISK OF EXTINCTION BY GENETIC DIVERSITY IN POPULATIONS OF THE CARABID BEETLE PTEROSTICHUS-OBLONGOPUNCTATUS F (COLEOPTERA, CARABIDAE) | NETHERLANDS JOURNAL OF ZOOLOGY |
| Denes, AL; Kolcsar, LP; Torok, E; Keresztes, L | 2016 | Phylogeography of the micro-endemic Pedicia staryi group (Insecta: Diptera): evidence of relict biodiversity in the Carpathians | BIOLOGICAL JOURNAL OF THE LINNEAN SOCIETY |
| Desprez-Loustau, ML; Courtecuisse, R; Robin, C; Husson, C; Moreau, PA; Blancard, D; Selosse, MA; Lung-Escarmant, B; Piou, D; Sache, I | 2010 | Species diversity and drivers of spread of alien fungi (sensu lato) in Europe with a particular focus on France | BIOLOGICAL INVASIONS |
| Diamandis, S; Perlerou, C | 2005 | The role of Spulerina simploniella in the spread of chestnut blight | FOREST PATHOLOGY |
| Dmyterko, E; Bruchwald, A | 2018 | Changes in the forests of the Bieszczady Mts. | SYLWAN |
| Drekic, M; Pajnik, LP; Vasic, V; Pap, P; Pilipovic, A | 2014 | CONTRIBUTION TO THE STUDY OF BIOLOGY OF ASH WEEVIL (Stereonychus fraxini De Geer) | SUMARSKI LIST |
| Drenkhan, R; Tomesova-Haataja, V; Fraser, S; Bradshaw, RE; Vahalik, P; Mullett, MS; Martin-Garcia, J; Bulman, LS; Wingfield, MJ; Kirisits, T; Cech, TL; Schmitz, S; Baden, R; Tubby, K; Brown, A; Georgieva, M; Woods, A; Ahumada, R; Jankovsky, L; Thomsen, IM; Adamson, K; Marcais, B; Vuorinen, M; Tsopelas, P; Koltay, A; Halasz, A; La Porta, N; Anselmi, N; Kiesnere, R; Markovskaja, S; Kacergius, A; Papazova-Anakieva, I; Risteski, M; Sotirovski, K; Lazarevic, J; Solheim, H; Boron, P; Braganca, H; Chira, D; Musolin, DL; Selikhovkin, AV; Bulgakov, TS; Keca, N; Karadzic, D; Galovic, V; Pap, P; Markovic, M; | 2016 | Global geographic distribution and host range of Dothistroma species: a comprehensive review | FOREST PATHOLOGY |

| | | | |
|---|---|---|---|
| Pajnik, LP; Vasic, V; Ondruskova, E; Piskur, B; Sadikovic, D; Diez, JJ; Solla, A; Millberg, H; Stenlid, J; Angst, A; Queloz, V; Lehtijarvi, A; Dogmus-Lehtijarvi, HT; Oskay, F; Davydenko, K; Meshkova, V; Craig, D; Woodward, S; Barnes, I | | | |
| Ducic, V; Lukovic, J; Milenkovic, M; Curcic, N | 2012 | NORTH ATLANTIC OSCILLATION (NAO) AND INSECT DAMAGE IN SERBIAN FORESTS | ARCHIVES OF BIOLOGICAL SCIENCES |
| Dvorak, M; Janos, P; Botella, L; Rotkova, G; Zas, R | 2017 | Spore Dispersal Patterns of Fusarium circinatum on an Infested Monterey Pine Forest in North-Western Spain | FORESTS |
| Dworschak, K; Meyer, D; Gruppe, A; Schopf, R | 2014 | Choice or constraint: Plasticity in overwintering sites of the European spruce bark beetle | FOREST ECOLOGY AND MANAGEMENT |
| Dziegielewska, M; Skwiercz, AT | 2018 | Co-occurrence of entomopathogenic nematodes and tree pests in forest communities of northern Poland | SYLWAN |
| Ebenhard, T; Forsberg, M; Lind, T; Nilsson, D; Andersson, R; Emanuelsson, U; Eriksson, L; Hultaker, O; Wide, MI; Stahl, G | 2017 | Environmental effects of brushwood harvesting for bioenergy | FOREST ECOLOGY AND MANAGEMENT |
| EIDMANN, HH | 1992 | IMPACT OF BARK BEETLES ON FORESTS AND FORESTRY IN SWEDEN | JOURNAL OF APPLIED ENTOMOLOGY-ZEITSCHRIFT FUR ANGEWANDTE ENTOMOLOGIE |
| Elliot, M; Schlenzig, A; Harris, CM; Meagher, TR; Green, S | 2015 | An improved method for qPCR detection of three Phytophthora spp. in forest and woodland soils in northern Britain | FOREST PATHOLOGY |
| Erturk, U; Akca, Y | 2014 | Overview of Walnut Culture in Turkey | VII INTERNATIONAL WALNUT SYMPOSIUM |
| Euser, SM; Nagelkerke, NJ; Schuren, F; Jansen, R; Den Boer, JW | 2012 | Genome Analysis of Legionella pneumophila Strains Using a Mixed-Genome Microarray | PLOS ONE |
| Faccoli, M; Bernardinelli, I | 2014 | Composition and Elevation of Spruce Forests Affect Susceptibility to Bark Beetle Attacks: | FORESTS |

| | | | |
|---|---|---|---|
| | | Implications for Forest Management | |
| Faccoli, M; Finozzi, V; Colombari, F | 2012 | Effectiveness of different trapping protocols for outbreak management of the engraver pine beetle Ips acuminatus (Curculionidae, Scolytinae) | INTERNATIONAL JOURNAL OF PEST MANAGEMENT |
| Faccoli, M; Stergulc, F | 2008 | Damage reduction and performance of mass trapping devices for forest protection against the spruce bark beetle, Ips typographus (Coleoptera Curculionidae Scolytinae) | ANNALS OF FOREST SCIENCE |
| Faccoli, M; Stergulc, F | 2006 | A practical method for predicting the short-time trend of bivoltine populations of Ips typographus (L.) (Col., Scolytidae) | JOURNAL OF APPLIED ENTOMOLOGY |
| Faccoli, M; Stergulc, F | 2004 | Ips typographus (L.) pheromone trapping in south Alps: spring catches determine damage thresholds | JOURNAL OF APPLIED ENTOMOLOGY |
| Falt-Nardmann, J; Klemola, T; Roth, M; Ruohomaki, K; Saikkonen, K | 2016 | Northern geometrid forest pests (Lepidoptera: Geometridae) hatch at lower temperatures than their southern conspecifics: Implications of climate change | EUROPEAN JOURNAL OF ENTOMOLOGY |
| Falt-Nardmann, JJJ; Tikkanen, OP; Ruohomaki, K; Otto, LF; Leinonen, R; Poyry, J; Saikkonen, K; Neuvonen, S | 2018 | The recent northward expansion of Lymantria monacha in relation to realised changes in temperatures of different seasons | FOREST ECOLOGY AND MANAGEMENT |
| Fassnacht, FE; Latifi, H; Ghosh, A; Joshi, PK; Koch, B | 2014 | Assessing the potential of hyperspectral imagery to map bark beetle-induced tree mortality | REMOTE SENSING OF ENVIRONMENT |
| Fernandez-Fernandez, M; Naves, P; Musolin, DL; Selikhovkin, AV; Cleary, M; Chira, D; Paraschiv, M; Gordon, T; Solla, A; Papazova-Anakieva, I; Drenkhan, T; Georgieva, M; Altunisik, | 2019 | Pine Pitch Canker and Insects: Regional Risks, Environmental Regulation, and Practical Management Options | FORESTS |

| | | | |
|---|---|---|---|
| A; Morales-Rodriguez, C; Tabakovic-Tosic, M; Avtzis, DN; Georgiev, G; Doychev, DD; Nacheski, S; Trestic, T; Elvira-Recuenco, M; Diez, JJ; Witzell, J | | | |
| Fernandez-Fernandez, M; Naves, P; Witzell, J; Musolin, DL; Selikhovkin, AV; Paraschiv, M; Chira, D; Martinez-Alvarez, P; Martin-Garcia, J; Munoz-Adalia, EJ; Altunisik, A; Cocuzza, GEM; Di Silvestro, S; Zamora, C; Diez, JJ | 2019 | Pine Pitch Canker and Insects: Relationships and Implications for Disease Spread in Europe | FORESTS |
| Fiedler, K; Truxa, C | 2012 | Species richness measures fail in resolving diversity patterns of speciose forest moth assemblages | BIODIVERSITY AND CONSERVATION |
| Fieseler, K; Heiermann, J; Schutz, S | 2012 | Is climate change going to affect the diversity of nocturnal Macrolepidoptera in the Solling-mountains? | MITTEILUNGEN DER DEUTSCHEN GESELLSCHAFT FUR ALLGEMEINE UND ANGEWANDTE ENTOMOLOGIE, BD 18 |
| Fischer, A; Lindner, M; Abs, C; Lasch, P | 2002 | Vegetation dynamics in central European forest ecosystems (near-natural as well as managed) after storm events | FOLIA GEOBOTANICA |
| Fonder, W | 2007 | Stand conversion for sustainable forestry | Quo Vadis, Forestry?, Proceedings |
| Forgach, P; Boncz, A; Erdelyi, K; Lorincz, M; Molnar, B; Zentai, J; Szucs, G; Reuter, G; Bakonyi, T | 2010 | Hepatitis E virus - literature review and situation in Hungary from veterinary point of view | MAGYAR ALLATORVOSOK LAPJA |
| Forgach, P; Nowotny, N; Erdelyi, K; Boncz, A; Zentai, J; Szucs, G; Reuter, G; Bakonyi, T | 2010 | Detection of Hepatitis E virus in samples of animal origin collected in Hungary | VETERINARY MICROBIOLOGY |
| Francis, A; Darbyshire, SJ; Clements, DR; DiTommaso, A | 2011 | The Biology of Canadian Weeds. 146. Lapsana communis L. | CANADIAN JOURNAL OF PLANT SCIENCE |
| Frigimelica, G; Faccoli, M | 1999 | Preliminary report on the occurrence of Cryphonectria parasitica (Murrill) Barr on different tree species in Friuli Venezia-Giulia (Italy) | SECOND INTERNATIONAL SYMPOSIUM ON CHESTNUT |
| Galinski, W; Witowski, J | 1996 | The carbon pulse resulting from forest dieback related to insect outbreaks: Case study of a forest district in the Sudety mountains (southwest Poland) | FOREST ECOSYSTEMS, FOREST MANAGEMENT AND THE GLOBAL CARBON CYCLE |

| | | | |
|---|---|---|---|
| Ganley, RJ; Watt, MS; Manning, L; Iturritxa, E | 2009 | A global climatic risk assessment of pitch canker disease | CANADIAN JOURNAL OF FOREST RESEARCH |
| Garbelotto, M; Gonthier, P | 2013 | Biology, Epidemiology, and Control of Heterobasidion Species Worldwide | ANNUAL REVIEW OF PHYTOPATHOLOGY, VOL 51 |
| Garcia, D; Minarro, M; Martinez-Sastre, R | 2018 | Birds as suppliers of pest control in cider apple orchards: Avian biodiversity drivers and insectivory effect | AGRICULTURE ECOSYSTEMS & ENVIRONMENT |
| Gencer, NS; Mert, C | 2019 | Studies on the Gall Characteristics of Dryocosmus kuriphilus in Chestnut Genotypes in Yalova and Bursa Provinces of Turkey | NOTULAE BOTANICAE HORTI AGROBOTANICI CLUJ-NAPOCA |
| Ghimire, RP; Kivimaenpaa, M; Blomqvist, M; Holopainen, T; Lyytikainen-Saarenmaa, P; Holopainen, JK | 2016 | Effect of bark beetle (Ips typographus L.) attack on bark VOC emissions of Norway spruce (Picea abies Karst.) trees | ATMOSPHERIC ENVIRONMENT |
| Gibbs, JN | 2001 | Vascular wilt diseases of trees - An Anglo-American perspective | SHADE TREE: WILT DISEASES |
| Giongo, S; Longa, CMO; Dal Maso, E; Montecchio, L; Maresi, G | 2017 | Evaluating the impact of Hymenoscyphus fraxineus in Trentino (Alps, Northern Italy): first investigations | IFOREST-BIOGEOSCIENCES AND FORESTRY |
| Giordano, L; Garbelotto, M; Nicolotti, G; Gonthier, P | 2013 | Characterization of fungal communities associated with the bark beetle Ips typographus varies depending on detection method, location, and beetle population levels | MYCOLOGICAL PROGRESS |
| Glavendekic, MM; Medarevic, MJ | 2010 | INSECT DEFOLIATORS AND THEIR INFLUENCE ON OAK FORESTS IN THE DJERDAP NATIONAL PARK, SERBIA | ARCHIVES OF BIOLOGICAL SCIENCES |
| Glowacka, B; Bystrowski, C; Skrzecz, I | 2018 | Efficacy of Mimic 240 LV in the protection of Scots pine Pinus sylvestris L. against the nun moth Lymantria monacha L. and the pine lappet moth Dendrolimus pini L. | SYLWAN |

| | | | |
|---|---|---|---|
| Gonthier, P; Anselmi, N; Capretti, P; Bussotti, F; Feducci, M; Giordano, L; Honorati, T; Lione, G; Luchi, N; Michelozzi, M; Paparatti, B; Sillo, F; Vettraino, AM; Garbelotto, M | 2014 | An integrated approach to control the introduced forest pathogen Heterobasidion irregulare in Europe | FORESTRY |
| Greco, S; Infusino, M; Scalercio, S | 2017 | Massive capture of Eilema lurideola (Lepidoptera: Erebidae) in a beech forest: outbreak vs dispersal | ENTOMOLOGIA GENERALIS |
| Greenberg, R; Pravosudov, V; Sterling, J; Kozlenko, A; Kontorschikov, V | 1999 | Divergence in foraging behavior of foliage-gleaning birds of Canadian and Russian boreal forests | OECOLOGIA |
| Gren, IM; Aklilu, AZ; Elofsson, K | 2018 | Forest Carbon Sequestration, Pathogens and the Costs of the EU's 2050 Climate Targets | FORESTS |
| Grodzinska, K; Szarek-Lukaszewska, G | 1997 | Section 4: Evaluation of forest health - Regional perspective - Polish mountain forests: Past, present and future | ENVIRONMENTAL POLLUTION |
| Grodzki, W | 2004 | Some reactions of Ips typographys (L.) (Col.: Scolytidae) to changing breeding conditions in a forest decline area in the Sudeten Mountains, Poland | JOURNAL OF PEST SCIENCE |
| Grodzki, W | 1997 | Pityogenes chalcographus (Coleoptera, Scolytidae) - An indicator of man-made changes in Norway spruce stands | BIOLOGIA |
| Grodzki, W | 1997 | Changes in the occurrence of bark beetles on Norway spruce in a forest decline area in the Sudety Mountains in Poland | INTEGRATING CULTURAL TACTICS INTO THE MANAGEMENT OF BARK BEETLE AND REFORESTATION PESTS, PROCEEDINGS |
| Grodzki, W; Jakus, R; Lajzova, E; Sitkova, Z; Maczka, T; Skvarenina, J | 2006 | Effects of intensive versus no management strategies during an outbreak of the bark beetle Ips typographus (L.) (Col.: Curculionidae, Scolytinae) in the Tatra | ANNALS OF FOREST SCIENCE |

| | | | |
|---|---|---|---|
| | | Mts. in Poland and Slovakia | |
| Groenen, F; Meurisse, N | 2012 | Historical distribution of the oak processionary moth Thaumetopoea processionea in Europe suggests recolonization instead of expansion | AGRICULTURAL AND FOREST ENTOMOLOGY |
| Gryczynska, A; Kowalec, M | 2019 | Different Competence as a Lyme Borreliosis Causative Agent Reservoir Found in Two Thrush Species: The Blackbird (Turdus merula) and the Song Thrush (Turdus philomelos) | VECTOR-BORNE AND ZOONOTIC DISEASES |
| Gryczynska, A; Zgodka, A; Ploski, R; Siemiathkowski, M | 2004 | Borrelia burgdorferi sensu lato infection in passerine birds from the Mazurian Lake region (Northeastern Poland) | AVIAN PATHOLOGY |
| Hanewinkel, M; Breidenbach, J; Neeff, T; Kublin, E | 2008 | Seventy-seven years of natural disturbances in a mountain forest area - the influence of storm, snow, and insect damage analysed with a long-term time series | CANADIAN JOURNAL OF FOREST RESEARCH-REVUE CANADIENNE DE RECHERCHE FORESTIERE |
| HANSEN, EM | 1995 | DOUGLAS-FIR TUSSOCK MOTH (ORGYIA-PSEUDOTSUGATA MCDUNNOUGH) ON SUB-ALPINE FIR IN NORTHERN UTAH | GREAT BASIN NATURALIST |
| Hansen, NM; Ims, RA; Hagen, SB | 2009 | No Impact of Pupal Predation on the Altitudinal Distribution of Autumnal Moth and Winter Moth (Lepidoptera: Geometridae) in Sub-Arctic Birch Forest | ENVIRONMENTAL ENTOMOLOGY |
| Hanssen, KH; Solberg, S | 2007 | Assessment of defoliation during a pine sawfly outbreak: Calibration of airborne laser scanning data with hemispherical photography | FOREST ECOLOGY AND MANAGEMENT |

| | | | |
|---|---|---|---|
| Haque, MM; Diez, JJ | 2012 | Susceptibility of common alder (Alnus glutinosa) seeds and seedlings to Phytophthora alni and other Phytophthora species | FOREST SYSTEMS |
| Harris, AR; Webber, JF | 2016 | Sporulation potential, symptom expression and detection of Phytophthora ramorum on larch needles and other foliar hosts | PLANT PATHOLOGY |
| Hartl-Meier, C; Esper, J; Liebhold, A; Konter, O; Rothe, A; Buntgen, U | 2017 | Effects of host abundance on larch budmoth outbreaks in the European Alps | AGRICULTURAL AND FOREST ENTOMOLOGY |
| Havasova, M; Ferencik, J; Jakus, R | 2017 | Interactions between windthrow, bark beetles and forest management in the Tatra national parks | FOREST ECOLOGY AND MANAGEMENT |
| Haynes, KJ; Allstadt, AJ; Klimetzek, D | 2014 | Forest defoliator outbreaks under climate change: effects on the frequency and severity of outbreaks of five pine insect pests | GLOBAL CHANGE BIOLOGY |
| Heiermann, J; Schutz, S | 2008 | The effect of the tree species ratio of European beech (Fagus sylvatica L.) and Norway spruce (Picea abies (L.) Karst.) on polyphagous and monophagous pest species - Lymantria monacha L. and Calliteara pudibunda L. (Lepidoptera : Lymantriidae) as an example | FOREST ECOLOGY AND MANAGEMENT |
| Heim, O; Treitler, JT; Tschapka, M; Knornschild, M; Jung, K | 2015 | The Importance of Landscape Elements for Bat Activity and Species Richness in Agricultural Areas | PLOS ONE |
| Heisswolf, A; Kaar, M; Klemola, T; Ruohomaki, K | 2010 | Local outbreaks of Operophtera brumata and Operophtera fagata cannot be explained by low vulnerability to pupal predation | AGRICULTURAL AND FOREST ENTOMOLOGY |

| | | | |
|---|---|---|---|
| Hejcman, M; Hejcmanova, P; Pavlu, V; Benes, J | 2013 | Origin and history of grasslands in Central Europe - a review | GRASS AND FORAGE SCIENCE |
| HelderJose; DeAndrade, HK | 1997 | Food and feeding habits of the neotropical river otter Lontra longicaudis (Carnivora, Mustelidae) | MAMMALIA |
| Heliasz, M; Johansson, T; Lindroth, A; Molder, M; Mastepanov, M; Friborg, T; Callaghan, TV; Christensen, TR | 2011 | Quantification of C uptake in subarctic birch forest after setback by an extreme insect outbreak | GEOPHYSICAL RESEARCH LETTERS |
| Helmens, KF; Valiranta, M; Engels, S; Shala, S | 2012 | Large shifts in vegetation and climate during the Early Weichselian (MIS 5d-c) inferred from multi-proxy evidence at Sokli (northern Finland) | QUATERNARY SCIENCE REVIEWS |
| Henin, JM; Huart, O; Rondeux, J | 2003 | Biogeographical observations on four scolytids (Coleoptera, Scolytidae) and one lymexylonid (Coleoptera, Lymexylonidae) in Wallonia (Southern Belgium) | BELGIAN JOURNAL OF ZOOLOGY |
| Hensel, A; Neubauer, H | 2002 | Human pathogens associated with on-farm practices - Implications for control and surveillance strategies | FOOD SAFETY ASSURANCE IN THE PRE-HARVEST PHASE, VOL 1: FOOD SAFETY ASSURANCE AND VETERINARY PUBLIC HEALTH |
| Hentschel, R; Moller, K; Wenning, A; Degenhardt, A; Schroder, J | 2018 | Importance of Ecological Variables in Explaining Population Dynamics of Three Important Pine Pest Insects | FRONTIERS IN PLANT SCIENCE |
| Hielscher, K | 2014 | Surveillance of nun moth (Lymantria monacha L., Lepidoptera: Lymantriidae) flight activity in the federal state of Brandenburg - Are there rationalization possibilities? | MITTEILUNGEN DER DEUTSCHEN GESELLSCHAFT FUR ALLGEMEINE UND ANGEWANDTE ENTOMOLOGIE, BD 19 |
| Hill, L; Hemery, G; Hector, A; Brown, N | 2019 | Maintaining ecosystem properties after loss of ash in Great Britain | JOURNAL OF APPLIED ECOLOGY |
| Hilszczanski, J; Jaworski, T | 2018 | Biodiversity conservation in the Bialowieza Forest in the context of natural and anthropogenic disturbances dynamics | SYLWAN |

| | | | |
|---|---|---|---|
| Hilszczanski, J; Jaworski, T; Plewa, R; Horak, J | 2016 | Tree species and position matter: the role of pests for survival of other insects | AGRICULTURAL AND FOREST ENTOMOLOGY |
| Hittenbeck, A; Bialozyt, R; Schmidt, M | 2019 | Modelling the population fluctuation of winter moth and mottled umber moth in central and northern Germany | FOREST ECOSYSTEMS |
| Hlasny, T; Trombik, J; Holusa, J; Lukasova, K; Grendar, M; Turcani, M; Zubrik, M; Tabakovic-Tosic, M; Hirka, A; Buksha, I; Modlinger, R; Kacprzyk, M; Csoka, G | 2016 | Multi-decade patterns of gypsy moth fluctuations in the Carpathian Mountains and options for outbreak forecasting | JOURNAL OF PEST SCIENCE |
| Hlasny, T; Turcani, M | 2009 | Insect Pests as Climate Change Driven Disturbances in Forest Ecosystems | BIOCLIMATOLOGY AND NATURAL HAZARDS |
| Hodar, JA; Castro, J; Zamora, R | 2003 | Pine processionary caterpillar Thaumetopoea pityocampa as a new threat for relict Mediterranean Scots pine forests under climatic warming | BIOLOGICAL CONSERVATION |
| Hodar, JA; Zamora, R | 2004 | Herbivory and climatic warming: a Mediterranean outbreaking caterpillar attacks a relict, boreal pine species | BIODIVERSITY AND CONSERVATION |
| Hodar, JA; Zamora, R; Cayuela, L | 2012 | Climate change and the incidence of a forest pest in Mediterranean ecosystems: can the North Atlantic Oscillation be used as a predictor? | CLIMATIC CHANGE |
| Hof, AR; Svahlin, A | 2016 | The potential effect of climate change on the geographical distribution of insect pest species in the Swedish boreal forest | SCANDINAVIAN JOURNAL OF FOREST RESEARCH |
| Hof, AR; Svahlin, A | 2016 | Not erroneous but cautious conclusions about the potential effect of climate change on the geographical distribution of insect pest species in the Swedish | SCANDINAVIAN JOURNAL OF FOREST RESEARCH |

| | | | |
|---|---|---|---|
| | | boreal forest. Response to Bjorklund et al. (2015) | |
| HOLAH, JC; WILSON, MV; HANSEN, EM | 1993 | EFFECTS OF A NATIVE FOREST PATHOGEN, PHELLINUS-WEIRII, ON DOUGLAS-FIR FOREST COMPOSITION IN WESTERN OREGON | CANADIAN JOURNAL OF FOREST RESEARCH-REVUE CANADIENNE DE RECHERCHE FORESTIERE |
| Holmer, L; Stenlid, J | 1997 | Resinicium bicolor; A potential biological control agent for Heterobasidion annosum | EUROPEAN JOURNAL OF FOREST PATHOLOGY |
| Holusa, J; Weiser, J; Drapela, K | 2007 | Pathogens of Ips duplicatus (Coleoptera : Scolytidae) in three areas in central Europe | ACTA PROTOZOOLOGICA |
| Hoshizaki, K; Nakabayashi, Y; Mamiya, Y; Matsushita, M | 2016 | Localized within- and between-tree variation in nematode distribution during latent state of pine wilt disease makes the disease status cryptic | FOREST PATHOLOGY |
| Huang, JB; Kautz, M; Trowbridge, AM; Hammerbacher, A; Raffa, KF; Adams, HD; Goodsman, DW; Xu, CG; Meddens, AJH; Kandasamy, D; Gershenzon, J; Seidl, R; Hartmann, H | 2020 | Tree defence and bark beetles in a drying world: carbon partitioning, functioning and modelling | NEW PHYTOLOGIST |
| Huttunen, L; Blande, JD; Li, T; Rousi, M; Klemola, T | 2013 | Effects of warming climate on early-season carbon allocation and height growth of defoliated mountain birches | PLANT ECOLOGY |
| Hytteborn, H; Svensson, BM; Kempe, K; Press, A; Rydin, H | 2017 | Century-long tree population dynamics in a deciduous forest stand in central Sweden | JOURNAL OF VEGETATION SCIENCE |
| Ibanez-Justicia, A; Cianci, D | 2015 | Modelling the spatial distribution of the nuisance mosquito species Anopheles plumbeus (Diptera: Culicidae) in the Netherlands | PARASITES & VECTORS |
| Inanc, S; Ayaz, H | 2019 | THE ROLE OF LOCAL PEOPLE IN THE FIGHT AGAINST FOREST PESTS: THE CASE OF ARTVIN REGIONAL DIRECTORATE OF FORESTRY | FRESENIUS ENVIRONMENTAL BULLETIN |

| | | | |
|---|---|---|---|
| Inoue, MN; Suzuki-Ohno, Y; Haga, Y; Aarai, H; Sano, T; Martemyanov, VV; Kunimi, Y | 2019 | Population dynamics and geographical distribution of the gypsy moth, Lymantria dispar, in Japan | FOREST ECOLOGY AND MANAGEMENT |
| Irmler, U | 2018 | Which carabid species (Coleoptera: Carabidae) profit from organic farming after a succession of 15 years? | AGRICULTURE ECOSYSTEMS & ENVIRONMENT |
| Jacob, J; Manson, P; Barfknecht, R; Fredricks, T | 2014 | Common vole ( Microtus arvalis) ecology and management: implications for risk assessment of plant protection products | PEST MANAGEMENT SCIENCE |
| Jacob, J; Ulrich, RG; Freise, J; Schmolz, E | 2014 | Monitoring populations of rodent reservoirs of zoonotic diseases. Projects, aims and results | BUNDESGESUNDHEITSBLATT-GESUNDHEITSFORSCHUNG-GESUNDHEITSSCHUTZ |
| Jacquemyn, H; Brys, R; Hutchings, MJ | 2008 | Biological Flora of the British Isles: Paris quadrifolia L. | JOURNAL OF ECOLOGY |
| Jacquet, JS; Bosc, A; O'Grady, AP; Jactel, H | 2013 | Pine growth response to processionary moth defoliation across a 40-year chronosequence | FOREST ECOLOGY AND MANAGEMENT |
| Jacquet, JS; Orazio, C; Jactel, H | 2012 | Defoliation by processionary moth significantly reduces tree growth: a quantitative review | ANNALS OF FOREST SCIENCE |
| Janik, T; Romportl, D | 2018 | Recent land cover change after the Kyrill windstorm in the Sumava NP | APPLIED GEOGRAPHY |
| Janousek, J; Krumbock, S; Kirisits, T; Bradshaw, RE; Barnes, I; Jankovsky, L; Stauffer, C | 2014 | Development of microsatellite and mating type markers for the pine needle pathogen Lecanosticta acicola | AUSTRALASIAN PLANT PATHOLOGY |
| Janousek, J; Wingfield, MJ; Monsivais, JGM; Jankovsky, L; Stauffer, C; Konecny, A; Barnes, I | 2016 | Genetic Analyses Suggest Separate Introductions of the Pine Pathogen Lecanosticta acicola Into Europe | PHYTOPATHOLOGY |
| JARDON, Y; FILION, L; CLOUTIER, C | 1994 | TREE-RING EVIDENCE FOR ENDEMICITY OF THE LARCH SAWFLY IN NORTH-AMERICA | CANADIAN JOURNAL OF FOREST RESEARCH-REVUE CANADIENNE DE RECHERCHE FORESTIERE |
| Jeger, M; Bragard, C; Caffier, D; Candresse, T; Chatzivassiliou, E; | 2017 | Pest categorisation of Entoleucamammata | EFSA JOURNAL |

| | | | |
|---|---|---|---|
| Dehnen-Schmutz, K; Gilioli, G; Gregoire, JC; Jaques Miret, JA; MacLeod, A; Navarro, MN; Niere, B; Parnell, S; Potting, R; Rafoss, T; Rossi, V; Urek, G; Van Bruggen, A; Van der Werf, W; West, J; Winter, S; Boberg, J; Gonthier, P; Pautasso, M | | | |
| Jeger, M; Bragard, C; Caffier, D; Candresse, T; Chatzivassiliou, E; Dehnen-Schmutz, K; Gilioli, G; Gregoire, JC; Miret, JAJ; MacLeod, A; Navarro, MN; Niere, B; Parnell, S; Potting, R; Rafoss, T; Rossi, V; Urek, G; Van Bruggen, A; Van der Werf, W; West, J; Winter, S; Boberg, J; Gonthier, P; Pautasso, M | 2018 | Pest categorisation of Melampsora medusae | EFSA JOURNAL |
| Jeger, M; Bragard, C; Caffier, D; Candresse, T; Chatzivassiliou, E; Dehnen-Schmutz, K; Gilioli, G; Gregoire, JC; Miret, JAJ; MacLeod, A; Navarro, MN; Niere, B; Parnell, S; Potting, R; Rafoss, T; Rossi, V; Urek, G; Van Bruggen, A; Van der Werf, W; West, J; Winter, S; Boberg, J; Gonthier, P; Pautasso, M | 2018 | Pest categorisation of Bretziella fagacearum | EFSA JOURNAL |
| Jeger, M; Bragard, C; Caffier, D; Candresse, T; Chatzivassiliou, E; Dehnen-Schmutz, K; Gilioli, G; Gregoire, JC; Miret, JAJ; MacLeod, A; Navarro, MN; Niere, B; Parnell, S; Potting, R; Rafoss, T; Rossi, V; Urek, G; Van Bruggen, A; Van Der Werf, W; West, J; Winter, S; Boberg, J; Gonthier, P; Pautasso, M | 2017 | Pest categorisation of Davidsoniella virescens | EFSA JOURNAL |
| Jeger, M; Bragard, C; Caffier, D; Candresse, T; Chatzivassiliou, E; Dehnen-Schmutz, K; Gilioli, G; Gregoire, JC; Miret, JAJ; MacLeod, A; Navarro, MN; Niere, B; Parnell, S; Potting, R; Rafoss, T; Rossi, V; Urek, G; Van Bruggen, A; Van Der Werf, W; West, J; Winter, S; Boberg, J; Gonthier, P; Pautasso, M | 2017 | Pest categorisation of Stegophora ulmea | EFSA JOURNAL |
| Jeger, M; Bragard, C; Caffier, D; Candresse, T; Chatzivassiliou, E; Dehnen-Schmutz, K; Gilioli, G; Gregoire, JC; Miret, JAJ; MacLeod, A; Navarro, MN; Niere, B; Parnell, S; Potting, R; Rafoss, T; Rossi, V; Urek, G; Van Bruggen, A; Van der Werf, W; | 2017 | Pest categorisation of Gremmeniella abietina | EFSA JOURNAL |

| | | | |
|---|---|---|---|
| West, J; Winter, S; Boberg, J; Gonthier, P; Pautasso, M | | | |
| Jeger, M; Bragard, C; Caffier, D; Candresse, T; Chatzivassiliou, E; Dehnen-Schmutz, K; Gilioli, G; Gregoire, JC; Miret, JAJ; Navarro, MN; Niere, B; Parnell, S; Potting, R; Rafoss, T; Rossi, V; Urek, G; Van Bruggen, A; Van der Werf, W; West, J; Winter, S; Kertesz, V; MacLeod, A | 2018 | Pest categorisation of Lopholeucaspis japonica | EFSA JOURNAL |
| Jeger, M; Bragard, C; Chatzivassiliou, E; Dehnen-Schmutz, K; Gilioli, G; Miret, JAJ; MacLeod, A; Navarro, MN; Niere, B; Parnell, S; Potting, R; Rafoss, T; Urek, G; Van Bruggen, A; Van der Werf, W; West, J; Winter, S; Santini, A; Tsopelas, P; Vloutoglou, I; Pautasso, M; Rossi, V | 2016 | Risk assessment and reduction options for Ceratocystis platani in the EU | EFSA JOURNAL |
| Jeger, M; Caffier, D; Candresse, T; Chatzivassiliou, E; Dehnen-Schmutz, K; Gilioli, G; Gregoire, JC; Miret, JAJ; MacLeod, A; Navarro, MN; Niere, B; Parnell, S; Potting, R; Rafoss, T; Rossi, V; Urek, G; Van Bruggen, A; Van der Werf, W; West, J; Winter, S; Almeida, R; Bosco, D; Jacques, MA; Landa, B; Purcell, A; Saponari, M; Czwienczek, E; Delbianco, A; Stancanelli, G; Bragard, C | 2018 | Updated pest categorisation of Xylella fastidiosa | EFSA JOURNAL |
| Jepsen, JU; Biuw, M; Ims, RA; Kapari, L; Schott, T; Vindstad, OPL; Hagen, SB | 2013 | Ecosystem Impacts of a Range Expanding Forest Defoliator at the Forest-Tundra Ecotone | ECOSYSTEMS |
| Jiang, HZ; Cao, CX; Chen, W; Fang, Z; Liu, C | 2016 | SIMULATION AND PREDICTION OF THE SPATIOTEMPORAL TRANSMISSION OF SUDDEN OAK DEATH (SOD) BASED ON SPATIAL INFORMATION TECHNOLOGY | 2016 IEEE INTERNATIONAL GEOSCIENCE AND REMOTE SENSING SYMPOSIUM (IGARSS) |
| Jnov, E | 2019 | Emerging and threatening vector-borne zoonoses in the world and in Europe: a brief update | PATHOGENS AND GLOBAL HEALTH |
| Johansson, T; Gibb, H; Hilszczanski, J; Pettersson, RB; Hjalten, J; Atlegrim, O; Ball, JP; Danell, K | 2006 | Conservation-oriented manipulations of coarse woody debris affect its value as habitat for spruce-infesting bark and | CANADIAN JOURNAL OF FOREST RESEARCH |

| | | | |
|---|---|---|---|
| | | ambrosia beetles (Coleoptera : Scolytinae) in northern Sweden | |
| Johnson, DM; Liebhold, AM; Bjornstad, ON; McManus, ML | 2005 | Circumpolar variation in periodicity and synchrony among gypsy moth populations | JOURNAL OF ANIMAL ECOLOGY |
| Jonsson, AM; Harding, S; Krokene, P; Lange, H; Lindelow, A; Okland, B; Ravn, HP; Schroeder, LM | 2011 | Modelling the potential impact of global warming on Ips typographus voltinism and reproductive diapause | CLIMATIC CHANGE |
| Joutsensaari, J; Yli-Pirila, P; Korhonen, H; Arola, A; Blande, JD; Heijari, J; Kivimaenpaa, M; Mikkonen, S; Hao, L; Miettinen, P; Lyytikainen-Saarenmaa, P; Faiola, CL; Laaksonen, A; Holopainen, JK | 2015 | Biotic stress accelerates formation of climate-relevant aerosols in boreal forests | ATMOSPHERIC CHEMISTRY AND PHYSICS |
| Junttila, S; Holopainen, M; Vastaranta, M; Lyytikainen-Saarenmaa, P; Kaartinen, H; Hyyppa, J; Hyyppa, H | 2019 | The potential of dual-wavelength terrestrial lidar in early detection of Ips typographus (L.) infestation - Leaf water content as a proxy | REMOTE SENSING OF ENVIRONMENT |
| Jurc, M; Cerny, M; Jurc, D | 2012 | FIRST RECORD OF ALIEN PEST Ophiomyia kwansonis (DIPTERA: AGROMYZIDAE) IN EUROPE AND ITS PHYTOSANITARY SIGNIFICANCE | SUMARSKI LIST |
| Kacprzyk, M | 2014 | Effect of microsite conditions on colonization of cambio-xylophagous insects on Norway spruce branches left after the silvicultural treatments | SYLWAN |
| Kaitera, J; Hantula, J; Nevalainen, S | 2011 | Distribution and frequency of Cronartium flaccidum on Melampyrum spp. in permanent sample plots in Finland | SCANDINAVIAN JOURNAL OF FOREST RESEARCH |
| Kantola, T; Vastaranta, M; Lyytikainen-Saarenmaa, P; Holopainen, M; Kankare, V; Talvitie, M; Hyyppa, J | 2013 | Classification of Needle Loss of Individual Scots Pine Trees by Means of Airborne Laser Scanning | FORESTS |
| Karlsson, PS; Tenow, O; Bylund, H; Hoogesteger, J; Weih, M | 2004 | Determinants of mountain birch growth in situ: effects of | ECOGRAPHY |

| | | | |
|---|---|---|---|
| | | temperature and herbivory | |
| Karvemo, S; Bjorkman, C; Johansson, T; Weslien, J; Hjalten, J | 2017 | Forest restoration as a double-edged sword: the conflict between biodiversity conservation and pest control | JOURNAL OF APPLIED ECOLOGY |
| Karvemo, S; Johansson, V; Schroeder, M; Ranius, T | 2016 | Local colonization-extinction dynamics of a tree-killing bark beetle during a large-scale outbreak | ECOSPHERE |
| Karvemo, S; Meurling, S; Berger, D; Hoglund, J; Laurila, A | 2018 | Effects of host species and environmental factors on the prevalence of Batrachochytrium dendrobatidis in northern Europe | PLOS ONE |
| Karvemo, S; Rogell, B; Schroeder, M | 2014 | Dynamics of spruce bark beetle infestation spots: Importance of local population size and landscape characteristics after a storm disturbance | FOREST ECOLOGY AND MANAGEMENT |
| Karvemo, S; Van Boeckel, TP; Gilbert, M; Gregoire, JC; Schroeder, M | 2014 | Large-scale risk mapping of an eruptive bark beetle - Importance of forest susceptibility and beetle pressure | FOREST ECOLOGY AND MANAGEMENT |
| Kausrud, K; Okland, B; Skarpaas, O; Gregoire, JC; Erbilgin, N; Stenseth, NC | 2012 | Population dynamics in changing environments: the case of an eruptive forest pest species | BIOLOGICAL REVIEWS |
| Kautz, M; Dworschak, K; Gruppe, A; Schopf, R | 2011 | Quantifying spatio-temporal dispersion of bark beetle infestations in epidemic and non-epidemic conditions | FOREST ECOLOGY AND MANAGEMENT |
| Kaynas, BY; Gurkan, B | 2005 | Changes in Buprestidae (Coleoptera) community with successional age after fire in a Pinus brutia forest | JOURNAL OF PEST SCIENCE |
| Kazenas, VL; Temreshev, II; Esenbekova, PA | 2016 | REVIEW OF THE SANITARY CONDITION OF CONIFEROUS FORESTS IN WINDFALL PLACES IN THE ILE-ALATAU NATIONAL PARK (KAZAKHSTAN) IN 2011-2015 | NATURE CONSERVATION RESEARCH |

| | | | |
|---|---|---|---|
| Keiner, R; Gruselle, MC; Michalzik, B; Popp, J; Frosch, T | 2015 | Raman spectroscopic investigation of (CO2)-C-13 labeling and leaf dark respiration of Fagus sylvatica L. (European beech) | ANALYTICAL AND BIOANALYTICAL CHEMISTRY |
| Keles, S; Sivrikaya, F; Cakir, G | 2007 | Temporal changes in forest landscape patterns in Artvin Forest Planning Unit, Turkey | ENVIRONMENTAL MONITORING AND ASSESSMENT |
| Keles, S; Sivrikaya, F; Cakir, G; Baskent, EZ; Kose, S | 2008 | Spatial and temporal changes in forest cover in Turkey's Artvin Forest, 1972-2002 | POLISH JOURNAL OF ENVIRONMENTAL STUDIES |
| Kesik-Maliszewska, J; Krzysiak, MK; Grochowska, M; Lechowski, L; Chase, C; Larska, M | 2018 | EPIDEMIOLOGY OF SCHMALLENBERG VIRUS IN EUROPEAN BISON (BISON BONASUS) IN POLAND | JOURNAL OF WILDLIFE DISEASES |
| Keskitalo, ECH; Klenk, N; Bullock, R; Smith, AL; Bazely, DR | 2011 | Preparing for and Responding to Disturbance: Examples from the Forest Sector in Sweden and Canada | FORESTS |
| Keskitalo, ECH; Pettersson, M; Ambjornsson, EL; Davis, EJ | 2016 | Agenda-setting and framing of policy solutions for forest pests in Canada and Sweden: Avoiding beetle outbreaks? | FOREST POLICY AND ECONOMICS |
| Keszthelyi, S; Feher, B; Somfalvi-Toth, K | 2019 | Worldwide distribution and theoretical spreading of Trichoferus campestris (Coleoptera: Cerambycidae) depending on the main climatic elements | ENTOMOLOGICAL SCIENCE |
| Khakimulina, T; Fraver, S; Drobyshev, I | 2016 | Mixed-severity natural disturbance regime dominates in an old-growth Norway spruce forest of northwest Russia | JOURNAL OF VEGETATION SCIENCE |
| Khasanov, BF; Sandlersky, RB | 2018 | Does insect induced defoliation affect anatomical structure of oak wood? | DENDROCHRONOLOGIA |
| Kirichenko, N; Flament, J; Baranchikov, Y; Gregoire, JC | 2011 | Larval performances and life cycle completion of the Siberian moth, Dendrolimus sibiricus | EUROPEAN JOURNAL OF FOREST RESEARCH |

| | | | |
|---|---|---|---|
| | | (Lepidoptera: Lasiocampidae), on potential host plants in Europe: a laboratory study on potted trees | |
| Kirkendall, LR; Faccoli, M; Ye, H | 2008 | Description of the Yunnan shoot borer, Tomicus yunnanensis Kirkendall & Faccoli sp n. (Curculionidae, Scolytinae), an unusually aggressive pine shoot beetle from southern China, with a key to the species of Tomicus | ZOOTAXA |
| Kivimaenpaa, M; Ghimire, RP; Sutinen, S; Haikio, E; Kasurinen, A; Holopainen, T; Holopainen, JK | 2016 | Increases in volatile organic compound emissions of Scots pine in response to elevated ozone and warming are modified by herbivory and soil nitrogen availability | EUROPEAN JOURNAL OF FOREST RESEARCH |
| Klapwijk, MJ; Csoka, G; Hirka, A; Bjorkman, C | 2013 | Forest insects and climate change: long-term trends in herbivore damage | ECOLOGY AND EVOLUTION |
| Klemola, T; Andersson, T; Ruohomaki, K | 2014 | Delayed density-dependent parasitism of eggs and pupae as a contributor to the cyclic population dynamics of the autumnal moth | OECOLOGIA |
| Klemola, T; Andersson, T; Ruohomaki, K | 2008 | Fecundity of the autumnal moth depends on pooled geometrid abundance without a time lag: implications for cyclic population dynamics | JOURNAL OF ANIMAL ECOLOGY |
| Klemola, T; Klemola, N; Andersson, T; Ruohomaki, K | 2007 | Does immune function influence population fluctuations and level of parasitism in the cyclic geometrid moth? | POPULATION ECOLOGY |
| Klemola, T; Ruohomaki, K; Andersson, T; Neuvonen, S | 2004 | Reduction in size and fecundity of the autumnal moth, Epirrita autumnata, in the increase phase of a population cycle | OECOLOGIA |

| | | | |
|---|---|---|---|
| Klimetzek, D; Yue, CF | 1997 | Climate and forest insect outbreaks | BIOLOGIA |
| Klitgaard, K; Hojgaard, J; Isbrand, A; Madsen, JJ; Thorup, K; Bodker, R | 2019 | Screening for multiple tick-borne pathogens in Ixodes ricinus ticks from birds in Denmark during spring and autumn migration seasons | TICKS AND TICK-BORNE DISEASES |
| Kocacevik, S; Sevim, A; Eroglu, M; Demirbag, Z; Demir, I | 2016 | Virulence and horizontal transmission of Beauveria pseudobassiana SA Rehner & Humber in Ips sexdentatus and Ips typographus (Coleoptera: Curculionidae) | TURKISH JOURNAL OF AGRICULTURE AND FORESTRY |
| Kohl, PL; Rutschmann, B | 2018 | The neglected bee trees: European beech forests as a home for feral honey bee colonies | PEERJ |
| Kohler, M; Kunz, J; Herrmann, J; Hartmann, P; Jansone, L; Puhlmann, H; von Wilpert, K; Bauhus, J | 2019 | The Potential of Liming to Improve Drought Tolerance of Norway Spruce [Picea abies (L.) Karst.] | FRONTIERS IN PLANT SCIENCE |
| Kolodziej-Sobocinska, M; Zalewski, A; Kowalczyk, R | 2014 | Sarcoptic mange vulnerability in carnivores of the BiaowieA1/4a Primeval Forest, Poland: underlying determinant factors | ECOLOGICAL RESEARCH |
| Koprowski, M; Duncker, P | 2012 | Tree ring width and wood density as the indicators of climatic factors and insect outbreaks affecting spruce growth | ECOLOGICAL INDICATORS |
| Kosik-Bogacka, D; Kuzna-Grygiel, W; Bukowska, K | 2004 | The prevalence of spirochete Borrelia burgdorferi sensu lato in ticks Ixodes ricinus and mosquitoes Aedes spp. within a selected recreational area in the city of Szczecin | ANNALS OF AGRICULTURAL AND ENVIRONMENTAL MEDICINE |
| Kosunen, M; Kantola, T; Starr, M; Blomqvist, M; Talvitie, M; Lyytikainen-Saarenmaa, P | 2017 | Influence of soil and topography on defoliation intensity during an extended outbreak of the common | IFOREST-BIOGEOSCIENCES AND FORESTRY |

| | | | |
|---|---|---|---|
| | | pine sawfly (Diprion pini L.) | |
| Kosunen, M; Lyytikainen-Saarenmaa, P; Ojanen, P; Blomqvist, M; Starr, M | 2019 | Response of Soil Surface Respiration to Storm and Ips typographus (L.) Disturbance in Boreal Norway Spruce Stands | FORESTS |
| Kotilinek, M; Tesitelova, T; Jersakova, J | 2015 | Biological Flora of the British Isles: Neottia ovata | JOURNAL OF ECOLOGY |
| Kouki, J; Lyytikainen-Saarenmaa, P; Henttonen, H; Niemela, P | 1998 | Cocoon predation an diprionid sawflies: the effect of forest fertility | OECOLOGIA |
| Kowalec, M; Szewczyk, T; Welc-Faleciak, R; Sinski, E; Karbowiak, G; Bajer, A | 2019 | Rickettsiales Occurrence and Co-occurrence in Ixodes ricinus Ticks in Natural and Urban Areas | MICROBIAL ECOLOGY |
| Kowalski, T; Sowa, J; Lakomy, P | 2019 | Mycobiota in trunks of dying spruce trees in the 'Puszcza Bialowieska' Promotional Forest Complex and its ecological function | SYLWAN |
| Kozlov, MV; Filippov, BY; Zubrij, NA; Zverev, V | 2015 | Abrupt changes in invertebrate herbivory on woody plants at the forest-tundra ecotone | POLAR BIOLOGY |
| Krascsenitsova, E; Kozanek, M; Ferencik, J; Roller, L; Stauffer, C; Bertheau, C | 2013 | Impact of the Carpathians on the genetic structure of the spruce bark beetle Ips typographus | JOURNAL OF PEST SCIENCE |
| Kruger, DH; Ulrich, RG; Hofmann, J | 2013 | Hantaviruses as Zoonotic Pathogens in Germany | DEUTSCHES ARZTEBLATT INTERNATIONAL |
| Krupa, SV; Moncrief, JF | 2002 | An integrative analysis of the roles of atmospheric deposition and land management practices on nitrogen in the US agricultural sector | ENVIRONMENTAL POLLUTION |
| Krzysiak, MK; Iwaniak, W; Kesik-Maliszewska, J; Olech, W; Larska, M | 2017 | Serological Study of Exposure to Selected Arthropod-Borne Pathogens in European Bison (Bison bonasus) in Poland | TRANSBOUNDARY AND EMERGING DISEASES |
| Kubiak, K; Damszel, M; Sikora, K; Przemieniecki, S; Malecka, M; Sierota, Z | 2017 | Colonization of Fungi and Bacteria in Stumps and Roots of Scots Pine after Thinning and Treatment with Rotstop | JOURNAL OF PHYTOPATHOLOGY |

| | | | |
|---|---|---|---|
| Kulakowski, D; Seidl, R; Holeksa, J; Kuuluvainen, T; Nagel, TA; Panayotov, M; Svoboda, M; Thorn, S; Vacchiano, G; Whitlock, C; Wohlgemuth, T; Bebi, P | 2017 | A walk on the wild side: Disturbance dynamics and the conservation and management of European mountain forest ecosystems | FOREST ECOLOGY AND MANAGEMENT |
| Kupkova, L; Potuckova, M; Lhotakova, Z; Albrechtova, J | 2018 | Forest cover and disturbance changes, and their driving forces: A case study in the Ore Mountains, Czechia, heavily affected by anthropogenic acidic pollution in the second half of the 20th century | ENVIRONMENTAL RESEARCH LETTERS |
| Labbe, F; Fontaine, MC; Robin, C; Dutech, C | 2017 | Genetic signatures of variation in population size in a native fungal pathogen after the recent massive plantation of its host tree | HEREDITY |
| Lackovic, N; Pernek, M; Bertheau, C; Franjevic, D; Stauffer, C; Avtzis, DN | 2018 | Limited Genetic Structure of Gypsy Moth Populations Reflecting a Recent History in Europe | INSECTS |
| Lamentowicz, M; Mueller, M; Galka, M; Barabach, J; Milecka, K; Goslar, T; Binkowski, M | 2015 | Reconstructing human impact on peatland development during the past 200 years in CE Europe through biotic proxies and X-ray tomography | QUATERNARY INTERNATIONAL |
| Langstrom, B; Annila, E; Hellqvist, C; Varama, M; Niemela, P | 2001 | Tree mortality, needle biomass recovery and growth losses in Scots pine following defoliation by Diprion pini (L.) and subsequent attack by Tomicus piniperda (L.) | SCANDINAVIAN JOURNAL OF FOREST RESEARCH |
| Latalowa, M; Pedziszewska, A; Maciejewska, E; Swieta-Musznicka, J | 2013 | Tilia forest dynamics, Kretzschmaria deusta attack, and mire hydrology as palaeoecological proxies for mid-Holocene climate reconstruction in the Kashubian Lake District (N Poland) | HOLOCENE |
| Latalowa, M; Swieta-Musznicka, J; Slowinski, M; Pedziszewska, A; Noryskiewicz, AM; Zimny, M; | 2019 | Abrupt Alnus population decline at the end of the first millennium CE in | HOLOCENE |

| | | | |
|---|---|---|---|
| Obremska, M; Ott, F; Stivrins, N; Pasanen, L; Ilvonen, L; Holmstrom, L; Seppa, H | | Europe - The event ecology, possible causes and implications | |
| Latifi, H; Schumann, B; Kautz, M; Dech, S | 2014 | Spatial characterization of bark beetle infestations by a multidate synergy of SPOT and Landsat imagery | ENVIRONMENTAL MONITORING AND ASSESSMENT |
| Lazdins, A; Miezite, O; Bardule, A | 2011 | CHARACTERIZATION OF SEVERE DAMAGES OF SPRUCE (PICEA ABIES (L.) H.KARST.) STANDS IN RELATION TO SOIL PROPERTIES | RESEARCH FOR RURAL DEVELOPMENT 2011, VOL 2 |
| le Mellec, A; Gerold, G; Michalzik, B | 2011 | Insect herbivory, organic matter deposition and effects on belowground organic matter fluxes in a central European oak forest | PLANT AND SOIL |
| le Mellec, A; Michalzik, B | 2008 | Impact of a pine lappet (Dendrolimus pini) mass outbreak on C and N fluxes to the forest floor and soil microbial properties in a Scots pine forest in Germany | CANADIAN JOURNAL OF FOREST RESEARCH |
| Leal, I; Allen, E; Foord, B; Anema, J; Reisle, C; Uzunovic, A; Varga, A; James, D | 2015 | Detection of living Bursaphelenchus xylophilus in wood, using reverse transcriptase loop-mediated isothermal amplification ( RT-LAMP) | FOREST PATHOLOGY |
| Leigh, EG; Davidar, P; Dick, CW; Puyravaud, JP; Terborgh, J; ter Steege, H; Wright, SJ | 2004 | Why do some tropical forests have so many species of trees? | BIOTROPICA |
| Leverkus, AB; Benayas, JMR; Castro, J; Boucher, D; Brewer, S; Collins, BM; Donato, D; Fraver, S; Kishchuk, BE; Lee, EJ; Lindenmayer, DB; Lingua, E; Macdonald, E; Marzano, R; Rhoades, CC; Royo, A; Thorn, S; Wagenbrenner, JW; Waldron, K; Wohlgemuth, T; Gustafsson, L | 2018 | Salvage logging effects on regulating and supporting ecosystem services - a systematic map | CANADIAN JOURNAL OF FOREST RESEARCH |
| Li, GS; Osborne, J; Asiegbu, FO | 2006 | A macroarray expression analysis of novel cDNAs vital for growth initiation and primary metabolism during development of | ENVIRONMENTAL MICROBIOLOGY |

| | | | |
|---|---|---|---|
| | | Heterobasidion parviporum conidiospores | |
| Li, J; Shi, J; Luo, YQ; Heliovaara, K | 2013 | RESPONSES OF MONOPHAGOUS IPS SUBELONGATUS MOTSCHULSKY (COLEOPTERA: CURCULIONIDAE) AND POLYPHAGOUS LYMANTRIA DISPAR L. (LEPIDOPTERA: LYMANTRIIDAE) TO TREE SPECIES MIXTURE | ENTOMOLOGICAL NEWS |
| Li, S; Daudin, JJ; Piou, D; Robinet, C; Jactel, H | 2015 | Periodicity and synchrony of pine processionary moth outbreaks in France | FOREST ECOLOGY AND MANAGEMENT |
| Lilja, A; Rytkonen, A; Hantula, J; Muller, M; Parikka, P; Kurkela, T | 2011 | Introduced pathogens found on ornamentals, strawberry and trees in Finland over the past 20 years | AGRICULTURAL AND FOOD SCIENCE |
| Linares, JC; Senhadji, K; Herrero, A; Hodar, JA | 2014 | Growth patterns at the southern range edge of Scots pine: Disentangling the effects of drought and defoliation by the pine processionary caterpillar | FOREST ECOLOGY AND MANAGEMENT |
| Lindbladh, M; Fraver, S; Edvardsson, J; Felton, A | 2013 | Past forest composition, structures and processes - How paleoecology can contribute to forest conservation | BIOLOGICAL CONSERVATION |
| Lindroth, A; Lagergren, F; Grelle, A; Klemedtsson, L; Langvall, O; Weslien, P; Tuulik, J | 2009 | Storms can cause Europe-wide reduction in forest carbon sink | GLOBAL CHANGE BIOLOGY |
| Linnakoski, R; Mahilainen, S; Harrington, A; Vanhanen, H; Eriksson, M; Mehtatalo, L; Pappinen, A; Wingfield, MJ | 2016 | Seasonal Succession of Fungi Associated with Ips typographus Beetles and Their Phoretic Mites in an Outbreak Region of Finland | PLOS ONE |
| Lukasova, K; Holusa, J; Knizek, M | 2014 | Dendroctonus micans populations on Picea pungens in the center of a non-outbreak region contain few pathogens, parasites or predators: A new threat for urban forests? | URBAN FORESTRY & URBAN GREENING |

| | | | |
|---|---|---|---|
| Lundstrom, JO | 1999 | Mosquito-borne viruses in western Europe: A review | JOURNAL OF VECTOR ECOLOGY |
| Lurz, PWW; Rushton, SP; Wauters, LA; Bertolino, S; Currado, I; Mazzoglio, P; Shirley, MDF | 2001 | Predicting grey squirrel expansion in North Italy: a spatially explicit modelling approach | LANDSCAPE ECOLOGY |
| MacDonald, WL | 2003 | Dominating North American forest pathology issues of the 20th century | PHYTOPATHOLOGY |
| MacLeod, A; Evans, HF; Baker, RHA | 2002 | An analysis of pest risk from an Asian longhorn beetle (Anoplophora glabripennis) to hardwood trees in the European community | CROP PROTECTION |
| Maestre, FT; Cortina, J | 2004 | Are Pinus hatepensis plantations useful as a restoration tool in semiarid Mediterranean areas? | FOREST ECOLOGY AND MANAGEMENT |
| Mageroy, MH; Parent, G; Germanos, G; Giguere, I; Delvas, N; Maaroufi, H; Bauce, E; Bohlmann, J; Mackay, JJ | 2015 | Expression of the beta-glucosidase gene Pg beta glu-1 underpins natural resistance of white spruce against spruce budworm | PLANT JOURNAL |
| Majunke, C | 1995 | The important of needle-eating and wood-boring forest insects in the lowlands of northeastern Germany | MITTEILUNGEN DER DEUTSCHEN GESELLSCHAFT FUR ALLGEMEINE UND ANGEWANDTE ENTOMOLOGIE, BAND 10, HEFT 1-6, DEZEMBER 1995: VORTRAGE DER ENTOMOLOGENTAGUN IN GOTTINGEN, VOM 27. MARZ - 1. APRIL 1995 |
| Mannu, R; Gilioli, G; Luciano, P | 2017 | OCCUPANCY OF THE TERRITORY BY LYMANTRIA DISPAR (L.) (LEPIDOPTERA EREBIDAE) EGG MASSES AS A PREDICTIVE INDEX OF DAMAGE | REDIA-GIORNALE DI ZOOLOGIA |
| Marciulynas, A; Sirgedaite-Seziene, V; Zemaitis, P; Baliuckas, V | 2019 | The Resistance of Scots Pine (Pinus sylvestris L.) Half-sib Families to Heterobasidion annosum | FORESTS |
| Marcone, C | 2017 | Elm yellows: A phytoplasma disease of | FOREST PATHOLOGY |

| | | | |
|---|---|---|---|
| | | concern in forest and landscape ecosystems | |
| Marini, L; Lindelow, A; Jonsson, AM; Wulff, S; Schroeder, LM | 2013 | Population dynamics of the spruce bark beetle: a long-term study | OIKOS |
| Marini, L; Okland, B; Jonsson, AM; Bentz, B; Carroll, A; Forster, B; Gregoire, JC; Hurling, R; Nageleisen, LM; Netherer, S; Ravn, HP; Weed, A; Schroeder, M | 2017 | Climate drivers of bark beetle outbreak dynamics in Norway spruce forests | ECOGRAPHY |
| MARKALAS, S | 1992 | SITE AND STAND FACTORS RELATED TO MORTALITY-RATE IN A FIR FOREST AFTER A COMBINED INCIDENCE OF DROUGHT AND INSECT ATTACK | FOREST ECOLOGY AND MANAGEMENT |
| Mason, RR; Jennings, DT; Paul, HG; Wickman, BE | 1997 | Patterns of spider (Araneae) abundance during an outbreak of western spruce budworm (Lepidoptera: Tortricidae) | ENVIRONMENTAL ENTOMOLOGY |
| Mason, RR; Wickman, BE; Paul, HG; Torgersen, TR | 1998 | A pilot experiment of forest fertilization during an outbreak of the western spruce budworm in northeastern Oregon. | USDA FOREST SERVICE PACIFIC NORTHWEST RESEARCH STATION RESEARCH PAPER |
| Matek, M; Pernek, M | 2018 | First Record of Dendrolimus pini Outbreak on Aleppo Pine in Croatia and Severe Case of Population Collapse Caused by Entomopathogen Beauveria bassiana | SEEFOR-SOUTH-EAST EUROPEAN FORESTRY |
| Matisone, I; Matisons, R; Jansons, A | 2019 | Health Condition of European Ash in Young Stands of Diverse Composition | BALTIC FORESTRY |
| Matiu, M; Bothmann, L; Steinbrecher, R; Menzel, A | 2017 | Monitoring succession after a non-cleared windthrow in a Norway spruce mountain forest using webcam, satellite vegetation indices and turbulent CO2 exchange | AGRICULTURAL AND FOREST METEOROLOGY |
| Matua, GA; Van der Wal, DM; Locsin, RC | 2015 | Ebola hemorrhagic fever outbreaks: strategies for effective epidemic management, containment and control | BRAZILIAN JOURNAL OF INFECTIOUS DISEASES |

| | | | |
|---|---|---|---|
| Matysiak, A; Kapuscinski, R | 2007 | The contemporary issues related to forest management in the State Forests National Forest Holding | Quo Vadis, Forestry?, Proceedings |
| Mendel, Z; Assael, F; Zeidan, S; Zehavi, A | 1998 | Classical biological control of Palaeococcus fuscipennis (Burmeister) (Homoptera : Margarodidae) in Israel | BIOLOGICAL CONTROL |
| Menichetti, L; Leifeld, J; Kirova, L; Szidat, S; Zhiyanski, M | 2017 | Consequences of planned afforestation versus natural forest regrowth after disturbance for soil C stocks in Eastern European mountains | GEODERMA |
| Metcalfe, DB; Cherif, M; Jepsen, JU; Vindstad, OPL; Kristensen, JA; Belsing, U | 2019 | Ecological stoichiometry and nutrient partitioning in two insect herbivores responsible for large-scale forest disturbance in the Fennoscandian subarctic | ECOLOGICAL ENTOMOLOGY |
| Mezei, P; Blazenec, M; Grodzki, W; Skvarenina, J; Jakus, R | 2017 | Influence of different forest protection strategies on spruce tree mortality during a bark beetle outbreak | ANNALS OF FOREST SCIENCE |
| Mezei, P; Potterf, M; Skvarenina, J; Rasmussen, JG; Jakus, R | 2019 | Potential Solar Radiation as a Driver for Bark Beetle Infestation on a Landscape Scale | FORESTS |
| Mierzejewska, EJ; Estrada-Pena, A; Bajer, A | 2017 | Spread of Dermacentor reticulatus is associated with the loss of forest area | EXPERIMENTAL AND APPLIED ACAROLOGY |
| Milanovic, S; Lazarevic, J; Popovic, Z; Miletic, Z; Kostic, M; Radulovic, Z; Karadzic, D; Vuleta, A | 2014 | Preference and performance of the larvae of Lymantria dispar (Lepidoptera: Lymantriidae) on three species of European oaks | EUROPEAN JOURNAL OF ENTOMOLOGY |
| Mjaaseth, RR; Hagen, SB; Yoccoz, NG; Ims, RA | 2005 | Phenology and abundance in relation to climatic variation in a sub-arctic insect herbivore-mountain birch system | OECOLOGIA |
| Modlinger, R; Liska, J | 2016 | Review of Lepidoptera with trophic relationships | CENTRAL EUROPEAN FORESTRY JOURNAL |

| | | | |
|---|---|---|---|
| | | to Picea abies (L.) in the conditions of Czechia | |
| Moller, K; Hentschel, R; Wenning, A; Schroder, J | 2017 | Improved Outbreak Prediction for Common Pine Sawfly (Diprion pini L.) by Analyzing Floating 'ClimaticWindows' as Keys for Changes in Voltinism | FORESTS |
| Moraal, LG | 1996 | Bionomics of Haematoloma dorsatum (Hem, Cercopidae) in relation to needle damage in pine forests | ANZEIGER FUR SCHADLINGSKUNDE PFLANZENSCHUTZ UMWELTSCHUTZ |
| Muller, M; Job, H | 2009 | Managing natural disturbance in protected areas: Tourists' attitude towards the bark beetle in a German national park | BIOLOGICAL CONSERVATION |
| Muller, MM; Henttonen, HM; Penttila, R; Kulju, M; Helo, T; Kaitera, J | 2018 | Distribution of Heterobasidion butt rot in northern Finland | FOREST ECOLOGY AND MANAGEMENT |
| Mullett, MS; Brown, AV; Fraser, S; Baden, R; Tubby, KV | 2017 | Insights into the pathways of spread and potential origins of Dothistroma septosporum in Britain | FUNGAL ECOLOGY |
| Munzbergova, Z; Herben, T | 2005 | Seed, dispersal, microsite, habitat and recruitment limitation: identification of terms and concepts in studies of limitations | OECOLOGIA |
| Musolin, DL; Selikhovkin, AV; Shabunin, DA; Zviagintsev, VB; Baranchikov, YN | 2017 | Between Ash Dieback and Emerald Ash Borer: Two Asian Invaders in Russia and the Future of Ash in Europe | BALTIC FORESTRY |
| Nagy, NE; Ballance, S; Kvaalen, H; Fossdal, CG; Solheim, H; Hietala, AM | 2012 | Xylem defense wood of Norway spruce compromised by the pathogenic white-rot fungus Heterobasidion parviporum shows a prolonged period of selective decay | PLANTA |
| Nasi, R; Honkavaara, E; Blomqvist, M; Lyytikainen-Saarenmaa, P; Hakala, T; Viljanen, N; Kantola, T; Holopainen, M | 2018 | Remote sensing of bark beetle damage in urban forests at individual tree level using a novel | URBAN FORESTRY & URBAN GREENING |

| | | | |
|---|---|---|---|
| | | hyperspectral camera from UAV and aircraft | |
| Nasi, R; Honkavaara, E; Lyytikainen-Saarenmaa, P; Blomqvist, M; Litkey, P; Hakala, T; Viljanen, N; Kantola, T; Tanhuanpaa, T; Holopainen, M | 2015 | Using UAV-Based Photogrammetry and Hyperspectral Imaging for Mapping Bark Beetle Damage at Tree-Level | REMOTE SENSING |
| Neary, DG | 2016 | Long-Term Forest Paired Catchment Studies: What Do They Tell Us That Landscape-Level Monitoring Does Not? | FORESTS |
| Negron, JF; Schaupp, WC; Johnson, E | 2000 | Development and validation of a fixed-precision sequential sampling plan for estimating brood adult density of Dendroctonus pseudotsugae (Coleoptera : Scolytidae) | CANADIAN ENTOMOLOGIST |
| Nemer, N; Bal, J; Bechara, E; Frerot, B | 2014 | Pheromone identification of the cedar shoot moth Dichelia cedricola Diakonoff (Lepidoptera: Tortricidae) | ANNALES DE LA SOCIETE ENTOMOLOGIQUE DE FRANCE |
| Netherer, S; Schopf, A | 2010 | Potential effects of climate change on insect herbivores in European forests-General aspects and the pine processionary moth as specific example | FOREST ECOLOGY AND MANAGEMENT |
| Neuvonen, S; Viiri, H | 2017 | Changing Climate and Outbreaks of Forest Pest Insects in a Cold Northern Country, Finland | INTERCONNECTED ARCTIC - UARCTIC CONGRESS 2016 |
| Nevalainen, S; Lindgren, M; Pouttu, A; Heinonen, J; Hongisto, M; Neuvonen, S | 2010 | Extensive tree health monitoring networks are useful in revealing the impacts of widespread biotic damage in boreal forests | ENVIRONMENTAL MONITORING AND ASSESSMENT |
| Neves, D; Caetano, P; Oliveira, J; Maia, C; Horta, M; Sousa, N; Salgado, M; Dionisio, L; Magan, N; Cravador, A | 2014 | Anti-Phytophthora cinnamomi activity of Phlomis purpurea plant and root extracts | EUROPEAN JOURNAL OF PLANT PATHOLOGY |
| Nielsen, MM; Heurich, M; Malmberg, B; Brun, A | 2014 | Automatic Mapping of Standing Dead Trees after an Insect Outbreak Using the Window | JOURNAL OF FORESTRY |

| | | | |
|---|---|---|---|
| | | Independent Context Segmentation Method | |
| Niemczyk, M; Karwanski, M; Grzybowska, U | 2017 | Effect of Environmental Factors on Occurrence of Cockchafers (Melolontha spp.) in Forest Stands | BALTIC FORESTRY |
| Niemczyk, M; Sierpinska, A; Tereba, A; Sokolowski, K; Przybylski, P | 2019 | Natural occurrence of Beauveria spp. in outbreak areas of cockchafers (Melolontha spp.) in forest soils from Poland | BIOCONTROL |
| Nikolov, C; Konopka, B; Kajba, M; Galko, J; Kunca, A; Jansky, L | 2014 | Post-disaster Forest Management and Bark Beetle Outbreak in Tatra National Park, Slovakia | MOUNTAIN RESEARCH AND DEVELOPMENT |
| Ocasio-Morales, RG; Tsopelas, P; Harrington, TC | 2007 | Origin of Ceratocystis platani on native Platanus orientalis in Greece and its impact on natural forests | PLANT DISEASE |
| O'Hanlon, R; Choiseul, J; Brennan, JM; Grogan, H | 2018 | Assessment of the eradication measures applied to Phytophthora ramorum in Irish Larix kaempferi forests | FOREST PATHOLOGY |
| Ohrn, P; Bjorklund, N; Langstrom, B | 2018 | Occurrence, performance and shoot damage of Tomicus piniperda in pine stands in southern Sweden after storm-felling | JOURNAL OF APPLIED ENTOMOLOGY |
| Okland, B; Liebhold, AM; Bjornstad, ON; Erbilgin, N; Krokene, P | 2005 | Are bark beetle outbreaks less synchronous than forest Lepidoptera outbreaks? | OECOLOGIA |
| Okland, B; Netherer, S; Marini, L | 2015 | The Eurasian Spruce Bark Beetle: The Role of Climate | CLIMATE CHANGE AND INSECT PESTS |
| Okland, B; Nikolov, C; Krokene, P; Vakula, J | 2016 | Transition from windfall- to patch-driven outbreak dynamics of the spruce bark beetle Ips typographus | FOREST ECOLOGY AND MANAGEMENT |
| Oliva, J; Bendz-Hellgren, M; Stenlid, J | 2011 | Spread of Heterobasidion annosum s.s. and Heterobasidion parviporum in Picea abies 15 years after stump inoculation | FEMS MICROBIOLOGY ECOLOGY |

| | | | |
|---|---|---|---|
| Oliva, J; Colinas, C | 2010 | Epidemiology of Heterobasidion abietinum and Viscum album on silver fir (Abies alba) stands of the Pyrenees | FOREST PATHOLOGY |
| Oliva, J; Rommel, S; Fossdal, CG; Hietala, AM; Nemesio-Gorriz, M; Solheim, H; Elfstrand, M | 2015 | Transcriptional responses of Norway spruce (Picea abies) inner sapwood against Heterobasidion parviporum | TREE PHYSIOLOGY |
| Olofsson, J; Ericson, L; Torp, M; Stark, S; Baxter, R | 2011 | Carbon balance of Arctic tundra under increased snow cover mediated by a plant pathogen | NATURE CLIMATE CHANGE |
| Olsson, PO; Kantola, T; Lyytikainen-Saarenmaa, P; Jonsson, AM; Eklundh, L | 2016 | Development of a method for monitoring of insect induced forest defoliation - limitation of MODIS data in Fennoscandian forest landscapes | SILVA FENNICA |
| Olsson, PO; Lindstrom, J; Eldundh, L | 2016 | Near real-time monitoring of insect induced defoliation in subalpine birch forests with MODIS derived NDVI | REMOTE SENSING OF ENVIRONMENT |
| Onet, A; Teusdea, A; Boja, N; Domuta, C; Onet, C | 2016 | Effects of common oak (Quercus robur L.) defolition on the soil properties of an oak forest in Western Plain of Romania | ANNALS OF FOREST RESEARCH |
| Orczewska, A; Czortek, P; Jaroszewicz, B | 2019 | The impact of salvage logging on herb layer species composition and plant community recovery in Bialowie(z) over dota Forest | BIODIVERSITY AND CONSERVATION |
| Orlova-Bienkowskaja, MJ | 2017 | Main Trends of Invasion Processes in Beetles (Coleoptera) of European Russia | RUSSIAN JOURNAL OF BIOLOGICAL INVASIONS |
| Orlova-Bienkowskaja, MJ; Bienkowski, AO | 2017 | ALIEN COCCINELLIDAE (LADYBIRDS) IN SOCHI NATIONAL PARK AND ITS VICINITY, RUSSIA | NATURE CONSERVATION RESEARCH |
| Otsu, K; Pla, M; Vayreda, J; Brotons, L | 2018 | Calibrating the Severity of Forest Defoliation by Pine Processionary Moth | SENSORS |

| | | | |
|---|---|---|---|
| | | with Landsat and UAV Imagery | |
| Overbeck, M; Schmidt, M | 2012 | Modelling infestation risk of Norway spruce by Ips typographus (L.) in the Lower Saxon Harz Mountains (Germany) | FOREST ECOLOGY AND MANAGEMENT |
| Ozcan, GE | 2017 | ASSESSMENT OF Ips sexdentatus POPULATION CONSIDERING THE CAPTURE IN PHEROMONE TRAPS AND THEIR DAMAGES UNDER NON-EPIDEMIC CONDITIONS | SUMARSKI LIST |
| Pamerleau-Couture, E; Rossi, S; Pothier, D; Krause, C | 2019 | Wood properties of black spruce (Picea mariana (Mill.) BSP) in relation to ring width and tree height in even- and uneven-aged boreal stands | ANNALS OF FOREST SCIENCE |
| Panzavolta, T; Panichi, A; Bracalini, M; Croci, F; Benigno, A; Ragazzi, A; Tiberi, R; Moricca, S | 2018 | Tree pathogens and their insect-mediated transport: Implications for oak tree die-off in a natural park area | GLOBAL ECOLOGY AND CONSERVATION |
| Parak, M; Kulfan, J; Zach, P | 2015 | Are the moth larvae able to withstand tree fall caused by wind storm? | ANNALS OF FOREST RESEARCH |
| Pennacchio, F; Danti, R; Benassai, D; Squarcini, M; Marziali, L; Di Lonardo, V; Roversi, PF | 2013 | A NEW ADDITIONAL RECORD OF PHLOEOSINUS ARMATUS REITTER FROM ITALY (COLEOPTERA CURCULIONIDAE SCOLYTINAE) | REDIA-GIORNALE DI ZOOLOGIA |
| Pennacchio, F; Santini, L; Francardi, V | 2012 | BIOECOLOGICAL NOTES ON XYLOSANDRUS COMPACTUS (EICHHOFF) (COLEOPTERA CURCULIONIDAE SCOLYTINAE), A SPECIES RECENTLY RECORDED INTO ITALY | REDIA-GIORNALE DI ZOOLOGIA |
| Peris-Felipo, FJ; Jimenez-Peydro, R | 2011 | Biodiversity within the subfamily Alyssinae (Hymenoptera, Braconidae) in the | REVISTA BRASILEIRA DE ENTOMOLOGIA |

| | | | |
|---|---|---|---|
| | | Natural Park Penas de Aya (Spain) | |
| Pernek, M | 2018 | NEW CALCULATION OF CRITICAL NUMBER OF GYPSY MOTH (LYMANTRIA DISPAR L.) EGG MASSES FOR BETTER POPULATION DENSITY PROGNOSIS | SUMARSKI LIST |
| Pernek, M; Matosevic, D; Hrasovec, B; Kucinic, M; Wegensteiner, R | 2009 | Occurrence of pathogens in outbreak populations of Pityokteines spp. (Coleoptera, Curculionidae, Scolytinae) in silver fir forests | JOURNAL OF PEST SCIENCE |
| Perot, T; Vallet, P; Archaux, F | 2013 | Growth compensation in an oak-pine mixed forest following an outbreak of pine sawfly (Diprion pini) | FOREST ECOLOGY AND MANAGEMENT |
| Persson, Y; Vasaitis, R; Langstrom, B; Ohrn, P; Ihrmark, K; Stenlid, J | 2009 | Fungi Vectored by the Bark Beetle Ips typographus Following Hibernation Under the Bark of Standing Trees and in the Forest Litter | MICROBIAL ECOLOGY |
| Pertot, I; Gobbin, D; De Luca, F; Prodorutti, D | 2008 | Methods of assessing the incidence of Armillaria root rot across viticultural areas and the pathogen's genetic diversity and spatial-temporal pattern in northern Italy | CROP PROTECTION |
| Peters, FS; Holweg, CL; Rigling, D; Metzler, B | 2012 | Chestnut blight in south-western Germany: multiple introductions of Cryphonectria parasitica and slow hypovirus spread | FOREST PATHOLOGY |
| Petucco, C; Andres-Domenech, P | 2018 | Land expectation value and optimal rotation age of maritime pine plantations under multiple risks | JOURNAL OF FOREST ECONOMICS |
| Peverieri, GS; Furlan, P; Simoni, S; Strong, WB; Roversi, PF | 2012 | Laboratory evaluation of Gryon pennsylvanicum (Ashmead) (Hymenoptera, Platygastridae) as a biological control agent | BIOLOGICAL CONTROL |

| | | | |
|---|---|---|---|
| | | of Leptoglossus occidentalis Heidemann (Heteroptera, Coreidae) | |
| PFEFFER, A; SKUHRAVY, V | 1995 | IPS-TYPOGRAPHUS L (COL, SCOLYTIDAE) IN THE CZECH-REPUBLIC | ANZEIGER FUR SCHADLINGSKUNDE PFLANZENSCHUTZ UMWELTSCHUTZ |
| Pimentel, C; Nilsson, JA | 2009 | RESPONSE OF PASSERINE BIRDS TO AN IRRUPTION OF A PINE PROCESSIONARY MOTH THAUMETOPOEA PITYOCAMPA POPULATION WITH A SHIFTED PHENOLOGY | ARDEOLA-INTERNATIONAL JOURNAL OF ORNITHOLOGY |
| Pimentel, C; Nilsson, JA | 2007 | Response of Great Tits Parus major to an irruption of a Pine Processionary Moth Thaumetopoea pityocampa population with a shifted phenology | ARDEA |
| Pineau, X; David, G; Peter, Z; Salle, A; Baude, M; Lieutier, F; Jactel, H | 2017 | Effect of temperature on the reproductive success, developmental rate and brood characteristics of Ips sexdentatus (Boern.) | AGRICULTURAL AND FOREST ENTOMOLOGY |
| Piri, T; Valkonen, S | 2013 | Incidence and spread of Heterobasidion root rot in uneven-aged Norway spruce stands | CANADIAN JOURNAL OF FOREST RESEARCH |
| Pisetta, M; Montecchio, L; Longa, CMO; Salvadori, C; Zottele, F; Maresi, G | 2012 | Green alder decline in the Italian Alps | FOREST ECOLOGY AND MANAGEMENT |
| Poher, Y; Ponel, P; Medail, F; Andrieu-Ponel, V; Guiter, F | 2017 | Holocene environmental history of a small Mediterranean island in response to sea-level changes, climate and human impact | PALAEOGEOGRAPHY PALAEOCLIMATOLOGY PALAEOECOLOGY |
| Potter, C; Kumar, V; Klooster, S; Nemani, R | 2007 | Recent history of trends in vegetation greenness and large-scale ecosystem disturbances in Eurasia | TELLUS SERIES B-CHEMICAL AND PHYSICAL METEOROLOGY |
| Poutsma, J; Loomans, AJM; Aukema, B; Heijerman, T | 2008 | Predicting the potential geographical distribution of the harlequin ladybird, Harmonia axyridis, using the CLIMEX model | BIOCONTROL |

| | | | |
|---|---|---|---|
| PRACH, K | 1994 | SUCCESSION OF WOODY SPECIES IN DERELICT SITES IN CENTRAL-EUROPE | ECOLOGICAL ENGINEERING |
| Probst, C; Gethmann, J; Amler, S; Globig, A; Knoll, B; Conraths, FJ | 2019 | The potential role of scavengers in spreading African swine fever among wild boar | SCIENTIFIC REPORTS |
| Prospero, S; Lung-Escarmant, B; Dutech, C | 2008 | Genetic structure of an expanding Armillaria root rot fungus (Armillaria ostoyae) population in a managed pine forest in southwestern France | MOLECULAR ECOLOGY |
| Przybyl, K; Karolewski, P; Oleksyn, J; Labedzki, A; Reich, PB | 2008 | Fungal diversity of Norway spruce litter: Effects of site conditions and premature leaf fall caused by bark beetle outbreak | MICROBIAL ECOLOGY |
| Pukkala, T | 2018 | Effect of species composition on ecosystem services in European boreal forest | JOURNAL OF FORESTRY RESEARCH |
| Punttila, P; Niemela, P; Karhu, K | 2004 | The impact of wood ants (Hymenoptera : Formicidae) on the structure of invertebrate community on mountain birch (Betula pubescens ssp czerepanovii) | ANNALES ZOOLOGICI FENNICI |
| Pureswaran, DS; De Grandpre, L; Pare, D; Taylor, A; Barrette, M; Morin, H; Regniere, J; Kneeshaw, DD | 2015 | Climate-induced changes in host tree-insect phenology may drive ecological state-shift in boreal forests | ECOLOGY |
| Quinn, L; O'Neill, PA; Harrison, J; Paskiewicz, KH; McCracken, AR; Cooke, LR; Grant, MR; Studholme, DJ | 2013 | Genome-wide sequencing of Phytophthora lateralis reveals genetic variation among isolates from Lawson cypress (Chamaecyparis lawsoniana) in Northern Ireland | FEMS MICROBIOLOGY LETTERS |
| Radocz, L; Tarcali, G | 2005 | Identification of natural infection of Quercus spp. by chestnut blight fungus (Cryphonectria parasistica) | PROCEEDINGS OF THE THIRD INTERNATIONAL CHESTNUT CONGRESS |

| | | | |
|---|---|---|---|
| Rahmani, R; Hedenstrom, E; Schroeder, M | 2015 | SPME collection and GC-MS analysis of volatiles emitted during the attack of male Polygraphus poligraphus (Coleoptera, Curcolionidae) on Norway spruce | ZEITSCHRIFT FUR NATURFORSCHUNG SECTION C-A JOURNAL OF BIOSCIENCES |
| Rassati, D; Toffolo, EP; Battisti, A; Faccoli, M | 2012 | Monitoring of the pine sawyer beetle Monochamus galloprovincialis by pheromone traps in Italy | PHYTOPARASITICA |
| Rizzi, A; Crotti, E; Borruso, L; Jucker, C; Lupi, D; Colombo, M; Daffonchio, D | 2013 | Characterization of the Bacterial Community Associated with Larvae and Adults of Anoplophora chinensis Collected in Italy by Culture and Culture-Independent Methods | BIOMED RESEARCH INTERNATIONAL |
| Roberge, JM; Bengtsson, SBK; Wulff, S; Snall, T | 2011 | Edge creation and tree dieback influence the patch-tracking metapopulation dynamics of a red-listed epiphytic bryophyte | JOURNAL OF APPLIED ECOLOGY |
| Robinet, C; Rousselet, J; Roques, A | 2014 | Potential spread of the pine processionary moth in France: preliminary results from a simulation model and future challenges | ANNALS OF FOREST SCIENCE |
| Rodriguez-Garcia, A; Martin, JA; Lopez, R; Sanz, A; Gil, L | 2016 | Effect of four tapping methods on anatomical traits and resin yield in Maritime pine (Pinus pinaster Ait.) | INDUSTRIAL CROPS AND PRODUCTS |
| Rolland, C; Baltensweiler, W; Petitcolas, V | 2001 | The potential for using Larix decidua ring widths in reconstructions of larch budmoth (Zeiraphera diniana) outbreak history: dendrochronological estimates compared with insect surveys | TREES-STRUCTURE AND FUNCTION |
| Rolland, C; Lemperiere, G | 2004 | Effects of climate on radial growth of Norway spruce and interactions with attacks by the bark beetle Dendroctonus | FOREST ECOLOGY AND MANAGEMENT |

| | | | |
|---|---|---|---|
| | | micans (Kug., Coleoptera : Scolytidae): a dendroecological study in the French Massif Central | |
| Romeralo, C; Santamaria, O; Pando, V; Diez, JJ | 2015 | Fungal endophytes reduce necrosis length produced by Gremmeniella abietina in Pinus halepensis seedlings | BIOLOGICAL CONTROL |
| Romon, P; Iturrondobeitia, JC; Gibson, K; Lindgren, BS; Goldarazena, A | 2007 | Quantitative association of bark beetles with pitch canker fungus and effects of verbenone on their semiochemical communication in monterey pine forests in Northern Spain | ENVIRONMENTAL ENTOMOLOGY |
| Root, C; Balbalian, C; Bierman, R; Geletka, LM; Anagnostakis, S; Double, M; MacDonald, W; Nuss, DL | 2005 | Multi-seasonal field release and spermatization trials of transgenic hypovirulent strains of Cryphonectria parasitica containing cDNA copies of hypovirus CHV1-EP713 | FOREST PATHOLOGY |
| Roques, A; Zhao, LL; Sun, JH; Robinet, C | 2015 | Pine Wood Nematode, Pine Wilt Disease, Vector Beetle and Pine Tree: How a Multiplayer System Could Reply to Climate Change | CLIMATE CHANGE AND INSECT PESTS |
| Roser, D; Pasanen, K; Asikainen, A | 2006 | Decision-support program "EnerTree" for analyzing forest residue recovery options | BIOMASS & BIOENERGY |
| Rossi, JP; Samalens, JC; Guyon, D; van Halder, I; Jactel, H; Menassieu, P; Piou, D | 2009 | Multiscale spatial variation of the bark beetle Ips sexdentatus damage in a pine plantation forest (Landes de Gascogne, Southwestern France) | FOREST ECOLOGY AND MANAGEMENT |
| Rouault, G; Candau, JN; Lieutier, F; Nageleisen, LM; Martin, JC; Warzee, N | 2006 | Effects of drought and heat on forest insect populations in relation to the 2003 drought in Western Europe | ANNALS OF FOREST SCIENCE |
| Roversi, PF | 2008 | Aerial spraying of Bacillus thuringiensis var. kurstaki for the control of | PHYTOPARASITICA |

| | | | |
|---|---|---|---|
| | | Thaumetopoea processionea in Turkey oak woods | |
| Roversi, PF; Sciarretta, A; Marziali, L; Marianelli, L; Bagnoli, M | 2013 | A GIS-BASED COST DISTANCE APPROACH TO ANALYSE THE SPREAD OF MATSUCOCCUS FEYTAUDI IN TUSCANY, ITALY (COCCOIDEA MATSUCOCCIDAE) | REDIA-GIORNALE DI ZOOLOGIA |
| Rullan-Silva, C; Olthoff, AE; Pando, V; Pajares, JA; Delgado, JA | 2015 | Remote monitoring of defoliation by the beech leaf-mining weevil Rhynchaenus fagi in northern Spain | FOREST ECOLOGY AND MANAGEMENT |
| Rullan-Silva, CD; Olthoff, AE; de la Mata, JAD; Pajares-Alonso, JA | 2013 | Remote monitoring of forest insect defoliation. A review | FOREST SYSTEMS |
| Rumbou, A; von Bargen, S; Buttner, C | 2009 | A model system for plant-virus interaction-infectivity and seed transmission of Cherry leaf roll virus (CLRV) in Arabidopsis thaliana | EUROPEAN JOURNAL OF PLANT PATHOLOGY |
| Ruohomaki, K; Tanhuanpaa, M; Ayres, MP; Kaitaniemi, P; Tammaru, T; Haukioja, E | 2000 | Causes of cyclicity of Epirrita autumnata (Lepidoptera, Geometridae): grandiose theory and tedious practice | POPULATION ECOLOGY |
| Salman, MHR; Hellrigl, K; Minerbi, S; Battisti, A | 2016 | Prolonged pupal diapause drives population dynamics of the pine processionary moth (Thaumetopoea pityocampa) in an outbreak expansion area | FOREST ECOLOGY AND MANAGEMENT |
| Samojlik, T; Selva, N; Daszkiewicz, P; Fedotova, A; Wajrak, A; Kuijper, DPJ | 2018 | Lessons from Bialowieza Forest on the history of protection and the world's first reintroduction of a large carnivore | CONSERVATION BIOLOGY |
| Sanguesa-Barreda, G; Camarero, JJ; Garcia-Martin, A; Hernandez, R; de la Riva, J | 2014 | Remote-sensing and tree-ring based characterization of forest defoliation and growth loss due to the Mediterranean pine processionary moth | FOREST ECOLOGY AND MANAGEMENT |

| | | | |
|---|---|---|---|
| Sarikaya, O; Catal, Y | 2014 | Effects of Dioryctria sylvestrella (Ratzeburg, 1840) on Basal Area Increment Loss of the young Brutian pine (Pinus brutia Ten.) Trees in the south-western of Turkey | RESEARCH JOURNAL OF BIOTECHNOLOGY |
| Sarikaya, O; Karaceylan, IB; Sen, I | 2018 | MAXIMUM ENTROPY MODELING (MAXENT) OF CURRENT AND FUTURE DISTRIBUTIONS OF IPS MANNSFELDI (WACHTL, 1879) (CURCULIONIDAE: SCOLYTINAE) IN TURKEY | APPLIED ECOLOGY AND ENVIRONMENTAL RESEARCH |
| Scala, E; Micheli, M; Ferretti, F; Maresi, G; Zottele, F; Piskur, B; Scattolin, L | 2019 | New diseases due to indigenous fungi in a changing world: The case of hop hornbeam canker in the Italian Alps | FOREST ECOLOGY AND MANAGEMENT |
| Schiegg, K | 2001 | Saproxylic insect diversity of beech: limbs are richer than trunks | FOREST ECOLOGY AND MANAGEMENT |
| Schmalholz, M; Hylander, K; Frego, K | 2011 | Bryophyte species richness and composition in young forests regenerated after clear-cut logging versus after wildfire and spruce budworm outbreak | BIODIVERSITY AND CONSERVATION |
| Schnee, H | 1999 | The American spruce needle-miner (Coleotechnites piceaella (KEARFOTT) (Lepidoptera, Gelechiidae) in Saxonia - distribution, bionomy, parasitoids. | COMMUNICATIONS OF THE GERMAN SOCIETY FOR GENERAL AND APPLIED ENTOMOLOGY, VOL 12, NOS 1-6, FEB 2000 |
| Schoebel, CN; Jung, E; Prospero, S | 2013 | Development of New Polymorphic Microsatellite Markers for Three Closely Related Plant-Pathogenic Phytophthora Species Using 454-Pyrosequencing and Their Potential Applications | PHYTOPATHOLOGY |
| Schott, T; Kapari, L; Hagen, SB; Vindstad, OPL; Jepsen, JU; Ims, RA | 2013 | Predator release from invertebrate generalists does not explain geometrid moth | CANADIAN ENTOMOLOGIST |

| | | | |
|---|---|---|---|
| | | (Lepidoptera: Geometridae) outbreaks at high altitudes | |
| Schroeder, H; Degen, B | 2008 | Spatial genetic structure in populations of the green oak leaf roller, Tortrix viridana L. (Lepidoptera, Tortricidae) | EUROPEAN JOURNAL OF FOREST RESEARCH |
| Schroeder, LM | 2007 | Escape in space from enemies: a comparison between stands with and without enhanced densities of the spruce bark beetle | AGRICULTURAL AND FOREST ENTOMOLOGY |
| Schulze, ED; Aas, G; Grimm, GW; Gossner, MM; Walentowski, H; Ammer, C; Kuhn, I; Bouriaud, O; von Gadow, K | 2016 | A review on plant diversity and forest management of European beech forests | EUROPEAN JOURNAL OF FOREST RESEARCH |
| Schumacher, J; Kehr, R; Leonhard, S | 2010 | Mycological and histological investigations of Fraxinus excelsior nursery saplings naturally infected by Chalara fraxinea | FOREST PATHOLOGY |
| Schumacher, VJ; Solger, A; Leonhard, S; Roloff, A | 2003 | Increasing occurrence of stem and sawn wood blue-stain in the tree species Norway spruce (Picea abies [L.] KARST.) in Saxony | ALLGEMEINE FORST UND JAGDZEITUNG |
| Seidl, R; Blennow, K | 2012 | Pervasive Growth Reduction in Norway Spruce Forests following Wind Disturbance | PLOS ONE |
| Seidl, R; Schelhaas, MJ; Lexer, MJ | 2011 | Unraveling the drivers of intensifying forest disturbance regimes in Europe | GLOBAL CHANGE BIOLOGY |
| Seifert, T | 2007 | Simulating the extent of decay caused by Heterobasidion annosum s. l. in stems of Norway spruce | FOREST ECOLOGY AND MANAGEMENT |
| Selas, V; Hogstad, A; Kobro, S; Rafoss, T | 2004 | Can sunspot activity and ultraviolet-B radiation explain cyclic outbreaks of forest moth pest species? | PROCEEDINGS OF THE ROYAL SOCIETY B-BIOLOGICAL SCIENCES |
| Shinya, R; Morisaka, H; Kikuchi, T; Takeuchi, Y; Ueda, M; Futai, K | 2013 | Secretome Analysis of the Pine Wood Nematode | PLOS ONE |

| | | | |
|---|---|---|---|
| | | Bursaphelenchus xylophilus Reveals the Tangled Roots of Parasitism and Its Potential for Molecular Mimicry | |
| Shorohova, E; Kapitsa, E; Kazartsev, I; Romashkin, I; Polevoi, A; Kushnevskaya, H | 2016 | Tree species traits are the predominant control on the decomposition rate of tree log bark in a mesic old-growth boreal forest | FOREST ECOLOGY AND MANAGEMENT |
| Shorohova, E; Kneeshaw, D; Kuuluvainen, T; Gauthier, S | 2011 | Variability and Dynamics of Old-Growth Forests in the Circumboreal Zone: Implications for Conservation, Restoration and Management | SILVA FENNICA |
| Shorohova, E; Kuuluvainen, T; Kangur, A; Jogiste, K | 2009 | Natural stand structures, disturbance regimes and successional dynamics in the Eurasian boreal forests: a review with special reference to Russian studies | ANNALS OF FOREST SCIENCE |
| Short, I; Hawe, J | 2018 | Living with ash dieback - Silviculture systems for Irish ash | 8TH HARDWOOD CONFERENCE WITH SPECIAL FOCUS ON NEW ASPECTS ON HARDWOOD UTILIZATION - FROM SCIENCE TO TECHNOLOGY |
| Sierota, Z; Grodzki, W; Szczepkowski, A | 2019 | Abiotic and Biotic Disturbances Affecting Forest Health in Poland over the past 30 Years: Impacts of Climate and Forest Management | FORESTS |
| Sierpinska, A | 1998 | Towards an integrated management of Dendrolimus pini L. | POPULATION DYNAMICS, IMPACTS, AND INTEGRATED MANAGEMENT OF FOREST DEFOLIATING INSECTS - PROCEEDINGS |
| Silingiene, G; Vasinauskiene, R; Brazaitis, G | 2013 | Damp Water Steam Impact on Fungi on Norway Spruce (Picea abies (L.) Karst.) Seeds | RURAL DEVELOPMENT 2013: PROCEEDINGS VOL 6, BOOK 3 |
| Sirocko, F; Knapp, H; Dreher, F; Forster, MW; Albert, J; Brunck, H; Veres, D; Dietrich, S; Zech, M; | 2016 | The ELSA-Vegetation-Stack: Reconstruction of Landscape Evolution Zones (LEZ) from | GLOBAL AND PLANETARY CHANGE |

| | | | |
|---|---|---|---|
| Hambach, U; Rohner, M; Rudert, S; Schwibus, K; Adams, C; Sigl, P | | laminated Eifel maar sediments of the last 60,000 years | |
| Sjoman, H; Ostberg, J | 2019 | Vulnerability of ten major Nordic cities to potential tree losses caused by longhorned beetles | URBAN ECOSYSTEMS |
| Slawski, M; Mokrzycki, T; Perlinski, S; Rutkiewicz, A; Slawska, M | 2019 | Distribution of wintering pre-imaginal stages of the great web-spinning pine sawfly Acantholyda posticalis Mats. in Scots pine stands being the outbreak centres | SYLWAN |
| Slowinski, M; Lamentowicz, M; Lucow, D; Barabach, J; Brykala, D; Tyszkowski, S; Pienczewska, A; Snieszko, Z; Dietze, E; Jazdzewski, K; Obremska, M; Ott, F; Brauer, A; Marcisz, K | 2019 | Paleoecological and historical data as an important tool in ecosystem management | JOURNAL OF ENVIRONMENTAL MANAGEMENT |
| Smits, A | 2002 | Performance of pine looper Bupalus piniarius larvae under population build-up conditions | ENTOMOLOGIA EXPERIMENTALIS ET APPLICATA |
| Solis-Hernandez, A; Rodriguez-Vivas, R; Esteve-Gasent, MD; Villegas-Perez, SL | 2018 | Detection of Borrelia burgdorferi sensu lato in dogs and its ticks in rural communities of Yucatan, Mexico | REVISTA DE BIOLOGIA TROPICAL |
| Sorvari, J | 2009 | Foraging distances and potentiality in forest pest insect control: an example with two candidate ants (Hymenoptera: Formicidae) | MYRMECOLOGICAL NEWS |
| Sousa-Silva, R; Verbist, B; Lomba, A; Valent, P; Suskevics, M; Picard, O; Hoogstra-Klein, MA; Cosofret, VC; Bouriaud, L; Ponette, Q; Verheyen, K; Muys, B | 2018 | Adapting forest management to climate change in Europe: Linking perceptions to adaptive responses | FOREST POLICY AND ECONOMICS |
| Speer, JH; Kulakowski, D | 2017 | Creating a Buzz: Insect Outbreaks and Disturbance Interactions | DENDROECOLOGY: TREE-RING ANALYSES APPLIED TO ECOLOGICAL STUDIES |
| Stadelmann, G; Bugmann, H; Meier, F; Wermelinger, B; Bigler, C | 2013 | Effects of salvage logging and sanitation felling on bark beetle (Ips typographus L.) infestations | FOREST ECOLOGY AND MANAGEMENT |
| Stadelmann, G; Bugmann, H; Wermelinger, B; Bigler, C | 2014 | Spatial interactions between storm damage and subsequent | FOREST ECOLOGY AND MANAGEMENT |

| | | | |
|---|---|---|---|
| | | infestations by the European spruce bark beetle | |
| Stadler, B; Michalzik, B; Muller, T | 1998 | Linking aphid ecology with nutrient fluxes in a coniferous forest | ECOLOGY |
| Stadler, B; Solinger, S; Michalzik, B | 2001 | Insect herbivores and the nutrient flow from the canopy to the soil in coniferous and deciduous forests | OECOLOGIA |
| Stalb, S; Polley, B; Danner, KJ; Reule, M; Tomaso, H; Hackbare, A; Wagner-Wiening, C; Sting, R | 2017 | Detection of tularemia in European brown hares (Lepus europaeus) and humans reveals endemic and seasonal occurrence in Baden-Wuerttemberg, Germany | BERLINER UND MUNCHENER TIERARZTLICHE WOCHENSCHRIFT |
| Steinbauer, MJ; Mc Quillan, PB; Young, CJ | 2001 | Life history and behavioural traits of Mnesampela privata that exacerbate population responses to eucalypt plantations: Comparisons with Australian and outbreak species of forest geometrid from the Northern Hemisphere | AUSTRAL ECOLOGY |
| Stivrins, N; Buchan, MS; Disbrey, HR; Kuosmanen, N; Latalowa, M; Lempinen, J; Muukkonen, P; Slowinski, M; Veski, S; Seppa, H | 2017 | Widespread, episodic decline of alder (Alnus) during the medieval period in the boreal forest of Europe | JOURNAL OF QUATERNARY SCIENCE |
| Stojanovic, D; Curcic, S; Orlovic, S; Galic, Z | 2011 | NOCTUID PEST SPECIES INVENTORY (LEPIDOPTERA: NOCTUIDAE) OF THE NATIONAL PARK "FRUSKA GORA" | SUMARSKI LIST |
| Stojanovic, DB; Levanic, T; Matovic, B; Stjepanovic, S; Orlovic, S | 2018 | Growth response of different tree species (oaks, beech and pine) from SE Europe to precipitation over time | DENDROBIOLOGY |
| Straw, NA; Williams, DT; Kulinich, O; Gninenko, YI | 2013 | Distribution, impact and rate of spread of emerald ash borer Agrilus planipennis (Coleoptera: Buprestidae) in the Moscow region of Russia | FORESTRY |

| | | | |
|---|---|---|---|
| Stursova, M; Snajdr, J; Cajthaml, T; Barta, J; Santruckova, H; Baldrian, P | 2014 | When the forest dies: the response of forest soil fungi to a bark beetle-induced tree dieback | ISME JOURNAL |
| Su, Y; Langhammer, J; Jarsjo, J | 2017 | Geochemical responses of forested catchments to bark beetle infestation: Evidence from high frequency in-stream electrical conductivity monitoring | JOURNAL OF HYDROLOGY |
| Sukovata, L; Kolk, A | 2007 | The use of semiochemicals in forest protection - New challenges | QUO VADIS, FORESTRY?, PROCEEDINGS |
| Sun, D; Walsh, D | 1998 | Review of studies on environmental impacts of recreation and tourism in Australia | JOURNAL OF ENVIRONMENTAL MANAGEMENT |
| Szekeres, S; Lugner, J; Fingerle, V; Margos, G; Foldvari, G | 2017 | Prevalence of Borrelia miyamotoi and Borrelia burgdorferi sensu lato in questing ticks from a recreational coniferous forest of East Saxony, Germany | TICKS AND TICK-BORNE DISEASES |
| Tabakovic-Tosic, M | 2014 | SUPPRESSION OF GYPSY MOTH POPULATION IN MOUNTAIN AVALA (REPUBLIC OF SERBIA) BY INTRODUCTION OF ENTOMOPATHOGENIC FUNGUS ENTOMOPHAGA MAIMAIGA | COMPTES RENDUS DE L ACADEMIE BULGARE DES SCIENCES |
| Tabakovic-Tosic, M; Georgiev, G; Mirchev, P; Tosic, D; Curguz, VG | 2013 | Gypsy Moth in Central Serbia Over the Previous Fifty Years | ACTA ZOOLOGICA BULGARICA |
| Tabakovic-Tosic, M; Tosic, D; Rajkovic, S; Golubovic-Curguz, V; Rakonjac, L | 2011 | Invasion species Coleophora laricella - One of the main limiting factor of Larix decidua during the forest aforestation and recultivation | AFRICAN JOURNAL OF AGRICULTURAL RESEARCH |
| Tammaru, T; Kaitaniemi, P; Ruohomaki, K | 1995 | Oviposition choices of Epirrita autumnata (Lepidoptera: Geometridae) in relation to its eruptive population dynamics | OIKOS |

| | | | |
|---|---|---|---|
| Tanney, JB; McMullin, DR; Miller, JD | 2018 | Toxigenic Foliar Endophytes from the Acadian Forest | ENDOPHYTES OF FOREST TREES: BIOLOGY AND APPLICATIONS, 2ND EDITION |
| Tenow, O; Bylund, H | 2000 | Recovery of a Betula pubescens forest in northern Sweden after severe defoliation by Epirrita autumnata | JOURNAL OF VEGETATION SCIENCE |
| Tenow, O; Nilssen, AC; Bylund, H; Hogstad, O | 2007 | Waves and synchrony in Epirrita autumnata/Operophtera brumata outbreaks. I. Lagged synchrony: regionally, locally and among species | JOURNAL OF ANIMAL ECOLOGY |
| Tenow, O; Nilssen, AC; Bylund, H; Pettersson, R; Battisti, A; Bohn, U; Caroulle, F; Ciornei, C; Csoka, G; Delb, H; De Prins, W; Glavendekic, M; Gninenko, YI; Hrasovec, B; Matosevic, D; Meshkova, V; Moraal, L; Netoiu, C; Pajares, J; Rubtsov, V; Tomescu, R; Utkina, I | 2013 | Geometrid outbreak waves travel across Europe | JOURNAL OF ANIMAL ECOLOGY |
| Tenow, O; Nilssen, AC; Holmgren, B; Elverum, F | 1999 | An insect (Argyresthia retinella, Lep., Yponomeutidae) outbreak in northern birch forests, released by climatic changes? | JOURNAL OF APPLIED ECOLOGY |
| Terhonen, E; Langer, GJ; Busskamp, J; Rascutoi, DR; Blumenstein, K | 2019 | Low Water Availability Increases Necrosis in Picea abies after Artificial Inoculation with Fungal Root Rot Pathogens Heterobasidion parviporum and Heterobasidion annosum | FORESTS |
| Thomas, P; Buntgen, U | 2019 | A risk assessment of Europe's black truffle sector under predicted climate change | SCIENCE OF THE TOTAL ENVIRONMENT |
| Thor, M; Arlinger, JD; Stenlid, J | 2006 | Heterobasidion annosum root rot in Picea abies: Modelling economic outcomes of stump treatment in Scandinavian coniferous forests | SCANDINAVIAN JOURNAL OF FOREST RESEARCH |
| Thorn, S; Werner, SAB; Wohlfahrt, J; Bassler, C; Seibold, S; Quillfeldt, P; Muller, J | 2016 | Response of bird assemblages to windstorm and salvage | ECOLOGICAL INDICATORS |

| | | | |
|---|---|---|---|
| | | logging - Insights from analyses of functional guild and indicator species | |
| Timmermann, V; Nagy, NE; Hietala, AM; Borja, I; Solheim, H | 2017 | Progression of Ash Dieback in Norway Related to Tree Age, Disease History and Regional Aspects | BALTIC FORESTRY |
| Toigo, M; Barraquand, F; Barnagaud, JY; Piou, D; Jactel, H | 2017 | Geographical variation in climatic drivers of the pine processionary moth population dynamics | FOREST ECOLOGY AND MANAGEMENT |
| Toivanen, T; Liikanen, V; Kotiaho, JS | 2009 | Effects of forest restoration treatments on the abundance of bark beetles in Norway spruce forests of southern Finland | FOREST ECOLOGY AND MANAGEMENT |
| Tomlinson, I; Potter, C; Bayliss, H | 2015 | Managing tree pests and diseases in urban settings: The case of Oak Processionary Moth in London, 2006-2012 | URBAN FORESTRY & URBAN GREENING |
| Tomlinson, JA; Boonham, N; Hughes, KJD; Griffen, RL; Barker, I | 2005 | On-site DNA extraction and real-time PCR for detection of Phytophthora ramorum in the field | APPLIED AND ENVIRONMENTAL MICROBIOLOGY |
| Trujillo-Toro, J; Navarro-Cerrillo, RM | 2019 | Analysis of Site-dependent Pinus halepensis Mill. Defoliation Caused by "Candidatus Phytoplasma pini' through Shape Selection in Landsat Time Series | REMOTE SENSING |
| Tsouvalis, J | 2019 | The post-politics of plant biosecurity: The British Government's response to ash dieback in 2012 | TRANSACTIONS OF THE INSTITUTE OF BRITISH GEOGRAPHERS |
| Tudoran, MM; Marquer, L; Jonsson, AM | 2016 | Historical experience (1850-1950 and 1961-2014) of insect species responsible for forest damage in Sweden: Influence of climate and land management changes | FOREST ECOLOGY AND MANAGEMENT |
| Tulik, M; Zakrzewski, J; Adamczyk, J; Tereba, A; Yaman, B; Nowakowska, JA | 2017 | Anatomical and genetic aspects of ash dieback: a | IFOREST-BIOGEOSCIENCES AND FORESTRY |

| | | | |
|---|---|---|---|
| | | look at the wood structure | |
| Turczanski, K | 2016 | Occurrence and spread of Chalara fraxinea on common ash (Fraxinus excelsior L.) in the selected countries of Northern Europe | SYLWAN |
| Turk, N; Margaletic, J; Markotic, A | 2009 | FOREST ECOSYSTEMS AND ZOONOSES | WILDLIFE: DESTRUCTION, CONSERVATION AND BIODIVERSITY |
| Ulyshen, MD | 2013 | Strengthening the case for saproxylic arthropod conservation: a call for ecosystem services research | INSECT CONSERVATION AND DIVERSITY |
| Vacek, S; Hejcmanova, P; Hejcman, M; Vacek, Z | 2013 | Growth, healthy status and seed production of differently aged allochtonous and autochtonous Pinus mugo stands in the Giant Mts. over 30 years | EUROPEAN JOURNAL OF FOREST RESEARCH |
| Vainio, EJ; Jurvansuu, J; Hyder, R; Kashif, M; Piri, T; Tuomivirta, T; Poimala, A; Xu, P; Makela, S; Nitisa, D; Hantula, J | 2018 | Heterobasidion Partitivirus 13 Mediates Severe Growth Debilitation and Major Alterations in the Gene Expression of a Fungal Forest Pathogen | JOURNAL OF VIROLOGY |
| Valadas, V; Laranjo, M; Barbosa, P; Espada, M; Mota, M; Oliveira, S | 2012 | The pine wood nematode, Bursaphelenchus xylophilus, in Portugal: possible introductions and spread routes of a serious biological invasion revealed by molecular methods | NEMATOLOGY |
| Valadas, V; Laranjo, M; Mota, M; Oliveira, S | 2013 | Molecular characterization of Portuguese populations of the pinewood nematode Bursaphelenchus xylophilus using cytochrome b and cellulase genes | JOURNAL OF HELMINTHOLOGY |
| Valenta, V; Moser, D; Kapeller, S; Essl, F | 2017 | A new forest pest in Europe: a review of Emerald ash borer | JOURNAL OF APPLIED ENTOMOLOGY |

| | | | |
|---|---|---|---|
| | | (Agrilus planipennis) invasion | |
| Valkama, H; Martikainen, P; Raty, M | 1997 | First record of North American ambrosia beetle Gnathotrichus materiarius (Fitch) (Coleoptera, scolytidae) in Finland - a new potential forest pest? | ENTOMOLOGICA FENNICA |
| Van Der Nest, A; Wingfield, MJ; Janousek, J; Barnes, I | 2019 | Lecanosticta acicola: A growing threat to expanding global pine forests and plantations | MOLECULAR PLANT PATHOLOGY |
| van Lierop, P; Lindquist, E; Sathyapala, S; Franceschini, G | 2015 | Global forest area disturbance from fire, insect pests, diseases and severe weather events | FOREST ECOLOGY AND MANAGEMENT |
| Vannini, A; Natili, G; Anselmi, N; Montaghi, A; Vettraino, AM | 2010 | Distribution and gradient analysis of Ink disease in chestnut forests | FOREST PATHOLOGY |
| Vasaitis, R; Burnevica, N; Uotila, A; Dahlberg, A; Kasanen, R | 2016 | Cut Picea abies Stumps Constitute Low Quality Substrate for Sustaining Biodiversity in Fungal Communities | BALTIC FORESTRY |
| Vindstad, OPL; Jepsen, JU; Ek, M; Pepi, A; Ims, RA | 2019 | Can novel pest outbreaks drive ecosystem transitions in northern-boreal birch forest? | JOURNAL OF ECOLOGY |
| Vindstad, OPL; Jepsen, JU; Klinghardt, M; Ek, M; Ims, RA | 2017 | Salvage logging of mountain birch after geometrid outbreaks: Ecological context determines management outcomes | FOREST ECOLOGY AND MANAGEMENT |
| Vindstad, OPL; Jepsen, JU; Yoccoz, NG; Bjornstad, ON; Mesquita, MDS; Ims, RA | 2019 | Spatial synchrony in sub-arctic geometrid moth outbreaks reflects dispersal in larval and adult life cycle stages | JOURNAL OF ANIMAL ECOLOGY |
| Vindstad, OPL; Schultze, S; Jepsen, JU; Biuw, M; Kapari, L; Sverdrup-Thygeson, A; Ims, RA | 2014 | Numerical Responses of Saproxylic Beetles to Rapid Increases in Dead Wood Availability following Geometrid Moth Outbreaks in Sub-Arctic Mountain Birch Forest | PLOS ONE |
| Virtanen, T; Neuvonen, S | 1999 | Performance of moth larvae on birch in relation | OECOLOGIA |

| | | | |
|---|---|---|---|
| | | to altitude, climate, host quality and parasitoids | |
| Virtanen, T; Neuvonen, S; Nikula, A | 1998 | Modelling topoclimatic patterns of egg mortality of Epirrita autumnata (Lepidoptera : Geometridae) with a Geographical Information System: predictions for current climate and warmer climate scenarios | JOURNAL OF APPLIED ECOLOGY |
| Voolma, K; Hiiesaar, K; Williams, IH; Ploomi, A; Jogar, K | 2016 | Cold hardiness in the pre-imaginal stages of the great web-spinning pine-sawfly Acantholyda posticalis | AGRICULTURAL AND FOREST ENTOMOLOGY |
| Waller, M | 2013 | Drought, disease, defoliation and death: forest pathogens as agents of past vegetation change | JOURNAL OF QUATERNARY SCIENCE |
| Wang, P; Chen, GF; Zhang, JS; Xue, Q; Zhang, JH; Chen, C; Zhang, QH | 2017 | Pheromone-trapping the nun moth, Lymantria monacha (Lepidoptera: Lymantriidae) in Inner Mongolia, China | INSECT SCIENCE |
| Wang, XM; Stenstrom, E; Boberg, J; Ols, C; Drobyshev, I | 2017 | Outbreaks of Gremmeniella abietina cause considerable decline in stem growth of surviving Scots pine trees | DENDROCHRONOLOGIA |
| Watt, AD; Hicks, BJ | 2000 | A reappraisal of the population dynamics of the pine beauty moth, Panolis flammea, on lodgepole pine, Pinus contorta, in Scotland | POPULATION ECOLOGY |
| WATT, AD; LEATHER, SR; FORREST, GI | 1991 | THE EFFECT OF PREVIOUS DEFOLIATION OF POLE-STAGE LODGEPOLE PINE ON PLANT CHEMISTRY, AND ON THE GROWTH AND SURVIVAL OF PINE BEAUTY MOTH (PANOLIS-FLAMMEA) LARVAE | OECOLOGIA |
| Wermelinger, B | 2004 | Ecology and management of the spruce bark beetle Ips typographus - a review of recent research | FOREST ECOLOGY AND MANAGEMENT |

| | | | |
|---|---|---|---|
| Werner, SM; Albers, MA; Cryderman, T; Diminic, D; Heyd, R; Hrasovic, B; Kobro, S; Larsson, S; Mech, R; Niemela, P; Rousi, M; Raffa, KF; Scanlon, K; Weber, S | 2006 | Is the outbreak status of Thrips calcaratus Uzel in North America due to altered host relationships? | FOREST ECOLOGY AND MANAGEMENT |
| Wesolowski, T; Rowinski, P | 2008 | Late leaf development in pedunculate oak (Quercus robur): An antiherbivore defence? | SCANDINAVIAN JOURNAL OF FOREST RESEARCH |
| Wesolowski, T; Rowinski, P | 2006 | Tree defoliation by winter moth Operophtera brumata L. during an outbreak affected by structure of forest landscape | FOREST ECOLOGY AND MANAGEMENT |
| Widmer, TL | 2015 | Differences in Virulence and Sporulation of Phytophthora kemoviae Isolates Originating From Two Distinct Geographical Regions | PLANT DISEASE |
| Wielgolaski, FE; Hofgaard, A; Holtmeier, FK | 2017 | Sensitivity to environmental change of the treeline ecotone and its associated biodiversity in European mountains | CLIMATE RESEARCH |
| Williams, DT; Straw, NA; Day, KR | 2003 | Defoliation of Sitka spruce by the European spruce sawfly, Gilpinia hercyniae (Hartig): a retrospective analysis using the needle trace method | AGRICULTURAL AND FOREST ENTOMOLOGY |
| Winde, I; Anderbrant, O; Jonsson, AM | 2018 | Tree recovery during the aftermath of an outbreak episode of the Hungarian spruce scale in southern Sweden | SCANDINAVIAN JOURNAL OF FOREST RESEARCH |
| Winter, MB; Ammer, C; Baier, R; Donato, DC; Seibold, S; Muller, J | 2015 | Multi-taxon alpha diversity following bark beetle disturbance: Evaluating multi-decade persistence of a diverse early-seral phase | FOREST ECOLOGY AND MANAGEMENT |
| Wolf, A; Kozlov, MV; Callaghan, TV | 2008 | Impact of non-outbreak insect damage on vegetation in northern Europe will be greater than expected during a changing climate | CLIMATIC CHANGE |

| | | | |
|---|---|---|---|
| Wu, N; Abril, C; Thomann, A; Grosclaude, E; Doherr, MG; Boujon, P; Ryser-Degiorgis, MP | 2012 | Risk factors for contacts between wild boar and outdoor pigs in Switzerland and investigations on potential Brucella suis spill-over | BMC VETERINARY RESEARCH |
| Wyka, SA; McIntire, CD; Smith, C; Munck, IA; Rock, BN; Asbjornsen, H; Broders, KD | 2018 | Effect of Climatic Variables on Abundance and Dispersal of Lecanosticta acicola Spores and Their Impact on Defoliation on Eastern White Pine | PHYTOPATHOLOGY |
| Xie, SA; Lv, SJ | 2013 | Effect of different semiochemicals blends on spruce bark beetle, Ips typographus (Coleoptera: Scolytidae) | ENTOMOLOGICAL SCIENCE |
| Young, AB; Cairns, DM; Lafon, CW; Moen, J | 2014 | Geometrid moth outbreaks and their climatic relations in northern Sweden | ARCTIC ANTARCTIC AND ALPINE RESEARCH |
| Zaluma, A; Muiznieks, I; Gaitnieks, T; Burnevica, N; Jansons, A; Jansons, J; Stenlid, J; Vasaitis, R | 2019 | Infection and spread of root rot caused by Heterobasidion spp. in Pinus contorta plantations in Northern Europe: three case studies | CANADIAN JOURNAL OF FOREST RESEARCH |
| Zamora-Ballesteros, C; Haque, MMU; Diez, JJ; Martin-Garcia, J | 2017 | Pathogenicity of Phytophthora alni complex and P.plurivora in Alnus glutinosa seedlings | FOREST PATHOLOGY |
| Zang, C; Helm, R; Sparks, TH; Menzel, A | 2015 | Forecasting bark beetle early flight activity with plant phenology | CLIMATE RESEARCH |
| Zeng, QY; Hansson, P; Wang, XR | 2005 | Specific and sensitive detection of the conifer pathogen Gremmeniella abietina by nested PCR | BMC MICROBIOLOGY |
| Zmudzki, J; Jablonski, A; Nowak, A; Zebek, S; Arent, Z; Bocian, L; Pejsak, Z | 2016 | First overall report of Leptospira infections in wild boars in Poland | ACTA VETERINARIA SCANDINAVICA |
| Zottele, F; Salvadori, C; Corradini, S; Andreis, D; Wolynski, A; Maresi, G | 2014 | Chrysomyxa rhododendri in Trentino: a First Analysis of Monitoring Data | BALTIC FORESTRY |
| Zubrik, M; Hajek, A; Pilarska, D; Spilda, I; Georgiev, G; Hrasovec, B; Hirka, A; | 2016 | The potential for Entomophaga maimaiga | JOURNAL OF APPLIED ENTOMOLOGY |

| | | | |
|---|---|---|---|
| Goertz, D; Hoch, G; Barta, M; Saniga, M; Kunca, A; Nikolov, C; Vakula, J; Galko, J; Pilarski, P; Csoka, G | | to regulate gypsy moth Lymantria dispar (L.) (Lepidoptera: Erebidae) in Europe | |
| Zubrik, M; Spilda, I; Pilarska, D; Hajek, AE; Takov, D; Nikolov, C; Kunca, A; Pajtik, J; Lukasova, K; Holusa, J | 2018 | Distribution of the entomopathogenic fungus Entomophaga maimaiga (Entomophthorales: Entomophthoraceae) at the northern edge of its range in Europe | ANNALS OF APPLIED BIOLOGY |
