## Supplementary Material 4 for "Probability of outbreaks of forest insects in Europe: a generic model calibrated on six forest insect profiles"

Supplementary Material 4: Comparison of data and prediction for the six pest profiles. Empty points indicate year where no outbreak occurs, black points indicates years where outbreak occurs.

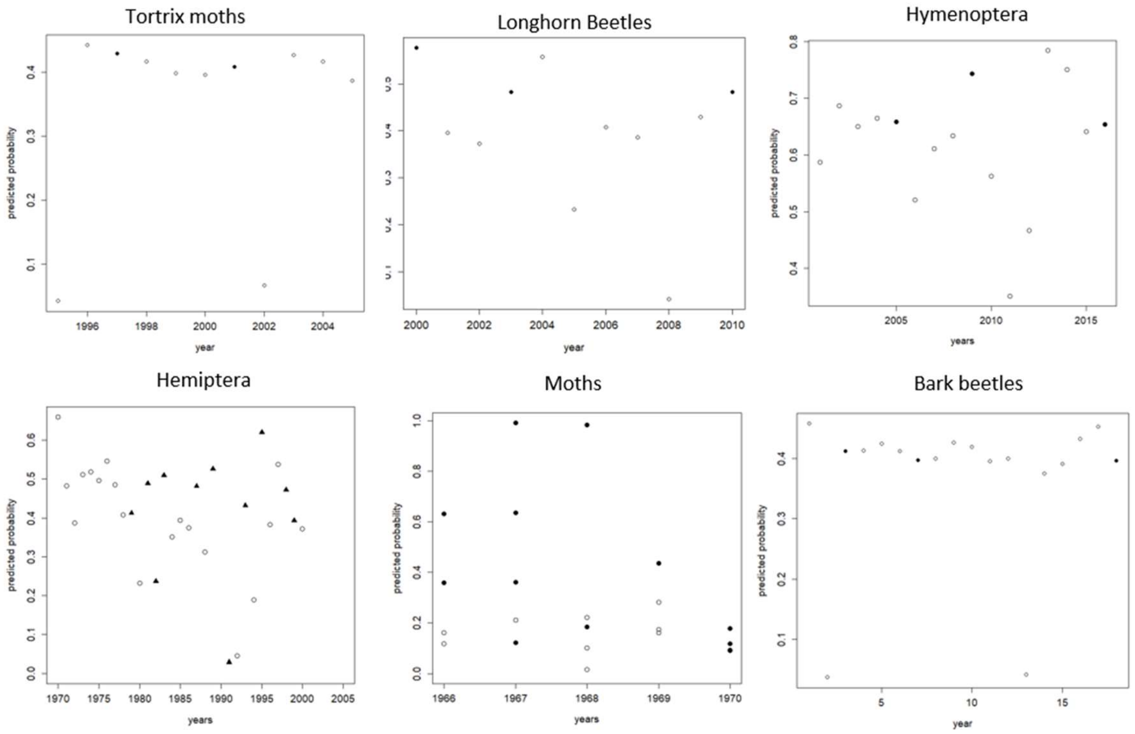
