## Supplementary Material 5 for "Probability of outbreaks of forest insects in Europe: a generic model calibrated on six forest insect profiles"

Supplementary Material 5: Outbreak probability for the Oak Processionary Moth for the North-Western part of France, from 1970 to 2018.

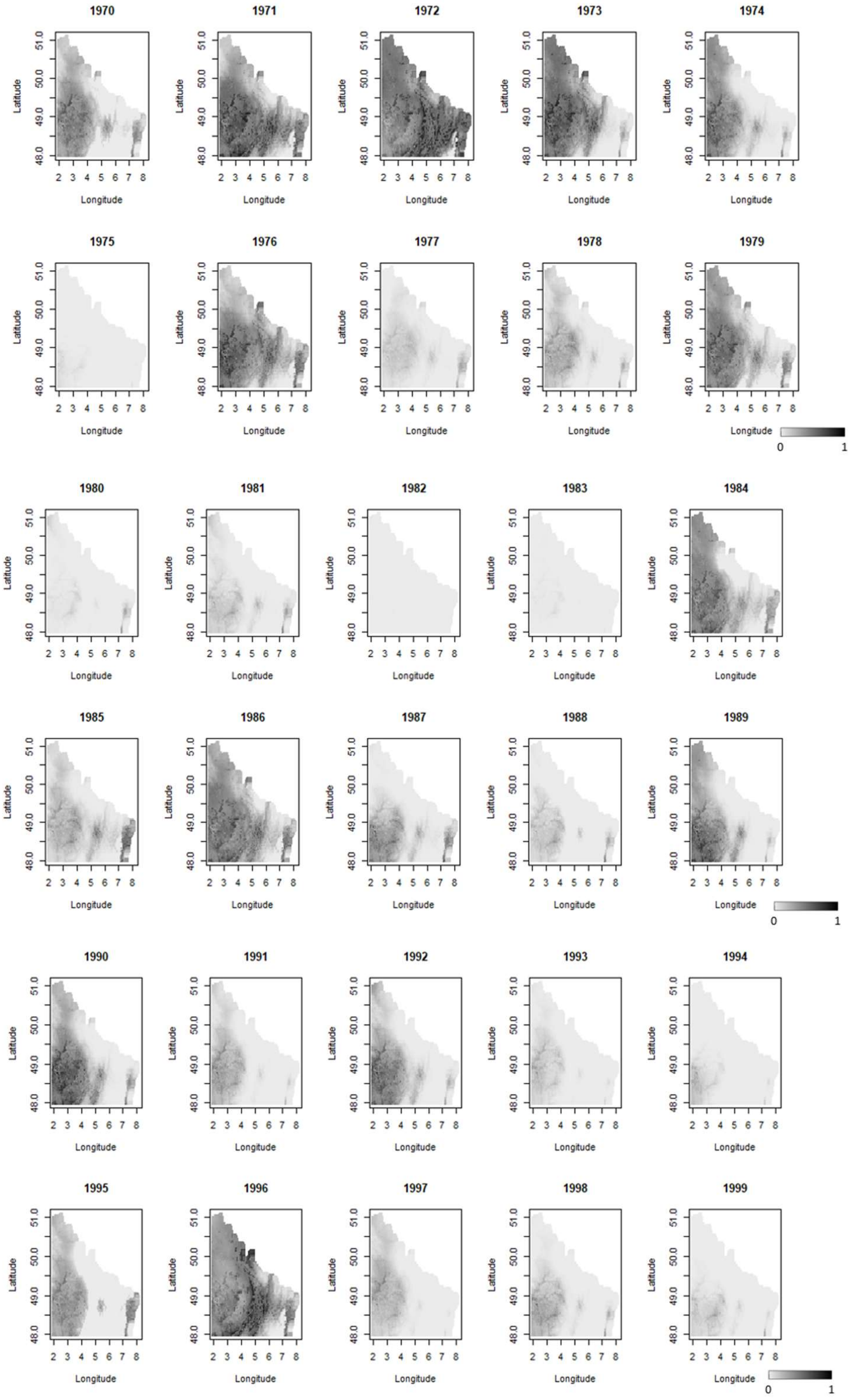

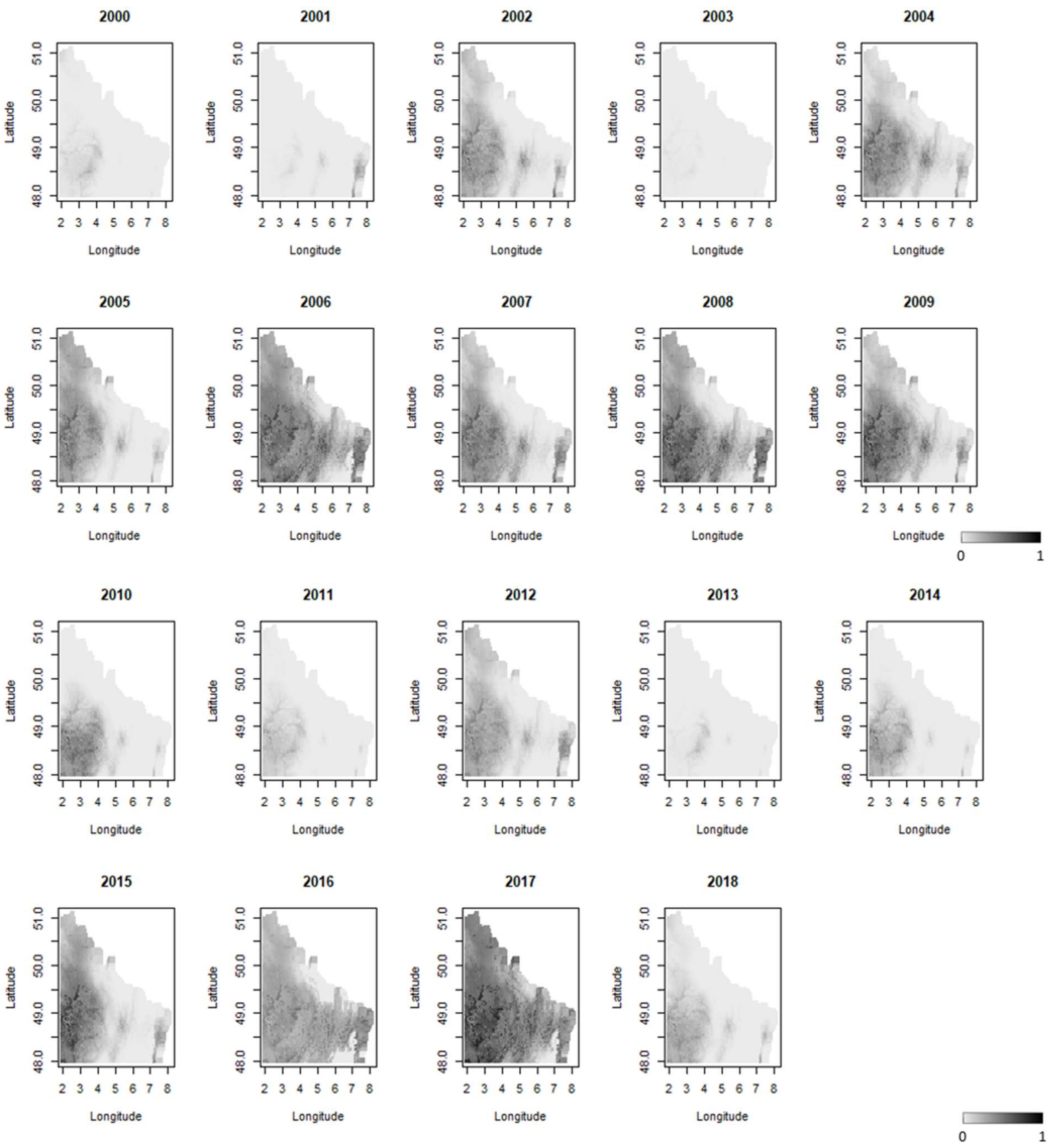
2000
2001
2002
2003
2004
2005
2006
2007
2008
2009
2010
2011
2012
2013
2014
2015
2016
2017
2018
Latitude
Longitude
51.0
50.0
49.0
48.0
2 3 4 5 6 7 8
0
1
